# Charting the development of *Drosophila* leg sensory organs at single-cell resolution

**DOI:** 10.1101/2022.10.07.511357

**Authors:** Ben R. Hopkins, Olga Barmina, Artyom Kopp

## Abstract

To respond to the world around them, animals rely on the input of a network of sensory organs distributed throughout the body. Distinct classes of sensory organ are specialized for the detection of specific stimuli such as strain, pressure, or taste. The features that underlie this specialization relate both to the neurons that innervate sensory organs and the accessory cells that comprise them. This diversity of cell types, both within and between sensory organs, raises two fundamental questions: what makes these cell types distinct from one another, and how is this diversity generated during development? To address these questions, we performed single-cell RNA sequencing on a developing tissue that displays a wide variety of functionally and structurally distinct sensory organs: the first tarsal segment of the pupal male *Drosophila melanogaster* foreleg. We characterize the cellular landscape in which the sensory organs reside, identify a novel cell type that contributes to the construction of the neural lamella, and characterize the transcriptomic differences among support cells within and between sensory organs. We identify the genes that distinguish between mechanosensory and chemosensory neurons, resolve a combinatorial transcription factor code that defines four distinct classes of gustatory neuron and several types of mechanosensory neuron, and match the expression of sensory receptors to specific neuron classes. Collectively, our work identifies core genetic features of a variety of sensory organs and provides a rich, annotated resource for studying their development and function.

## Introduction

All behaviour rests upon the ability of animals to detect variation in the internal and external environments. In multicellular animals, the detection of such variation is a function performed by sensory organs. With much of its external surface covered by many different classes of sensory organ, the fruit fly *Drosophila melanogaster* has long been used to investigate the mechanisms through which animals sense the world around them. Much attention has focused on the eyes, antennae, and maxillary palps, but the male *Drosophila* forelegs, which perform wide-ranging roles in locomotion, grooming, and courtship (e.g. Spieth, 1974; Bray & Amrein, 2003; Guo, Zhang, & Simpson, 2022), display a distinct repertoire of sensory organs (Figure 1A-B). Internal mechanosensory receptors known as chordotonal organs sense proprioceptive stimuli around leg joints (Mamiya *et al*., 2018) and substrate-borne vibrations (McKelvey *et al*., 2021)(Figure 1A). Campaniform sensilla, singly-innervated, shaftless sensors embedded in the cuticle, detect and relay cuticular strain, allowing for posture and intra-leg coordination to be maintained (Akay *et al*., 2004; Zill, Schmitz, & Büschges, 2004; Dinges *et al*., 2021)(Figure 1C). Mechanosensory bristles line the surface of the leg, detecting contact by deflection of an external, hair-like process (Hannah-Alava, 1958; Walker, Willingham, & Zuker, 2000) (Figure 1D). This organ class isn’t uniform: in the males of a subset of *Drosophila* species, including *D. melanogaster*, some mechanosensory bristles are heavily modified to generate a ‘sex comb’, an innovation critical for male mating success (Tokunaga, 1962; Ng & Kopp, 2008; Tanaka, Barmina, & Kopp, 2009; Massey *et al*., 2019)(Figure 1B). Finally, the foreleg also contains chemosensory taste bristles, which are sexually dimorphic in number and innervated by both multiple gustatory receptor neurons (GRNs) and a single mechanosensory neuron (Nayak & Singh, 1983; Stocker, 1994; Vosshall & Stocker, 2007)(Figure 1E). As in other parts of the body, such as the labellum (e.g. Montell, 2009), a degree of functional diversity exists between chemosensory taste bristles on the legs. The tunings and sensitivities of these bristles to a wide panel of tastants vary in relation to both the pair of legs on which they’re housed and their position within a given leg (Ling *et al*., 2014). This variation is, at least in part, achieved by restricting the expression of certain gustatory receptors to subsets of taste bristles (Bray & Amrein, 2003; Ling *et al*., 2014). A level below the bristles themselves, the multiple GRNs that innervate each bristle appear to perform distinct functions. Three distinct GRN classes involved in the detection and evaluation of conspecifics have been resolved in the leg, each of which makes a critical contribution to normal sexual behaviour (Thistle *et al*., 2012; Koh *et al*., 2014; Kallman, Kim, & Scott, 2015). Unravelling how this varied sensory apparatus is constructed through development and identifying the molecular basis of specialization in each sensory organ remain central objectives of developmental neurobiology.

**Figure 1.**
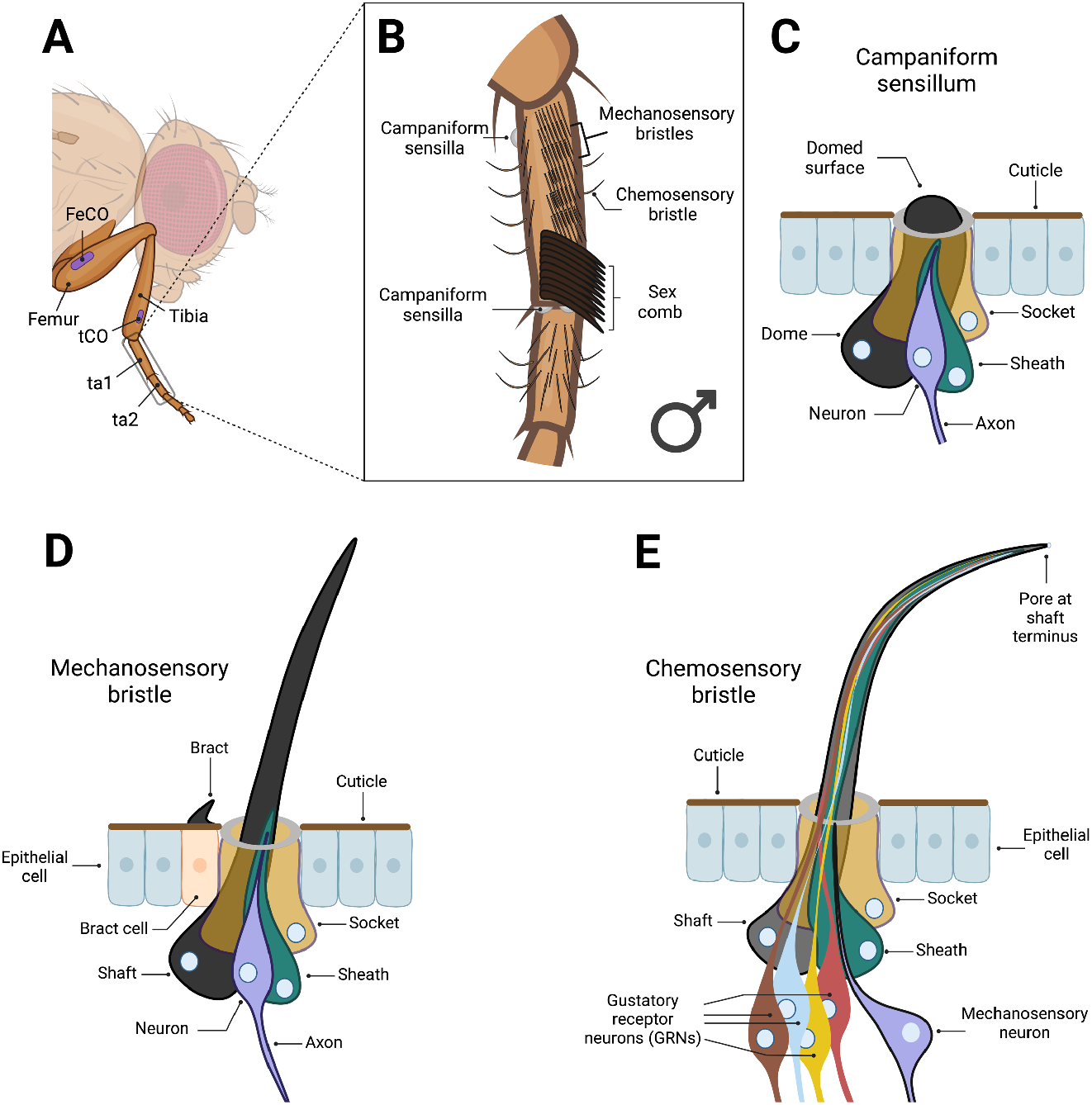
The first tarsal segment of the male *Drosophila melanogaster* foreleg carries multiple functionally and structurally distinct sensory organs. (A) Anatomy of the *Drosophila melanogaster* foreleg. The first tarsal segment (ta1), the focal region of this study, is distal to the tibia. Two chordotonal organs (CO) are present outside of the tarsal segments (approximate positions shown in purple). One is situated in the proximal femur (FeCO) and the other in the distal tibia (tCO) (Mamiya *et al*., 2018; McKelvey *et al*., 2021). (B) The ta1 of the *D. melanogaster* foreleg is enriched for a range of functionally and structurally diverse sensory organs. This region has the highest concentration of mechanosensory bristles of any part of the leg. Here, mechanosensory bristles are arranged in transverse rows on the ventral side, an arrangement thought to aid in grooming, and longitudinal rows on the anterior, dorsal, and posterior sides (Hannah-Alava, 1958). In males, the most distal transverse bristle row is transformed into the sex comb: the mechanosensory bristles, now ‘teeth’, are modified to be thicker, longer, blunter, and more heavily melanized, while the whole row is rotated 90° (Tokunaga, 1962; Tanaka *et al*., 2009). Males also show a sex-specific increase in the number of chemosensory taste bristles in ta1, bearing ~11 compared to the female’s ~7 (Nayak & Singh, 1983). Three campaniform sensilla are present in ta1, two on the dorsal, distal end of ta1 and one on the proximal, ventral side (Ta1GF and Ta1SF, respectively, using the nomenclature of Dinges *et al*., 2021); no campaniform sensilla are present in the distal tibia, ta2, or proximal ta3. (C-E) Campaniform sensilla, mechanosensory bristles, and chemosensory bristles are all composed of modified versions of four core cell-types: a socket (or ‘tormogen’), shaft/dome (or ‘trichogen’), sheath (or ‘thecogen’), and neuron (Keil, 1997a). The shaft and socket construct the external apparatus that provides the point of contact for mechanical or chemical stimuli and form a subcuticular lymph cavity that provides the ion source for the receptor current (Keil, 1997b; Chung *et al*., 2001). The sheath has glia-like properties, ensheathing the neuron and, as is thought, providing it with protection (Keil, 1997b). Ultimately, however, the contributions of these non-neuronal cells to sensory processing remain poorly characterized (Prelic *et al*., 2022). (C) Campaniform sensilla detect strain in the cuticle. They are singly innervated and capped with a dome, rather than a hair-like projection, which extends across the surface of the socket cell (Tuthill & Wilson, 2016). The dendrite tip attaches to the dome cuticle (Keil, 1997b). (D) Mechanosensory bristles detect deflection of the hair-like projection. They are innervated by a single neuron, the dendritic projections of which terminate at the base of the shaft. Specific to this bristle class, the most proximal epithelial cell to the developing sense organ is induced to become a bract cell (Tokunaga, 1962; Tobler, 1969). Bract cells secrete a thick, pigmented, hair-like, cuticular protrusion. (E) The chemosensory taste bristles of the leg differ in their morphology from mechanosensory bristles, appearing less heavily melanized and more curved. They also house a pore at the terminus of the shaft and lack bracts. Each is innervated by a single mechanosensory neuron and 4 gustatory receptor neurons (GRNs) (Nayak & Singh, 1983). Figure created using Biorender.com.

The function of sensory organs and their tuning to particular stimuli is not only a product of the neurons that innervate them. In each case, the sensory organ is a composite of multiple distinct cell types and dependent upon the involvement of glia to effectively relay detected signals to the brain. Different organ classes appear to share a common developmental blueprint, such that, despite variation in their form and function, mechanosensory bristles, chemosensory taste bristles, and campaniform sensilla each contain four homologous cell types (Lai & Orgogozo, 2004). In the bristle lineage, these are the neurons, which may vary in number between different sensory organ classes (such as between the polyinnervated chemosensory and monoinnervated mechanosensory bristles), along with three sensory support cells: the trichogen (shaft or, in campaniform sensilla, the dome), tormogen (socket), and thecogen (sheath). These sensory support cells bear features that clearly define the sensory capabilities of the organ. For example, the elongated shafts of mechanosensory bristles support the deflection-based mechanism through which stimuli are detected (Walker *et al*., 2000), the pore at the tip of chemosensory taste bristles enables the receipt of non-volatile compounds (Montell, 2009), and the elliptical shape of many campaniform sensilla confers sensitivity to the directionality of cuticular compression and strain (Pringle, 1938; reviewed by Tuthill & Wilson, 2016). But beyond these morphological features, our understanding of the wider, organ-specific contributions that support cells make to the specific sensory capabilities of each organ class remains poor (Prelic *et al*., 2022). Yet there is clear potential for their broader involvement in defining an organ type’s capabilities, given both their close physical associations with the neurons and, at least in taste bristles, their role in producing the lymph fluid that bathes the dendrites of GRNs and which is central to tastant detection (Ebbs & Amrein, 2007; Schmidt & Benton, 2020).

High-throughput single-cell RNA sequencing (scRNA-seq) technologies allow for the transcriptional profiles of many thousands of cells to be recorded from a single tissue. The advent of these technologies has precipitated an explosion of interest in cell type-specific patterns of gene expression. Over the last few years, ‘atlases’ describing the cellular diversity of tissues (Croset, Treiber, & Waddell, 2018; Hung *et al*., 2020; Allen *et al*., 2020; Rust *et al*., 2020), embryos (e.g. He *et al*., 2020; Mittnenzweig *et al*., 2021; Seroka, Lai, & Doe, 2022), and whole adult animals (e.g. Cao *et al*., 2017; Chari *et al*., 2021; Li *et al*., 2022) have been published for a variety of species. Through such work, regulators of development have been identified (e.g. Plass *et al*., 2018; Qiu *et al*., 2022), novel cell types described (e.g. Fu *et al*., 2020; Basil *et al*., 2022), and the effects of age (e.g. Almanzar *et al*., 2020; Zhang *et al*., 2021) and infection (e.g. Xie *et al*., 2020; Bomidi *et al*., 2021) on the gene expression profiles of individual cell types characterized. But in arthropods, tissues associated with the cuticle have presented a challenge to single-cell approaches because the cuticle prevents isolation of single cells from peripheral tissues without significant damage (McLaughlin *et al*., 2021; Li *et al*., 2022). Although single-nuclei RNA-seq methods have been used in such tissues (e.g. Li *et al*., 2022), scRNA-seq approaches generally offer substantially greater read and gene detection with reduced ‘gene dropout’ and lower expression variability between cells (Bakken *et al*., 2018). Consequently, techniques and approaches that help overcome these barriers are of significant value.

Here, we use scRNA-seq to profile the sensory organs of the male *D. melanogaster* foreleg at two developmental timepoints that follow the specification of sensory organ cells (24h and 30h after puparium formation, APF). Using a fine-scale dissection technique, we specifically target the first tarsal segment (the ‘basitarsus’) to maximize the detection of rare sensory organ types, including the campaniform sensilla, chemosensory taste bristles, and sex comb teeth. We begin by examining the transcriptomic landscape of the tissues in which the sensory organs reside, constructing a spatial reference map of epithelial cells based on intersecting axes of positional marker expression, resolving joint-specific gene expression, and characterizing the distinct repertoires of expressed genes in tendon cells, hemocytes, and bract cells. We then focus on the non-neuronal component of the nervous system, describing the complement of glial cells present in the region, identifying and visualizing wrapping glia, surface glia, and a novel axon-associated cell population that is negative for the canonical glia marker *repo* and appears to contribute towards the construction of the neural lamella. We then resolve and validate a combinatorial transcription factor code unique to the neurons of each of mechanosensory bristles, campaniform sensilla, chordotonal organs, and the sex comb. We further identify and validate a transcription factor code unique to four transcriptomically distinct GRN classes, including known male- and female-pheromone sensing neurons, and recover this same code in a published adult leg dataset. With these annotations in place, we link a wide range of genes, including receptors and membrane channels, to specific neurons. Finally, we detail the transcriptomic differences that distinguish between sensory organ support cells, both within a single organ class (e.g., sheaths vs. sockets) and between classes (e.g., chemosensory sheaths vs. mechanosensory sheaths).

## Results

### Homologous clustering of 24h and 30h transcriptomes

We generated two scRNA-seq datasets from freshly dissected male first tarsal segments using 10x Chromium chemistry. One sample comprised males collected at 24h APF and the other at 30h APF. After filtering based on cell-level quality control metrics (see Materials and methods), we recovered 9877 and 10332 cells in our 24h and 30h datasets, respectively (Figure 2–figure supplement 1A). In the 24h dataset, the median number of genes and transcripts detected per cell was 2083 and 11292, respectively (Figure 2–figure supplement 1B,D). The equivalent values for the 30h dataset were 1245 and 5050 (Figure 2–figure supplement 1C,E). We began our analysis by constructing separate UMAP plots for each dataset. The clustering pattern of the 30h dataset largely recapitulated that of the 24h dataset, which is to say that each cluster in the 24h dataset had a clear homologue in the 30h dataset (and *vice versa*) based on marker gene expression (Figure 2A-J). Given this concordance, we opted to integrate our two datasets and subcluster our data to facilitate closer analysis of rare cell subtypes (see Materials and methods). First, we subclustered epithelial cells and then separated them into joint and non-joint datasets (Figure 2K). Epithelial cells represented the major cell type in both datasets (70.7% in 24h: joints = 16.5%, non-joints = 54.1%; 70.2% in 30h: joints =13.6%, non-joints = 56.5%; Figure 2L). We then subclustered the non-epithelial cells (Figure 2M) and separated them into neurons (based on the expression of *fne*; Figure 2O), sensory support cells (based on the expression of *pros, nompA, Su(H)*, and *sv*; Figure 2Q), and the remaining non-sensory cells (Figure 2P). The relative proportions of cells in each of these classes was similar between the two datasets (24h: 19.5% neurons, 38.8% sensory support, 41.8% non-sensory; 30h: 17.1% neurons, 37.0% sensory support, 45.9% non-sensory; Figure 2N). We include our final annotations in Figure 2 and work through the supporting evidence throughout this paper, with an emphasis on sensory and glial cell types.

**Figure 2.**
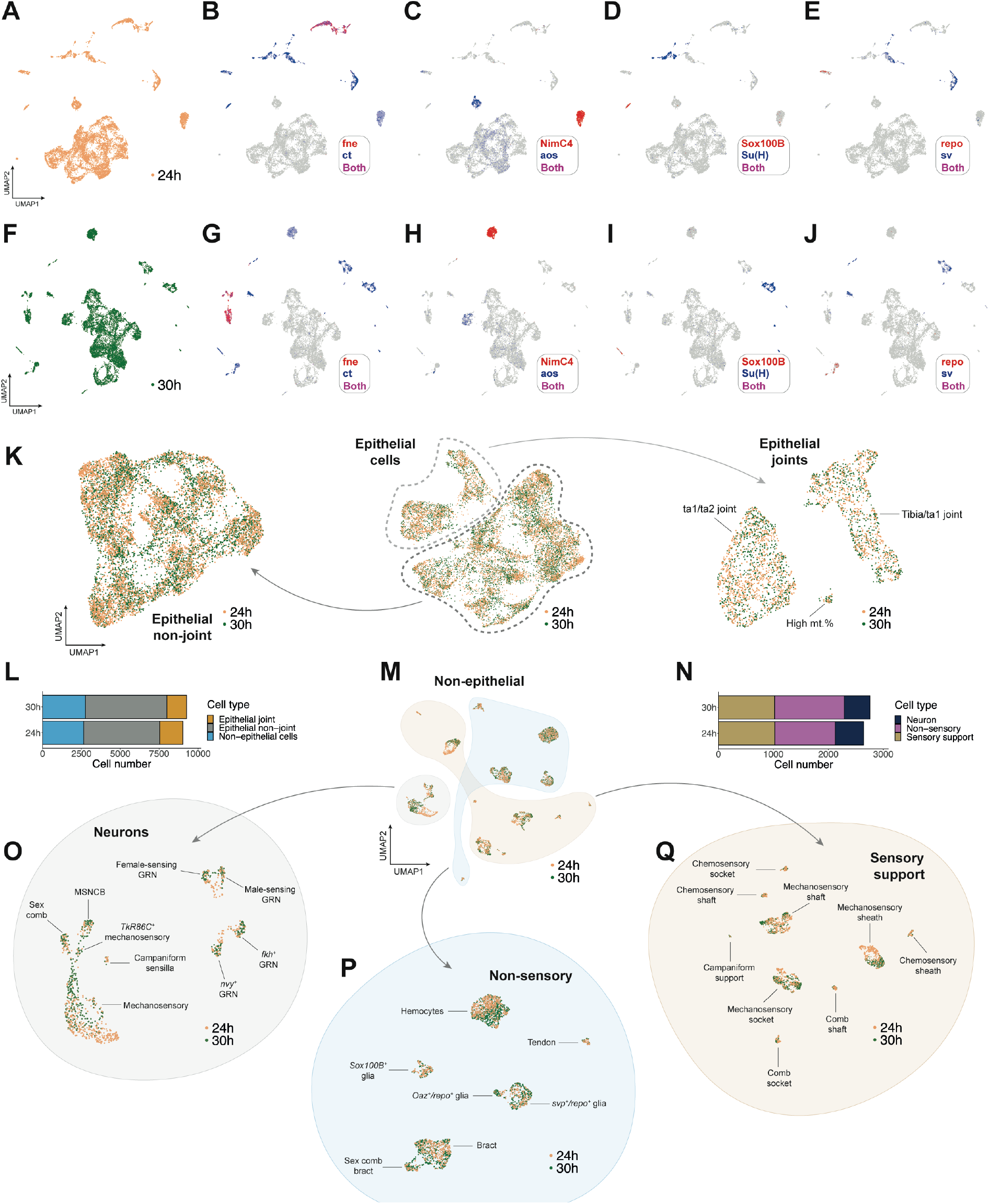
Clustering and iterative subsetting of integrated 24h and 30h APF scRNA-seq datasets identifies tarsal cell types with high resolution. (A-J) UMAP plots showing cell clustering in the 24h (A-E) and 30h (F-J) datasets separately. Expression of a series of cluster markers is overlaid on the full 24h (B-E) and 30h (G-J) dataset UMAPs. At this resolution, each higher-level cluster in the 24h dataset has a clear homologue in the 30h dataset based on a selected subset of marker genes and *vice versa. fne* for neurons; *ct* for non-epithelial cells; *NimC4* for hemocytes; *aos* for bracts; *Sox100B* and *repo* for different subtypes of glia and axon-associated cells; *Su(H)* for socket cells; *sv* for shafts and sheaths. (K) The central UMAP shows the clustering pattern observed in an integrated dataset containing just the epithelial joint and non-joint cells from both 24h (gold dots) and 30h (green dots) samples. Joints are circled with a dashed light grey line, non-joints with a dashed dark grey line. Reclustering the non-joint and joint cells gave rise to the two flanking UMAP plots. See Figure 3 for details on how the annotations were determined. (L) The number of cells in the post-filtration, doublet-removed epithelial joint (yellow), epithelial non-joint (grey), and non-epithelial cell (blue) datasets, plotted separately based on which sample (24h APF or 30h APF) the cells originated from. (M) UMAP showing the clustering pattern observed in an integrated dataset containing all non-epithelial cells from both the 24h (gold dots) and 30h (green dots) samples. Three major subsets of cells are grouped by coloured shapes: neurons, non-sensory cells, and sensory support cells. (N) The number of cells in the post-filtration, doublet-removed sensory support (gold), neuron (navy), and non-sensory (pink) datasets, plotted separately based on which sample the cells originated from (24h APF or 30h APF). (O-Q) UMAPs showing the clustering pattern observed in integrated datasets containing all neurons (O), non-sensory cells (P), and sensory support cells (Q) from both the 24h (gold dots) and 30h (green dots) samples. See Figures 4–8 for details on how the annotations were determined. GRN = gustatory receptor neuron; MSNCB = mechanosensory neuron in chemosensory bristle.

### Transcriptomic divergence between the tibia/tarsus and inter-tarsal joints

Joint cells separated from the main body of epithelial cells in our initial epithelial clustering analysis (circled in Figure 3A,B). These cells showed enriched expression of genes with known involvement in the formation of or localization at joints, including *drm*, *nub*, and *TfAP-2* (Figure 3A,B) (Kerber *et al*., 2001; Monge *et al*., 2001; Hao *et al*., 2003). After subclustering these cells (Figure 3C,D), we used the non-overlapping expression of *nub* and *TfAP-2* to identify sub-regions of the joint. We observed expression of GFP-tagged TfAP-2 (Kudron *et al*., 2018) in both the tibia/tarsal and inter-tarsal joints, with somewhat weaker staining in the former (Figure 3E). In contrast, expression of *nub-GAL4* was restricted to the joint-adjacent region of the distal tibia (Figure 3E)(see also Rauskolb & Irvine, 1999; Mirth & Akam, 2002). One of our joint clusters was *TfAP-2^+^/nub^−^* and additionally positive for *bab2*, a gene with a documented tarsus-restricted expression profile (Figure 3A-D; Godt *et al*., 1993). The other was divided into separate *TfAP-2^+^* and *nub^+^* domains. Based on our imaging, we propose that the *TfAP-2^+^/nub^−^* cluster corresponds to the ta1/ta2 joint, while the other corresponds to the tibia/ta1 joint. The tibia/ta1 joint can be further subdivided into the *nub*^+^ proximal and *TfAP-2*^+^ distal regions (Figure 3F; see Figure 3–figure supplement 1A for details of the high mt% cluster).

**Figure 3.**
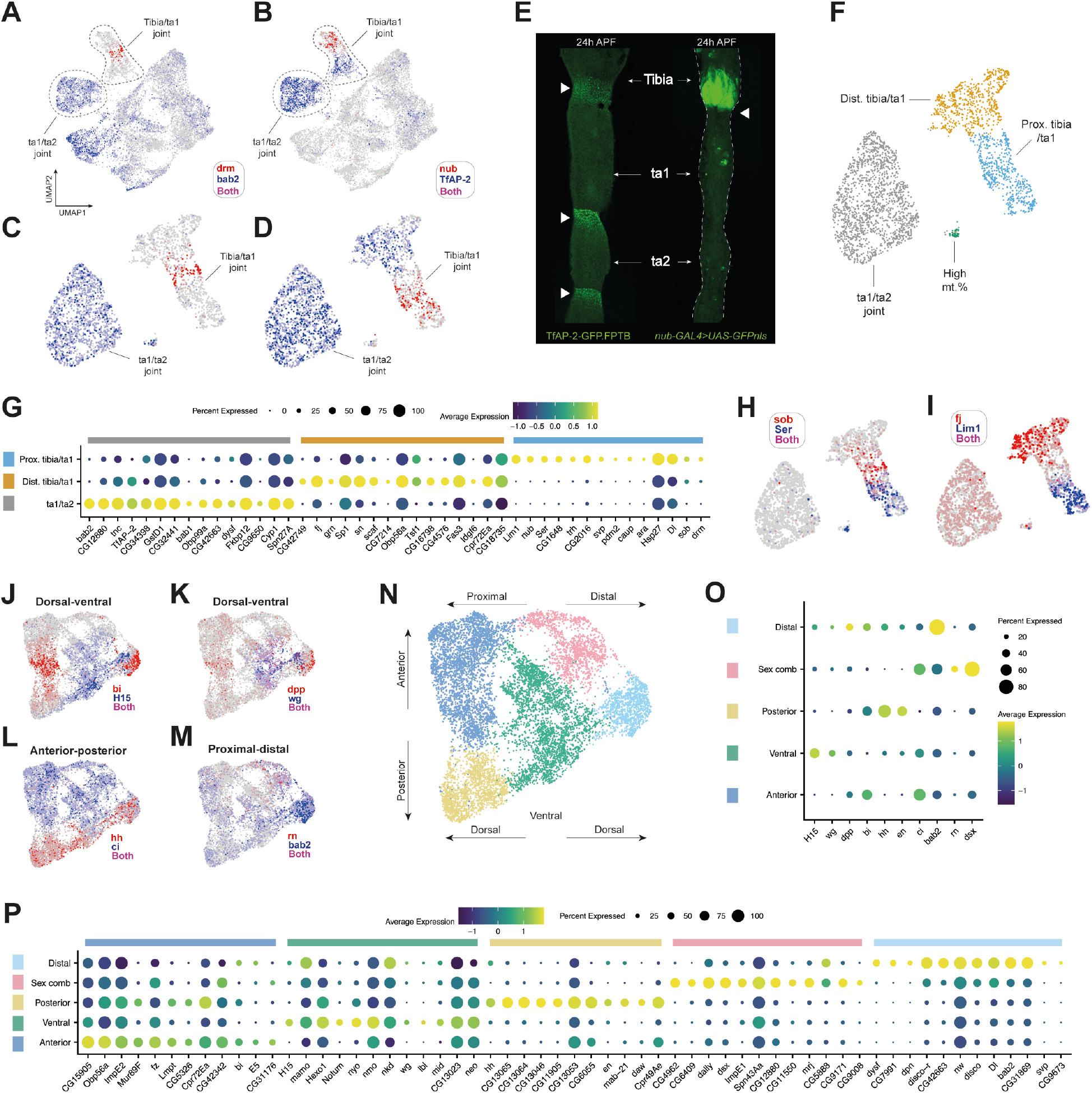
Single-cell sequencing recovers positional information in the leg epithelium. (A-B) UMAP plots of the integrated epithelial joint and non-joint dataset overlaid with the expression of (A) *drm* (red) and *bab2* (blue), and (B) *nub* (red) and *TfAP-2* (blue). The two major joint clusters are circled and can be distinguished from one another based on the expression of these 4 genes. (C-D) As in (A) and (B) but with a UMAP plot of just the joint cells. (E) Confocal images showing 24h pupal legs. On the left is a leg from a *TfAP-2-GFP* male. Staining is concentrated at the joints, which are each marked with a white triangle. Staining appears stronger at the inter-tarsal joints compared to the tibia/ta1 joint, consistent with the expression pattern of *TfAP-2* in the scRNA-seq data. On the right is a leg from a *nub-GAL4 > UAS-GFP.nls* male. Staining is concentrated in the distal tibia, proximal to the tibia/ta1 joint (marked with a white arrow). Note that some non-specific staining from contaminating fat body is present in ta1 and ta2. (F) Joint UMAP with clusters identified through shared nearest neighbour clustering and annotated based on the data presented in A-E. For details on the high mt. % cluster, see Figure 3–figure supplement 1A. (G) Dot plot of a selection of top marker genes for each of the joint clusters given in F (excluding the high mt. % cluster). Marker genes were identified by comparing each cluster to the remaining joint clusters. Dot size reflects the number of cells in the cluster in which a transcript for the marker gene was detected, while colour represents the expression level. (H-I) Expression of a selection of top marker genes identified in the analysis presented in G overlaid on the joint UMAP plot. (H) *sob* (red) and *Ser* (blue). (I) *fj* (red) and *Lim1* (blue). (J-M) UMAP plots of the non-joint epithelial cells overlaid with markers of spatial identity. Both (J) and (K) show expression of dorsal (red: *bi* and *dpp*) and ventral (blue: *H15* and *wg*) markers. (L) shows anterior (blue: *ci*) and posterior (red: *hh*) markers. (M) shows proximal (blue: *bab2*) and distal (red: *rn*) markers. As is clear from the expression patterns, separation based on spatial markers is apparent for each axis, although stronger for dorsal-ventral and anterior-posterior than proximal-distal. This is likely due to us recovering only a small fraction of the proximal-distal axis by focusing in on just a single tarsal segment. (N) UMAP plot of the non-joint epithelial cells coloured by cluster identity as determined through shared nearest neighbour clustering. Spatial axes are illustrated by arrows based on the expression data presented in J-M. (O) A dot plot showing the expression of positional markers across each cluster given in (N). Clusters are assigned to regions based on the positional gene expression signature they display. (P) A dot plot of the top markers for each cluster given in (N, O). Marker genes were identified by comparing each cluster to the remaining non-joint epithelial clusters.

To understand the broader transcriptomic differences between joint regions, we tested for differential gene expression in each cluster compared to all other clusters in our joint dataset (a selection is given in Figure 3G). We recovered several genes with known roles in joint formation, including the *odd-skipped* family transcription factors *drm* and *sob* (Figures 3A,H; Figure 3–figure supplement 1B,C). These were enriched at the interface between the proximal and distal tibia/ta1 clusters, consistent with their known absence from the upper tarsal segments (Hao *et al*., 2003). We also looked at two further members of the same gene family: *odd*, which showed a similar expression pattern but was present in fewer cells, and *bowl*, which was widely expressed among the joint clusters and in the wider body of epithelial cells (Figure 3–figure supplement 1D,E-G). We observed two major patterns among the top differentially expressed genes (DEGs) defining the proximal region of the tibia/ta1 joint. First, genes that appeared widespread among epithelial cells, but which were excluded from the other joint clusters (*e.g. CG1648* and *Ser;* Figure 3–figure supplement 1H,I). Second, and more commonly, there were genes that showed strong specificity to the proximal tibia/ta1 joint (*e.g. Lim1* – Figure 3I & see also Figure 5V for stainings – *nub*, *trh, caup, ara, pdm2, CG2016*, and *svp*; Figure 3–figure supplement 1J-P). To the best of our knowledge, no role for *pdm2* in leg development has been characterized, but it is believed to have evolved via duplication of *nub*, to which it is adjacent in the genome (Ross *et al*., 2015). Our data suggests that the two are co-expressed, a conclusion that’s further supported by the recent discovery that the two genes share enhancers in the wing (Loker & Mann, 2022). In contrast to the distinctiveness of the proximal tibia/ta1 expression profile, there was greater overlap in the DEGs for the distal tibia/ta1 and ta1/ta2 joints, with fewer cluster specific genes (Figure 3G; *e.g. fj*, Figure 3I; For UMAPs and discussion see Figure 3–figure supplement 1Q-V and Figure 3–figure supplement 2A-R). This suggests that while the transcriptomic distinctiveness of the proximal tibia/ta1 region is dominated by qualitative differences in expression, the more distal joint clusters show a more quantitative signal.

**Figure 4.**
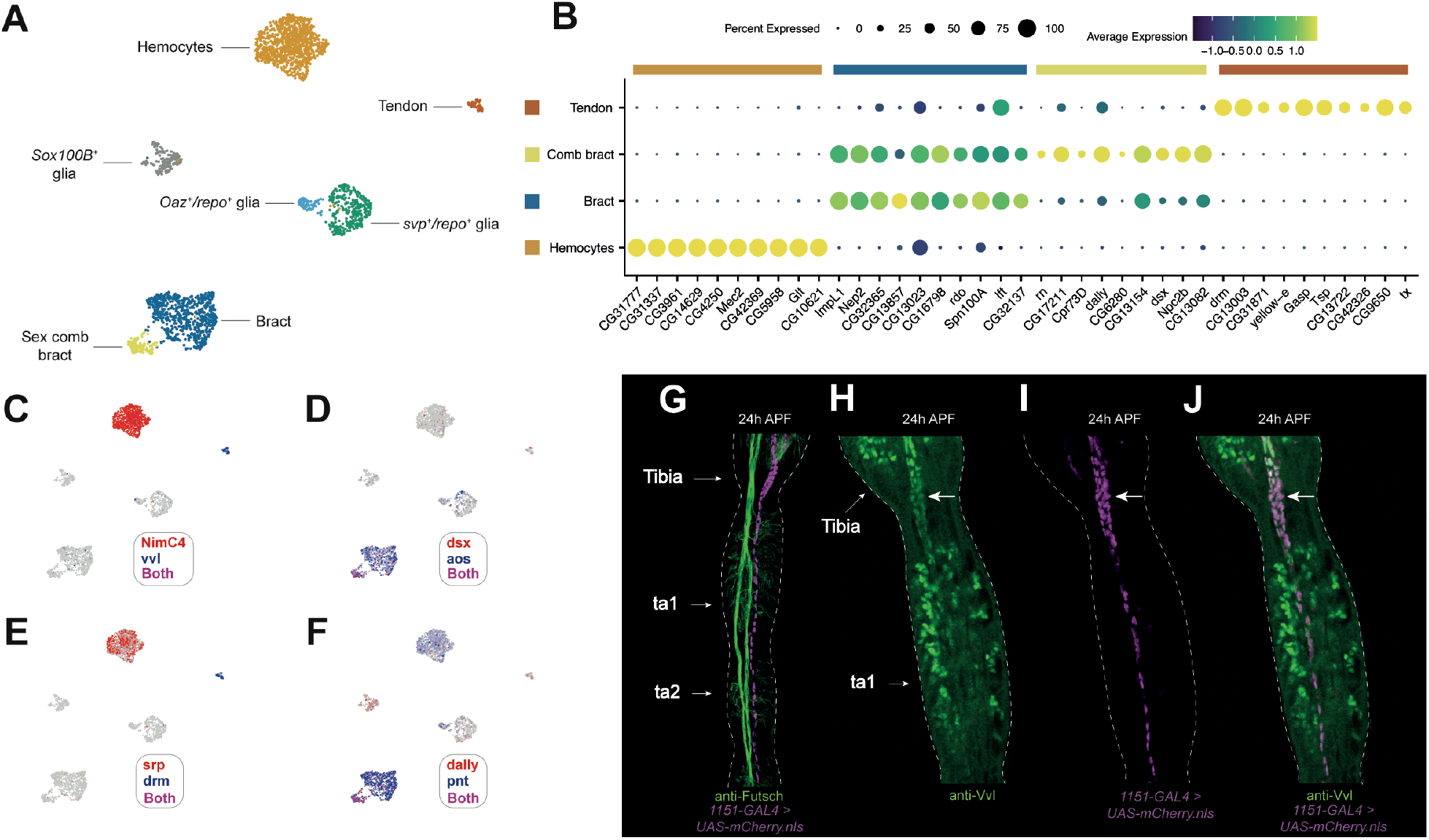
The tarsus contains several types of non-sensory, non-epithelial cells. (A) Annotated UMAP plot of non-sensory cells. (B) Dot plot of the expression of top differentially expressed genes identified through comparisons between each named cluster in (B) and all remaining clusters in (A). Glia are discussed in Figure 5. (C-F) The non-sensory UMAP shown in (A) overlaid with expression of key marker genes for each cluster. Note that *dsx* (D) and *dally* (F) are expressed in a distinct subset of bract cells, which likely corresponds to sex comb bracts. (G) 24h APF male pupal upper tarsal segments showing staining from *1151-GAL4* > *UAS-mCherry.nls* (magenta) and the neuronal marker anti-Futsch (green). *1151-GAL4* marks tendons (Soler *et al*., 2004). The arrangement of tendon cells is clearly distinct from the paired nerve fibers that run along the same axis. (H-J) 24h APF first tarsal segment and distal tibia from an *1151-GAL4* > *UAS-mCherry.nls* (magenta) male counter-stained with anti-Vvl (green). Co-staining is clearer in the levator and depressor tendons at the distal tibia/ta1 joint (marked with an arrow) than in the long tendon, which extends along the proximal-distal axis of the tarsal segments. This may be due to the greater concentration of tendon cells in this region and difficulties distinguishing between anti-Vvl staining in mechanosensory bristle cells (see Figure 6Q-S) and tendon cells.

**Figure 5.**
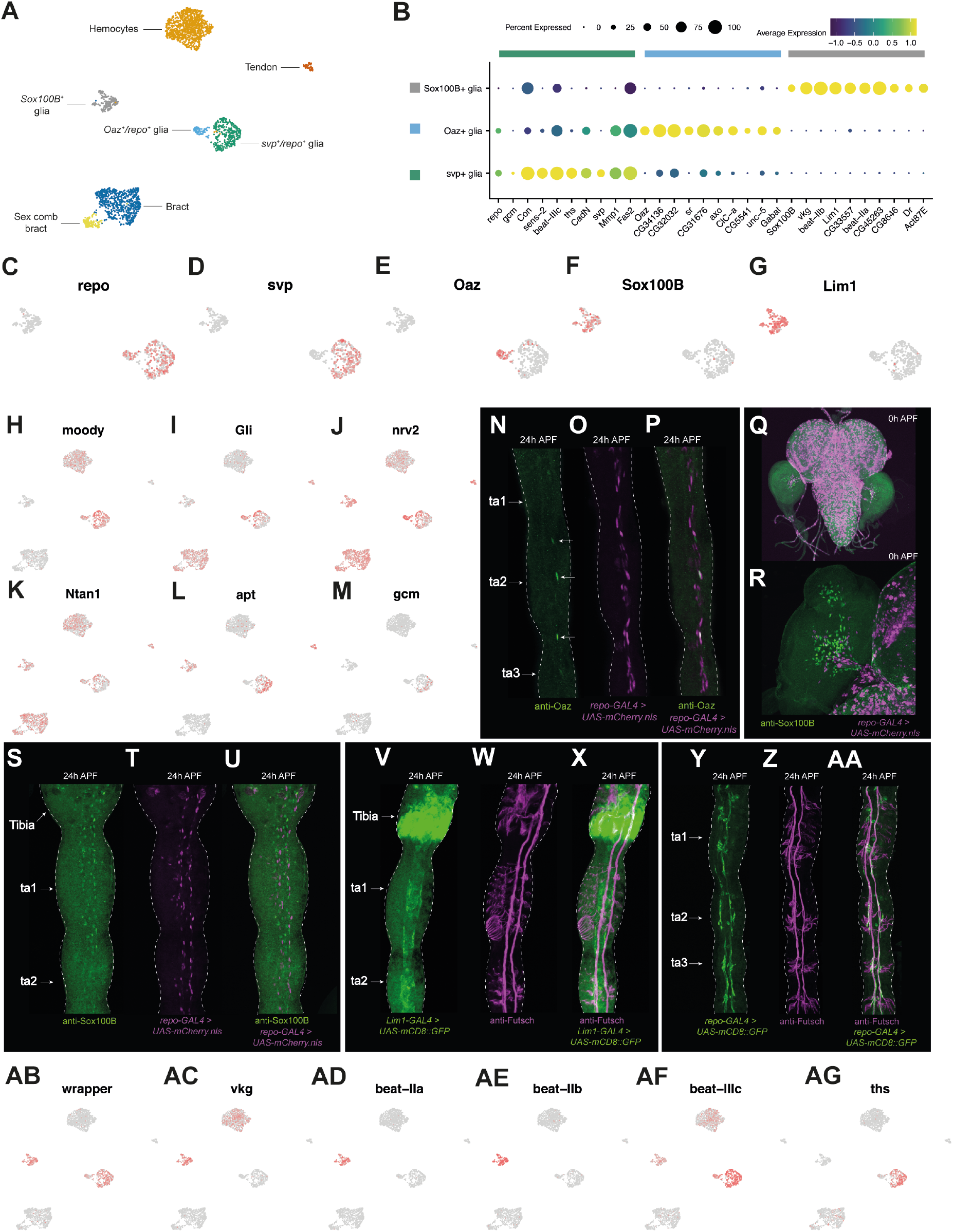
Non-canonical expression patterns in leg glia and a new cell type associated with the neural lamella. (A) Annotated UMAP plot of non-sensory cells. While the *Oaz*^+^ and *svp*^+^ glia cells express the canonical glia marker *repo*, the *Sox100B^+^* cells do not. (B) Dot plot of the expression of *repo* and *gcm*, the canonical glia markers, along with top differentially expressed genes identified through comparisons between each of *Sox100B^+^* glia, *Oaz^+^*/*repo^+^* glia, and *svp^+^*/*repo^+^* glia against all the clusters named in (A). Of these genes, *sr* is known to induce tendon cell fate and to the best of our knowledge, no functions have previously been reported for *sr* in glia. *Unc-5* and *Fas2* are both required for glial migration (reviewed in Yildirim *et al*., 2019). Genes identified as differentially expressed through between-glia comparisons are given in Figure 5–figure supplement 1A. (C-G) UMAP plots of the subsetted *Sox100B^+^* glia, *Oaz^+^*/*repo^+^* glia, and *svp^+^*/*repo^+^* glia from (A) overlaid with the expression of a series of top marker genes for each cluster. (H-M) The UMAP plot shown in (A) overlaid with the expression of a series of glia markers. *moody* and *Gli* are subperineural glia markers, *nrv2* and *Ntan1* are wrapping glia markers, *apt* is a surface glia marker (*i.e*. a marker of both perineural and subperineural glia), and *gcm* is the upstream determinant of glial identity (Xie & Auld, 2011; Sasse & Klämbt, 2016). (N-P) 24h APF legs from *repo-GAL4>UAS-mCherry.nls* males counterstained with anti-Oaz, a marker of wrapping glia (Lassetter *et al*., 2021). Oaz^+^ cells are denoted by an arrow in the left-hand image. In Oaz^+^ cells at 24h APF, the *repo-GAL4^+^* signal was often weak and in one of the four legs we imaged, the one shown here, we observed a single *Oaz^+^* cell that appeared *repo-GAL4^−^*. This is the topmost of the Oaz^+^ cells to which an arrow is pointing. (Q-R) Brain, ventral nerve cord, and leg discs (Q) and a closeup of a leg disc (R) from *repo-GAL4>UAS-mCherry.nls* males counterstained with anti-Sox100B. Note how *repo-GAL4^+^* cells can be seen migrating into the disc from the CNS, while Sox100B^+^ cells appear to originate within the disc itself. (S-U) 24h APF male upper tarsal segments from *repo-GAL4 > UAS-mCherry.nls* counterstained with anti-Sox100B. Both show a similar, but non-overlapping, distribution of stained cells. (V-X) 24h APF male upper tarsal segments from *Lim1-GAL4 > UAS-mCD8∷GFP* counterstained with anti-Futsch. Above the tibia/ta1 joint, *Lim1-GAL4* was expressed in the epithelial cells of the distal tibia, as predicted by our epithelial joint analysis (Figure 3I). Below the joint, the staining surrounded and spanned the distance between the two central axon trunks into which the sensory neuron axons project. (Y-AA) 24h APF male upper tarsal segments from *repo-GAL4 > UAS-mCD8∷GFP* counterstained with anti-Futsch. Unlike the *Lim1-GAL4, repo-GAL4* staining does not span the gap between the two axon trunks with which it is closely associated, and cell bodies are clearly seen branching away from the fibers. (AB-AG) The UMAP plot shown in (A) overlaid with the expression of a series of top markers identified in this study.

### Epithelial cells express a signature of anatomical position

We next turned to the largest portion of our dataset, the non-joint epithelial cells. Here, we observed clear separation between dorsal and ventral cells. This separation was clearest when mapping the expression of *H15* (ventral) and *bi* (dorsal)(Figure 3J). *wg* (ventral) and *dpp* (dorsal) showed a similar, albeit weaker separation (Figure 3K). Unlike the joints, anterior-posterior separation was also clear, as delineated by the expression of *ci* (anterior) and *hh* (posterior)(Figure 3L). A weaker signature of proximal-distal separation could also be discerned from the expression of *bab2*, which is absent from the tibia and increases in expression between ta1 and ta2 (reviewed in Kopp, 2011)(Figure 3M). We also detected localized expression of *rn*, the expression of which in ta1 is limited to the distal region (Natori *et al*., 2012)(Figure 3M). Recovery of spatial patterning in epithelial cells has recently been demonstrated in *Drosophila* wing imaginal disc scRNA-seq data (Deng *et al*., 2019; Everetts *et al*., 2021). But unlike in wing discs, we find that in this region of the leg the anterior-posterior signature is stronger than the proximal-distal, which may be due to our sequencing only a small fraction of the proximal-distal axis (*i.e*., just one tarsal segment). We then used the intersecting axes of positional marker expression as a spatial reference map to assign clusters to regions of the dissected leg tissue (Figure 3N,O). We tested for DEGs by comparing each region to the remainder (Figure 3P). Many DEGs showed signatures of localized upregulation rather than cluster-specific expression, as would reasonably be expected from a tissue composed of a single cell type (exceptions include *lbl, CG13064, CG13065*, and *CG13046*; Figure 3–figure supplement 2S-V). But the cluster enriched for the distal ta1 marker *rn* exhibited a more specific gene expression profile, including showing enriched expression of the effector of sex determination, *dsx* (reviewed in Hopkins & Kopp, 2021), consistent with the localization of the sex comb to this region (Robinett *et al*., 2010; Tanaka *et al*., 2011). Genes enriched here represent candidate components of the sex-specific gene regulatory network that drives sex comb rotation (Figure 3–figure supplement 2W-AE).

### Pupal leg hemocytes form a uniform population

We identified a single hemocyte cluster based on enriched expression of *He*, *Hml, srp*, and Nimrod-type receptor genes (Figure 4C,E; Figure 4–figure supplement 1A-N)(Evans, Hartenstein, & Banerjee, 2003; Kocks *et al*., 2005; Kurucz *et al*., 2007). In our differential gene expression analysis, we observed strongly hemocyte-enriched expression of genes including *CG31777, CG31337, CG3961, CG14629, CG4250, Mec2, CG42369, CG5958, Glt*, and *CG10621* (Figure 4B; see Figure 4–figure supplement 1X-AG for UMAPs of the first 5 genes). To test for the presence of hemocyte subtypes, we ran a wide panel of recently identified lamellocyte, crystal cell, and plasmatocyte subtype-specific genes against our data (Tattikota *et al*., 2020). However, we saw no obvious subclustering in relation to these genes: they were either widely expressed among hemocyte cells, too patchily expressed to reflect a clear subpopulation, or absent from our dataset (Figure 4–figure supplement 1O-W). The absence of clear subclustering may reflect the rarity of these hemocyte subpopulations, that recovered cells are insufficiently differentiated at these timepoints to discriminate subclasses, that these notably fragile cells (Rizki & Rizki, 1959) lyse during tissue dissociation, or point to differences between the larval and pupal immune cell repertoire, for which there is some evidence (Grigorian, Mandal, & Hartenstein, 2011; Hultmark & Andó, 2022).

### The induction of bract identity is accompanied by a transcriptomic shift away from epithelial cells

Most but not all mechanosensory bristles on the legs are associated with a bract cell (Hannah-Alava, 1958; Reed, Murphy, & Fristrom, 1975). We identified bracts based on the expression of the transcription factor *pnt* and *aos*, an EGF inhibitor selectively expressed in cells assuming bract fate (del Álamo, Terriente, & Díaz-Benjumea, 2002; Peng, Han, & Axelrod, 2012; Mou *et al*., 2012). Among our non-sensory clusters, we find two enriched for *aos* and *pnt* (Figure 4D,F; Figure 4–figure supplement 2A-D). Differential gene expression analysis comparing among our non-sensory cell clusters showed that the top markers of the major bract cluster also showed elevated expression in the minority cluster, but that the top markers of the minority cluster were more specific in their expression (Figure 4B). The top DEGs for the minority cluster included several genes that were among the top markers of the putative sex comb bearing region identified in our epithelial cell analysis (*rn, dsx*, and *dally*; Figure 4D,F). Thus, these cells likely correspond to sex comb bracts. Given that bract identity is induced in epithelial cells, rather than emerging through the sensory organ precursor lineage, the natural comparison to make to identify putative determinants of bract fate is to compare bract cells with other epithelial cells (Tokunaga, 1962; Tobler, 1969; Held, 1979, 2002; Lawrence, Struhl, & Morata, 1979; del Álamo *et al*., 2002). Comparing the two bract clusters with the non-joint epithelial cells, we observed several genes that were highly enriched in bracts and largely absent from epithelial cells (Figure 4–figure supplement 2E). It was common to find expression of some of these genes (*e.g. CG33110, Nep2, CG32365, neur*; Figure 4–figure supplement 2F-P) in bristle shaft and socket cells, which may reflect both the crucial role played by bristle support cells in inducing bract cell identity (Peng *et al*., 2012) and, considering the short bristle-hair like protrusion that bracts develop, their partially overlapping morphological characteristics. The distinct expression profile of the bracts suggests that induction of this identity in epithelial cells is followed by remodeling of its transcriptome.

### Tendon cells express the POU transcription factor *vvl*, but not *sr*, between 24h and 30h APF

We identified a cluster of tendon cells based on the expression of *Tsp, tx*, and the joint marker *drm* (Figure 4B,E)(Armand *et al*., 1994; Subramanian *et al*., 2007; Chanana *et al*., 2007). To visualize the anatomical distribution of tendon cells in the focal leg region, we crossed the verified tendon marker line *1151-GAL4* (Soler *et al*., 2004) to *UAS-mCherry.nls* and counter-stained with an antibody against the neuronal marker Futsch to distinguish between tendons and axonal trunks (Figure 4G). The dissected region contains the ‘long tendon’, which runs along the proximal-distal axis of the tarsal segments, as well as the distal portion of the ‘tarsus levator’ and ‘tarsus depressor’ tendons, which are housed in the distal tibia (Soler *et al*., 2004). One of the top DEGs for our *Tsp^+^/tx^+^* cells was the POU homeobox transcription factor *vvl* (Figure 4C). When counter-staining *1151-GAL4>UAS-mCherry.nls* legs with an antibody raised against Vvl we observed clear co-staining, supporting a tendon cell identity for this cluster (Figure 4H-J). Of the top DEGs we identified for tendon cells, none were entirely specific to this cluster when looking across all cells in the dataset. A common pattern was to see localized expression among epithelial cells (*e.g. drm, Tsp, CG13003*) or among sockets and shaft cells (*e.g. tx* and *CG42326*)(Figure 4–figure supplement 3A-R). Beyond the top 10 tendon cell DEGs, we detected significantly enriched expression of *trol*, which encodes the extracellular matrix proteoglycan Perlecan (validated using *trol-GAL4*; Figure 4–figure supplement 3S-V). Surprisingly, we did not detect enriched expression of the tendon-specifying transcription factor *sr* in this cluster, nor its tendon-specific downstream targets *slow* and *Lrt* (Wayburn & Volk, 2009; Gilsohn & Volk, 2010), which may be due to the relatively early developmental window during which we sequenced these cells (Figure 4–figure supplement 3W-AE).

### Glial cells in the developing first tarsal segment have non-canonical expression profiles

Three major classes of glia have been described in the leg: perineural and subperineural glia (which collectively comprise the surface glia), and the PNS-specific wrapping glia (Sasse & Klämbt, 2016). To identify these populations in our dataset, we mapped the expression of the canonical glia marker *repo*, which is thought to be expressed in all lateral/embryonic glial cells except for a subset of specialized wrapping glia in the CNS, known as midline glia (Xiong *et al*., 1994; Halter *et al*., 1995; Yuasa *et al*., 2003; Beckervordersandforth *et al*., 2008; Yildirim *et al*., 2019). We observed two *repo^+^* clusters, one of which was *svp^+^* and the other *Oaz*^+^ (Figure 5A–E). Previous work has shown that Oaz specifically labels wrapping glia in larval peripheral nerves, where it is coexpressed with Repo (Lassetter *et al*., 2021). In our stainings, we observed that anti-Oaz labelled a small number of *repo-GAL4^+^* cells (Figure 5N-P). Beyond *Oaz*, however, the expression patterns of the *repo^+^* cells were atypical with respect to known marker genes. The subperineural glia marker genes *moody* and *Gli* were widely expressed across *repo^+^* cells, including in the *Oaz^+^* putative wrapping glia cells (Figure 5H,I)(Xie & Auld, 2011; Sasse & Klämbt, 2016). Conversely, the wrapping glia markers *nrv2* and *Ntan1* weren’t restricted to the *Oaz^+^* cluster (Figure 5J,K), while the perineural glia marker *Jupiter* was widely detected across all non-sensory cells (Figure 5–figure supplement 1G). Indeed, genes previously shown to be enriched in surface glia (DeSalvo *et al*., 2014) were similarly represented in the DEGs of each cluster (*svp^+^* glia: 20/285, ~7.0%; *Oaz^+^* glia: 14/223, ~6.3%; top DEGs are given in Figure 5B, with an expanded list in Figure 5–figure supplement 1A).

Despite these surprisingly broad expression patterns of established marker genes, two features point to known identities. First, *svp-*lacZ is known to be expressed in all surface glia (Beckervordersandforth *et al*., 2008); among non-sensory cells in our data, *svp* was restricted to the *Oaz^−^/repo^+^*cells (Figure 5D). Second, among *repo^+^* cells the surface glia markers *apt* and *Gs2* (Sasse & Klämbt, 2016) appeared similarly restricted to the *Oaz^−^* cells (Figure 5L; Figure 5–figure supplement 1H). Collectively, this suggests that our *Oaz^+^* population correspond to wrapping glia, while the *svp^+^* population corresponds to surface glia. This latter population appears heterogeneous, with localized upregulation of *Gli* detectable (Figure 5I). Several processes may underlie this heterogeneity. One possibility is that it reflects developmental staging differences among surface glia. This is supported by the localized expression of the upstream determinant of glial identity, *gcm*, which is known to act early in development (Figure 5M)(Stork, Bernardos, & Freeman, 2012). Another possibility is that this cluster includes a mix of both perineural and subperineural glia. Perineural glia are far more numerous than subperineural glia in the leg (Sasse & Klämbt, 2016), so any we recover may present as a subregion within a surface glia cluster otherwise dominated by perineural glia. Finally, surface glia may naturally be heterogenous in their expression profiles, as has been suggested by others (Sasse & Klämbt, 2016).

### A novel cell type associated with the neural lamella

Beyond the *repo^+^* clusters, we observed a *repo^−^* population, enriched for expression of the transcription factors *Sox100B* and *Lim1* (Figure 5F,G), that expressed the midline glia marker *wrapper* (Figure 5AB; Noordermeer *et al*., 1998; Wheeler *et al*., 2009; Stork *et al*., 2009). To verify the mutually exclusive expression patterns of *repo* and *Sox100B*, we counterstained *repo-GAL4 > UAS-mCherry.nls* legs with an anti-Sox100B antibody at two timepoints. First, we looked at 0h leg discs (Figure 5Q,R). Here, we observed no overlap in expression but noted a difference in the behaviour of these two cell populations: while *repo-GAL4^+^* cells could be seen migrating into the leg disc, as CNS-derived glia are known to do (Sasse & Klämbt, 2016), Sox100B^+^ cells appeared to instead originate within the leg disc itself. At 24h APF, we also detected no overlap in expression. Despite being Repo^−^, at this timepoint the nuclei of Sox100B^+^ cells occupied glia-like positions in relation to the axon trunks (Figure 5S-U). The morphology and neuronal associations of Sox100B^*+*^ and *repo-GAL4^+^* cells were also distinct. By expressing a membrane tethered form of GFP under the control of a GAL4 driver for one of the top markers of the *Sox100B^+^* cluster (*Lim1-GAL4*; Suzuki *et al*., 2016), we observed that these cells appear to surround the axon trunks (Figure 5V-X). When compared to the equivalent staining for *repo-GAL4*, the staining around the axon trunks in *Lim1-GAL4 > UAS-mCD8∷GFP* legs appeared larger in diameter, suggesting that it comprises a layer that is outer to that of the *repo-GAL4^+^* cells (Figure 5Y-AA; Figure 5–figure supplement 1B-C). To exclude the possibility that these cells correspond to myoblasts, we compared the distribution of Sox100B^+^ cells to those stained by an antibody raised against the myoblast marker Mef2 (Figure 5–figure supplement 1D). Unlike Sox100B^+^ cells, myoblasts were restricted to the tibia and absent from tarsal segments. We have also recovered separate myoblast and Sox100B^+^ cell populations in a 16h APF scRNA-seq dataset (data not shown), further arguing against a myoblast identity for Sox100B^+^ cells.

We next visualized a protein trap of one of the *Sox100B^+^* cluster’s most specifically enriched markers, *vkg*, which encodes a subunit of the extracellular matrix component Collagen IV and which was essentially absent from the *repo^+^* glia (Figure 5AC; Figure 5–figure supplement 1E). The vkg∷GFP staining pattern resembled *Lim1-GAL4>UAS-mCD8∷GFP*: a broad ensheathing of the axon trunks that extended more laterally than the equivalent staining observed for *repo-GAL4* (Figure 5–figure supplement 1E). This provides further support for the ‘outerness’ of these cells. vkg∷GFP is known to label the neural lamella, a dense network of extracellular matrix that surrounds the central and peripheral nervous systems and which is required to help control the shape of the nervous system (Stork *et al*., 2008; Xie & Auld, 2011; Meyer, Schmidt, & Klämbt, 2014). These features suggest that the *Sox100B^+^* cells we identify here may be involved in the construction of the neural lamella. This role is thought to be performed by migrating hemocytes during embryogenesis (Olofsson & Page, 2005; Xie & Auld, 2011) and consistent with this we find that leg hemocytes also express *vkg*, albeit less strongly (Figure 5AC). But beyond this, the only other similarity we detect between our hemocyte and *Sox100B^+^* cells was the expression of *NimC3*, which was highly expressed in hemocytes and showed low-level expression in *Sox100B^+^* cells (Figure 4–figure supplement 1K). It’s therefore possible that migrating hemocytes work alongside *Sox100B^+^* cells to construct the neural lamella. Consistent with this, we see both cluster-specific and overlapping expression of many extracellular matrix component genes: *SPARC* is expressed in *Sox100B^+^* cells, hemocytes, and *repo^+^* glia; *trol* shows low-level expression in *Sox100B^+^* cells, hemocytes, and *svp^+^/repo^+^* glia; *Col4a1* and *vkg* are present in both *Sox100B^+^* cells and hemocytes; and *Pxn* is hemocyte-specific (Figure 5AC; Figure 5– figure supplement 1I-L). However, the position of *Sox100B^+^* cells at the outer layer of glia, along with their strong enrichment of the canonical neural lamella marker *vkg*, suggest that this cell type may play the primary role in neural lamella synthesis, with any contribution by the hemocytes being secondary.

### *repo^+^* glia and *Sox100B^+^* cells express distinct cell-cell communication gene repertoires

Comparing between the *repo^+^* glia and *Sox100B^+^* cells, we observed cluster-specific expression of beaten path family genes. *beat-IIa* and *beat-IIb* were restricted to the *Sox100B^+^* cluster, while *beat-IIIc* was enriched in *repo*^+^ cells (Figure 5AD-AF). *beaten path* genes are thought to act as neuronal receptors for sidestep gene family ligands expressed in peripheral tissues (Li *et al*., 2017b). Thus, the differential expression of different subsets of beaten path family genes between *repo^+^* and *Sox100B^+^* cells point both to the importance of these genes in the non-neuronal component of the nervous system and to cell type-specific patterns of between-cell communication in the developing nervous system. These *beat* genes have been recorded elsewhere in the fly: in the visual system, *beat-IIb* is expressed in L3 and L4 lamina neurons, the glia beneath L5, and in the lamina neuropil, while *beat-IIIc* is expressed in a subset of retinal neurons (Tan *et al*., 2015). Other cell communication pathway elements also showed cell type-specificity. For example, we observed that the Fibroblast Growth Factor (FGF) ligand *ths* was, among glia, largely restricted to *svp^+^*/*repo*^+^ cells (Figure 5AG). FGF signaling is known to underlie aspects of neuron-glia communication in *Drosophila*, although in these cases the source of Ths is neuronal. In one example, Ths is thought to act in neurons as a directional chemoattractant for the migration of and outgrowth of processes from astrocytes (Stork *et al*., 2014). In two others, the release of Ths from olfactory neurons directs ensheathing glia to wrap each glomerulus (Wu *et al*., 2017), while Ths in photoreceptor neurons induces differentiation of glia in the developing eye (Franzdóttir *et al*., 2009). The expression of *ths* in one of our glia populations, specifically the population we believe to correspond to the surface glia, is therefore surprising. It’s possible that FGF-mediated interactions between different glia populations guide their concerted differentiation and the development of the close physical associations they form. Similar interactions with neurons may also help guide the growth of neuronal projections in the vicinity of these glia.

### A combinatorial transcription factor code for sensory neurons

We recovered multiple distinct sensory neuron populations in our clustering analysis, each defined by the expression of a unique combination of transcription factors (Figure 6A,B). We recovered a similar clustering pattern in our analysis of male neurons from the Fly Cell Atlas (FCA) adult leg data (Figure 6G,H)(Li *et al*., 2022). In contrast to our pupal data, however, the FCA dataset is derived from all segments of all 3 pairs of legs and is single-nuclei, rather than single-cell. We failed to recover clear sex comb or MSNCB (mechanosensory neuron in chemosensory bristle) populations in the FCA data. But in their place, we recovered three populations apparently absent from our pupal dataset. These novel clusters were enriched for *CG9650*, a transcription factor we found to be enriched in joint and tendon cells, and showed cluster-specific, combinatorial expression of transcription factors, including *erm* and *bab1* (Figure 6I,J). Consistent with signs of a joint identity, we believe these are likely to correspond to chordotonal neuron populations, a class that is absent from the first tarsal segment. Despite the concordance between pupal and FCA neurons in terms of clustering pattern, we found they integrated poorly (Figure 6–figure supplement 1I,J) and therefore opted to analyze them separately.

**Figure 6.**
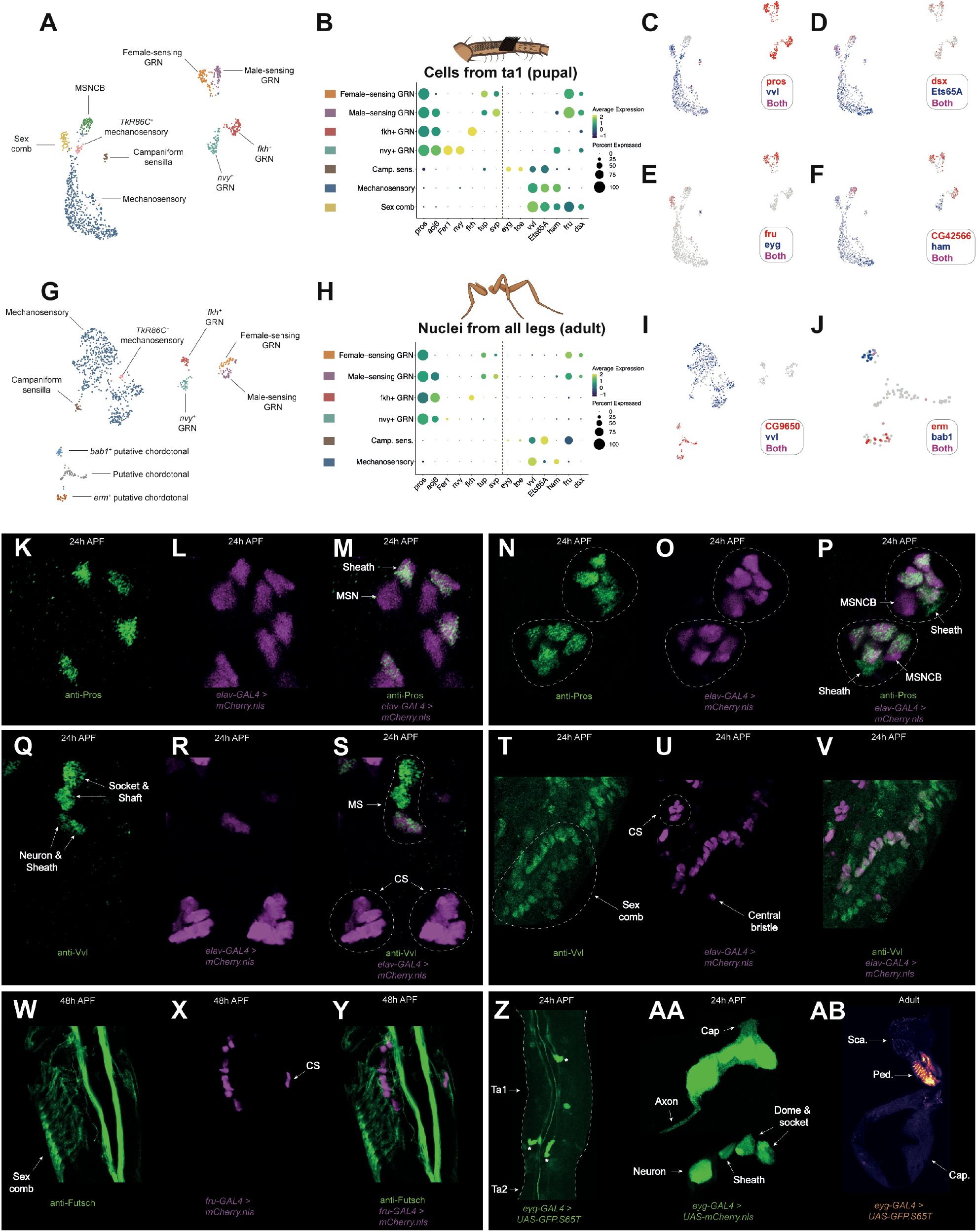
Identification of a combinatorial transcription factor code for leg sensory neurons. (A) Annotated UMAP plot of neuronal cells from the integrated 24h AFP and 30h APF first tarsal segment dataset. GRN = Gustatory Receptor Neuron; MSNCB = mechanosensory neuron in chemosensory bristle. See Figure 6–figure supplement 1A-H for details on the *TkR86C^+^* mechanosensory neurons. (B) Dot plot showing the expression of a series of transcription factors across the clusters labelled in the UMAP given in (A). Each cluster expresses a unique combination. MSNCBs and *TkR86C*^+^ mechanosensory neurons are not shown to focus on major neuron classes. The dotted line separates the chemoreceptor from mechanoreceptor organ neurons. (C-F) UMAP plot of neuronal cells from the integrated 24h AFP and 30h APF first tarsal segment dataset overlaid with the expression of members of the transcription factor code depicted in (B). (C) Note how the MSNCB cluster branching off from the top of the mechanosensory neuron population is negative for both *vvl* and *pros*. (D) *Ets65A* is present in all non-GRN populations in the UMAP, while an effector of sex differentiation, *dsx*, is expressed in GRNs, sex comb neurons, and MSNCBs. (E) *fru*, the other effector of sex differentiation, is enriched in two GRN populations and sex comb neurons, while *eyg* is restricted to campaniform sensilla neurons. (F) *CG42566* is the only non-transcription factor plotted. It is a top marker of MSNCBs and its expression in both MSNCBs and GRNs contributed to this cluster’s chemosensory bristle annotation. *ham* is enriched in mechanosensory neuron classes and two GRN populations. (G) Annotated UMAP plot of male neuronal cells subsetted from the Fly Cell Atlas single-nuclei RNA-seq leg dataset (Li *et al*., 2022). Note the presence of 3 clusters, annotated as ‘putative chordotonal’, that are absent from the pupal dataset – chordotonal organs are not present in the upper tarsal segments. No clear MSNCB or sex comb clusters could be resolved in this dataset. (H) As (B) but for the male neuronal cells subsetted from the Fly Cell Atlas single-nuclei RNA-seq leg dataset. Only those clusters present in the pupal single cell data are shown. (I) UMAP plot of male neuronal cells subsetted from the Fly Cell Atlas single-nuclei RNA-seq leg dataset overlaid with expression of the mechanosensory neuron marker *vvl* (blue) and a top marker of the putative chordotonal organs, the predicted transcription factor *CG9650* (red). (J) A subset of (I), showing only the putative chordotonal clusters overlaid with expression of two transcription factors, *bab1* (blue) and *erm* (red). (K-V) Confocal images of 24h APF male first tarsal segments. (K-M) Mechanosensory bristles from *elav-GAL4 > UAS-mCherry.nls* (magenta) stained with anti-Pros (green). Two *elav-GAL4^+^* cells are present per mechanosensory bristle, one of which, the sheath, is Pros^+^. *elav-GAL4* expression in the sheath is likely due to the legs being imaged soon after the division of the common pIIIb progenitor cell from which they derive (see also Simon *et al*., 2019 and Figure 6–figure supplement 1K-M). MSN= mechanosensory neuron. (N-P) Two chemosensory bristles (circled) from *elav-GAL4 > UAS-mCherry.nls* (magenta) stained with anti-Pros (green). Note that each bristle includes 4 Pros^+^/*elav-GAL4^+^* cells (the gustatory receptor neurons), 1 Pros^+^/*elav-GAL4^−^* cell (the chemosensory sheath cell), and 1 Pros^−^/*elav-GAL4^+^* cell (the MSNCB, mechanosensory neuron in chemosensory bristle). (Q-S) Two chemosensory (CS) and one mechanosensory (MS) bristles from *elav-GAL4 > UAS-mCherry.nls* (magenta) stained with anti-Vvl (green). Note that anti-Vvl staining is entirely absent from the CS bristle including, therefore, the mechanosensory neuron (MSNCB) that innervates it. Conversely, anti-Vvl staining is observed in all 4 constituent cells of a MS bristle. (T-V) The same stainings performed in (Q-S) but centered on the sex comb. Anti-Vvl staining is present in both the neuronal (*elav-GAL4*^+^) and non-neuronal cells of the sex comb. The ‘central bristle’, which develops from the same bristle row as the sex comb is labelled. (W-Y) Confocal images of 48h APF male first tarsal segments showing the expression of *fru-GAL4* (magenta) and anti-Futsch (green). *fru-GAL4* expression is restricted to the sex comb and chemosensory (CS) neurons. The later 48h timepoint was used as *fru-GAL4* was undetectable up until 40h and weak up until 48h. (Z) Confocal image of the first tarsal segment from a 24h male from *eyg-GAL4 > UAS-GFP.S65T*. Campaniform sensilla are marked with asterisks. The axonal projections can be seen as parallel lines running either side of the central autofluorescence. Note that some non-specific fat body staining is also present in this image. (AA) Confocal image of a distal first tarsal segment campaniform sensillum from a 24h male where *eyg-GAL4* is driving the expression of *UAS-mCherry.nls*. The top and bottom image in this panel show the same sensillum but with different levels of saturation to variously highlight the domed structure (top) and the individual cells of the organ (bottom). (AB) As (Z) but showing an adult haltere. Note that the staining is restricted to the campaniform sensilla field on the pedicel (‘Ped.’) and apparently absent from the field on the scabellum (‘Sca.’).

### Unique combinations of the transcription factors *vvl, pros*, and *fru* distinguish between mechanosensory, chemosensory, and sex comb neurons

In both our pupal and the FCA data, the non-overlapping expression of *vvl* and *pros* marked the highest order difference between neuron clusters (Figure 6B,C,H). Among sensory organ cells, *pros* is known to be expressed in sheath cells (Simon *et al*., 2009) and has been detected in DA1, DL3, and VA1d olfactory receptor neurons in scRNA-seq data from antennae (McLaughlin *et al*., 2021). There are also unpublished reports of Pros in leg chemosensory bristle neurons (M. Gho, personal communication). Consistent with sheath cell expression, we observed anti-Pros staining in 1 cell per mechanosensory bristle (Figure 6K). Surprisingly, however, in many cases the Pros^*+*^ cell was also positive for the canonical neuron marker *elav* (Figure 6L,M). We generally observed *elav-GAL4* expression in two cells per bristle, despite mechanosensory bristles being singly innervated (Figure 6L,M). The expression of *elav-GAL4* in the mechanosensory sheath cell likely reflects the early timepoint (24h APF) at which we imaged. This conclusion was also reached by Simon *et al*. (2019) after detecting *elav* expression in sheaths in 28h APF mechanosensory bristles on the notum. Consistent with this conclusion, we observed a heterogenous mix of Pros*^+^/elav-GAL4^+^* and Pros*^+^/elav-GAL4^−^* mechanosensory bristle sheaths in some legs, suggesting between-bristle variation in developmental stage (Figure 6–figure supplement 1K-M). Expression of *elav* beyond the neuron has now been noted several times: its expression appears surprisingly broad and its neural specificity dependent on post-transcriptional repression outside of neurons (Sanfilippo *et al*., 2016; Seroka *et al*., 2022).

Unlike mechanosensory bristles at 24h APF, we did not observe co-staining between anti-Pros and *elav-GAL4* in chemosensory sheath cells – these were all Pros*^+^/elav-GAL4^−^* (Figure 6N-P). This discrepancy between the sheaths of different sensilla classes presumably stems from developmental timing differences as chemosensory bristles are specified earlier than all but the largest mechanosensory bristles (Rodriguez *et al*., 1990; Held, 1990). However, in contrast to mechanosensory bristles, anti-Pros staining in chemosensory bristles was not restricted to the sheath, extending to a further four *elav-GAL4^+^* cells (Figure 6N-P). Mirroring this arrangement, we detected 4 Pros*^+^* populations in both the pupal and FCA data, suggesting that these correspond to four distinct gustatory receptor neuron (GRN) subtypes. Not all *elav-GAL4^+^* cells in the chemosensory bristle were Pros*^+^* though: we observed a single Pros*^-^/elav-GAL4^+^* cell per bristle, which likely corresponds to the MSNCB (mechanosensory neuron in chemosensory bristle; see below). Surprisingly, we failed to detect *Poxn*, which is known to be both necessary and sufficient for a chemosensory rather than mechanosensory fate, in GRNs in either dataset (Bopp *et al*., 1989; Nottebohm, Dambly-Chaudière, & Ghysen, 1992; Nottebohm *et al*., 1994; Dambly-Chaudière *et al*., 1992; Awasaki & Kimura, 1997).

In mechanosensory bristles, anti-Vvl marked all 4 of the constituent cell types (socket, shaft, neuron, and sheath)(Figure 6Q-S). In contrast, no cells in chemosensory bristles were marked by anti-Vvl. Consequently, the mechanosensory neurons innervating mechanosensory and chemosensory bristles can be distinguished based on the presence of Vvl in the former and not the latter. In our pupal scRNA-seq data, we observe a cluster of *vvl^−^* cells branching off from the major *vvl^+^* neuron population (Figure 6C). This cluster is otherwise positive for transcription factors present in the major mechanosensory population, such as *Ets65A* and *ham* (Figure 6D,F), and therefore likely corresponds to MSNCBs. Consistent with this, one of the top markers for the cluster, *CG42566*, was absent from the major mechanosensory population, but expressed in 3 of the 4 GRN populations (Figure 6F). *CG42566* remains largely restricted to GRNs in the adult FCA data (Figure 6–figure supplement 1N). Similar to MSNCBs, we observed an additional cluster branching off from the major mechanosensory population in the pupal data. However, this population was *vvl^+^* and, uniquely among non-chemosensory neurons in the region, heavily enriched for *fru*. Our stainings suggest that these cells correspond to the sex comb neurons, which we observed to be both Vvl^+^ and *fru-GAL4^+^* (Figure 6T-Y; see also Mellert *et al*., 2010 for a previous report of *fru* expression in sex comb neurons). Collectively, these three transcription factors – *pros, fru*, and *vvl* – represent candidate high-level regulators of networks of downstream genes specific to the neurons of three major sensory organ classes: mechanosensory bristles, chemosensory bristles, and the sex comb.

### Campaniform sensilla express the Pax family transcription factors *eyg* and *toe*

In our pupal data, we resolved a small cluster of cells enriched for the Pax family transcription factors *eyg* and *toe*. Expressing *UAS-GFP* under the control of *eyg-GAL4*, we observed staining in the regions of the first tarsal segment corresponding to the positions of the campaniform sensilla (1 proximal organ, 2 distal organs; Dinges *et al*., 2021)(Figure 6Z). Repeating the experiment but with a nuclear-localizing *UAS-mCherry* we detected expression in four cells within the tarsal campaniform sensilla, which presumably correspond to the neuron, sheath, socket, and dome cell (Figure 6AA). Although the expression of *eyg* and *toe* were relatively low in the adult nuclei campaniform sensilla neuron cluster, we observed *eyg-GAL4* activity in both adult legs (Figure 6–figure supplement 1O) and in the adult haltere (Figure 6AA,AB). In the haltere, *eyg-GAL4* activity was detectable in the field of campaniform sensilla on the pedicel, but not the scabellum, raising the possibility that there exist distinct subtypes of campaniform sensilla that express unique gene repertoires.

### The legs contain four gustatory receptor neuron classes, each expressing a unique combination of the transcription factors *acj6, fru, nvy*, and *fkh*

Our stainings showed that among neurons Pros was restricted to chemosensory bristles, indicating that these populations represent subclasses of GRN. Each GRN cluster in our scRNA-seq data expressed a unique combination of 5 transcription factors: *pros^+^*/*acj6^+^*/*nvy^+^, pros^+^*/*acj6^+^*/*fkh^+^*, *pros^+^/acj6^+^/fru^+^*, and *pros^+^/fru^+^* (Figure 7A-F). We validated these combinations using *fru-GAL4* in conjunction with antibodies raised against Pros, Acj6, Nvy, and Fkh (Figure 7G-AA). Multiple bristles are often closely associated within a single region of the leg, while the neurons themselves frequently overlap within a single bristle. Consequently, it wasn’t possible to definitively determine whether one cell of each GRN class was present in every ta1 bristle, but from our observations this seems likely to be the case. Consistent with this, the numbers of cells recovered in each cluster generally appeared similar (Figure 7A). We recovered the same four populations, marked by the same transcription factor code, in the FCA full leg dataset, again recovering a similar number of cells in each GRN population (Figure 7AC). The expression of *nvy* was, however, far less extensive than in the pupal data, suggesting that expression drops off during later pupal development or that *nvy* is restricted to a subset of cells in this GRN class. Correspondence between the *nvy^+^* cluster in the pupal and adult data was supported by additional marker genes, such as *foxo* and *Fer1* (see below). Among neurons, the four GRN populations showed specific or enriched expression of *Ir25a, Ir40a, Gluclalpha, RhoGAP102A, Snmp2, CG42540, CG13578, Tsp47F*, and *CG34342* (Figure 7–figure supplement 1A-T). Taken together, the recovery of the same 4 GRN classes across different timepoints, technologies, and dissected regions suggests that despite substantial between-bristle variation in receptor expression and sensitivity to given stimuli (Ling *et al*., 2014), just 4 core GRN classes are present in the leg and that these classes are defined by combinatorial expression of a small set of transcription factors (Figure 7AB).

**Figure 7.**
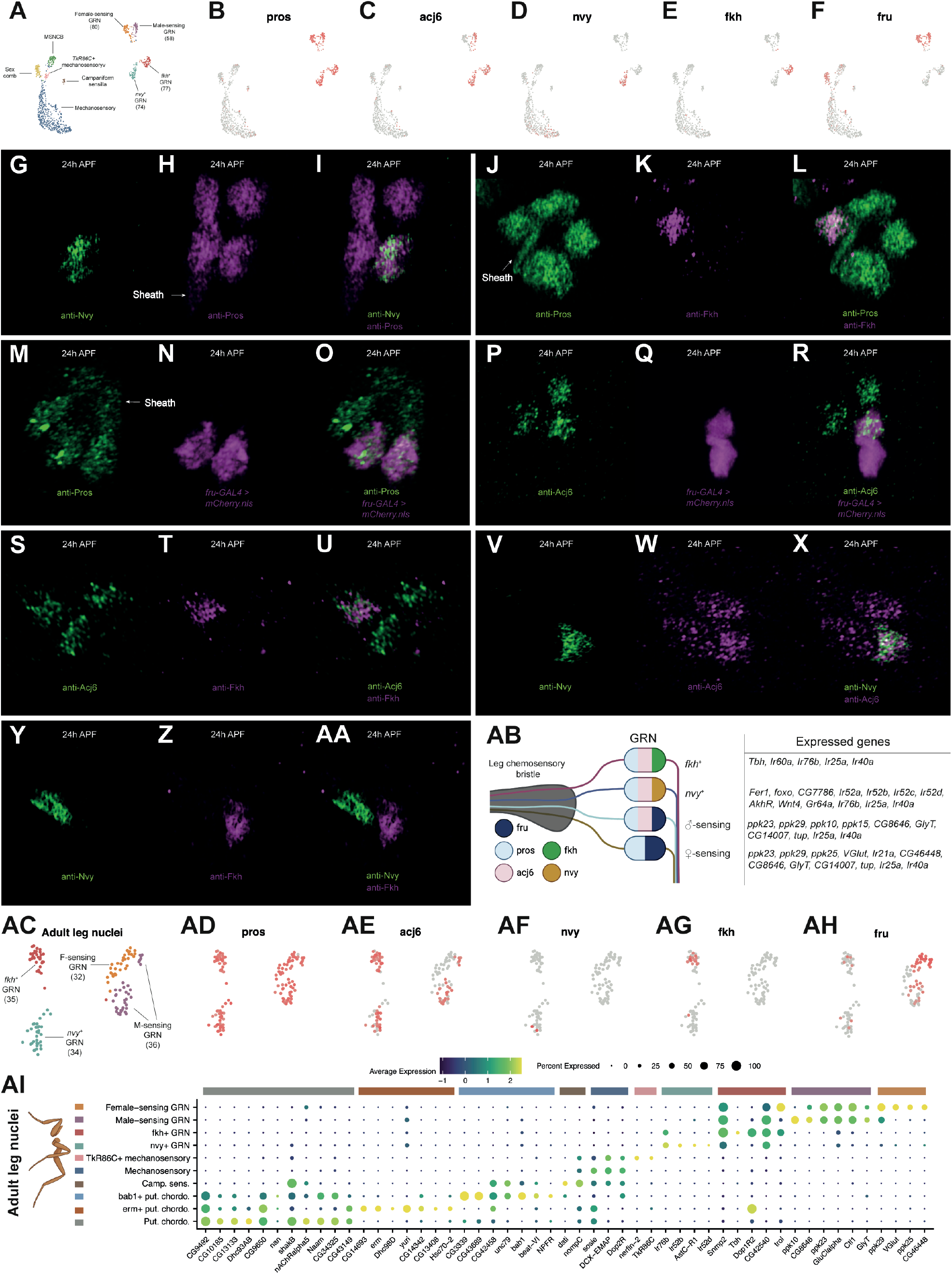
Four gustatory receptor neuron classes express a combinatorial transcription factor code and unique gene repertoires. (A) Annotated UMAP of the pupal integrated neuron data. GRN = gustatory receptor neuron; MSNCB = Mechanosensory neuron in chemosensory bristle. The number of cells in each GRN cluster is presented. The numbers are generally similar between each GRN population, with the exception of the *fru^+^* male-sensing GRNs. This population was closely associated with the *fru^+^* female-sensing GRNs, more so than were any other two GRN subtypes, and the interface between them in UMAP space contained several cells bearing intermediate characteristics. Consequently, the discrepancy in cell numbers between *fru^+^* GRN populations may reflect classification errors due to transcriptomic similarities. (B-F) The UMAP shown in (A) overlaid with the expression of 5 transcription factors (*pros, acj6, nvy, fkh*, and *fru*) that are expressed in unique combinations in each of the 4 GRN clusters. (G-AA) Testing the GRN transcription factor code derived from the scRNA-seq data on 24h APF male first tarsal segments. (G-I) anti-Nvy (green) and anti-Pros (magenta). Note that one Pros^*+*^ cell is partially obscuring the Pros^+^ sheath cell, which has a distinct, elongated morphology. (J-L) anti-Pros (green) and anti-Fkh (magenta). (M-O) anti-Pros (green) and *fru-GAL4 > UAS-mCherry.nls* (magenta). (P-R) anti-Acj6 (green) and *fru-GAL4 > UAS-mCherry.nls* (magenta). (S-U) anti-Acj6 (green) and anti-Fkh (magenta). (V-X) anti-Nvy (green) and anti-Acj6 (magenta). (Y-AA) anti-Nvy (green) and anti-Fkh (magenta). (AB) A schematic summarizing the expression patterns of each transcription factor across GRNs, along with a selection of other genes detected in each subtype. (AC-AH) Recovery of the same transcription factor code in the Fly Cell Atlas single-nuclei adult male leg neuron data. Note that in the adult data, *nvy* is barely detected. Correspondence between the *nvy^+^* cluster in the pupal and adult data was supported by additional marker genes, such as *foxo* and *Fer1* (see Figure 7–figure supplement 2B,C,H,I). (AC) As in (A), the number of cells in each GRN population is presented. In this dataset, a subregion of what unsupervised clustering labelled as the *fru^+^/acj6^−^* population showed *acj6* expression, suggestive of a classification error. This conclusion is further supported by the *VGlut, ppk25*, and *ppk10* data below (see Figure 7-figure supplement 1Y-AB). We therefore manually labelled these as part of the *fru^+^/acj6^+^* cluster. As in the pupal data, the interface between the two *fru^+^* populations appeared particularly close. (AI) A dot plot summarizing the expression of a selection of top differentially expressed genes for each cluster that we identified in the Fly Cell Atlas single-nuclei adult male leg neuron data.

### Male- and female-sensing GRNs express distinct receptor repertoires

Previous work has shown that there are two *fru^+^* GRNs per leg chemosensory bristle (Thistle *et al*., 2012; Kallman *et al*., 2015). Both express the ion channels *ppk23* and *ppk29*, while one additionally expresses *ppk25* and *VGlut* (Lu *et al*., 2012; Starostina *et al*., 2012; Thistle *et al*., 2012; Toda, Zhao, & Dickson, 2012; Vijayan *et al*., 2014; Kallman *et al*., 2015). These *ppk25^+^/VGlut^+^* neurons respond to female pheromones, while the *ppk25^−^/VGlut^−^* cells respond to male pheromones (Kallman *et al*., 2015). In the pupal data, we observe *ppk23* in both *fru^+^* populations, with a small number of *ppk29^+^* cells also distributed across both (Figure 7–figure supplement 1U,V). Although *ppk25* was absent in our dataset, we observed a sharp divide between *fru^+^*/*acj6^+^* and *fru^+^*/*acj6^−^* cells based on *VGlut* expression (Figure 7–figure supplement 1W). *VGlut* is restricted to the *fru^+^*/*acj6^−^* population, identifying these as the female-sensing neurons and the *fru^+^*/*acj6^+^* cells as male-sensing. Receptor expression was more readily detected in the adult nuclei data, suggesting that receptors are generally expressed later in development than the timepoints we sequenced. In this dataset, we again observed that *VGlut* was restricted to the *fru^+^/acj6^−^* population and recovered the previously documented expression patterns of *ppk23, ppk25*, and *ppk29* (Figure 7–figure supplement 1X-AA). However, we also observed GRN-specific expression patterns of several other *ppk* genes. While *ppk25* was restricted to female-sensing cells, *ppk10* and the less frequently detected *ppk15* were restricted to male-sensing cells (Figure 7–figure supplement 1AB-AD). These therefore represent candidate male pheromone receptors. Conversely, *Ir21a* and *CG46448* appeared to be largely restricted to female-sensing neurons (Figure 7–figure supplement 1AE-AF). *CG46448* is adjacent to *VGlut* on chromosome 2L, so their apparent co-expression in female-sensing neurons points to the possibility of their transcriptional control by shared *cis*-regulatory elements. These genes aside, the expression differences we observed between male- and female-sensing neurons were generally limited, but several genes were common to both pheromone-sensing populations, including the transcription factors *tup* and *svp*, the predicted sulfuric ester hydrolase *CG8646, CG14007*, and the glycine transporter *GlyT*, the latter suggesting that these neurons are glycinergic (Frenkel *et al*., 2017)(Figure 7–figure supplement 1AG-AK).

### *nvy^+^* gustatory receptor neurons correspond to a sexually dimorphic population required for normal mating behaviour

The *nvy^+^* GRNs showed specific expression of a further set of transcription factors in the pupal data, namely *Fer1, foxo*, and *CG7786* (Figure 7–figure supplement 2A-D). At these timepoints, few other genes showed enrichment in the *nvy^+^* GRN cluster – exceptions include the phosphodiesterase *Pde6* and G-protein coupled receptor *TrissinR* (Figure 7–figure supplement 2E-F). To further probe the identity of these neurons, we turned to the adult dataset. The transcription factor code identified in the pupal data largely persisted in the adult data, although each was detected in fewer cells and *CG7786* was absent (Figure 7AF; Figure 7–figure supplement 2G-J). *AkhR* and *Gr64a* were among the top markers of the *nvy^+^* GRNs in the adult data (Figure 7–figure supplement 2M,N). But like the other 5 *Gr* genes we detected across the two datasets, *Gr64a* was present in only a handful of cells (Figure 7– figure supplement 2O-T). The sparse detection of *Grs* may stem from gene dropout due to low abundance or reflect their restricted expression among GRNs of the same class. *Ir52a, Ir52b, Ir52c*, and *Ir52d* were also among the top markers of the *nvy^+^* GRN cluster (Figure 7–figure supplement 2U-X). With the exception of *Ir52b*, these ionotropic receptor genes have been well characterized: they are known to be largely coexpressed in a subset of leg taste bristles enriched in the first tarsal segment, to show quantitative differences in expression between males and females, to show sexual dimorphism in their projections (they cross the midline in a commissure in males), to be required for normal sexual behaviour, and to be expressed in neurons distinct from those involved in sweet or bitter sensing (Koh *et al*., 2014; He *et al*., 2019). *Ir52c* and *Ir52d* are further known to be restricted to the forelegs (Koh *et al*., 2014). Despite their sexually dimorphic characteristics, the neurons expressing these receptors are mutually exclusive from those expressing *fru-LEXA* within the same bristle (Koh *et al*., 2014). This accords with the scRNA-seq data, where *fru* appeared largely restricted to the male- and female-sensing populations (Figure 7F,AH). Sexual dimorphism in these neurons is therefore instead likely driven by *dsx*; indeed, at least some *Ir52c^+^* neurons have been shown to descend from a *dsx^+^* lineage (Koh *et al*., 2014). In the adult dataset, *dsx* expression was patchy among GRNs, appearing enriched in the male- and female-sensing populations (Figure 7–figure supplement 2Y). In the pupal data, *dsx* was widely detected across all GRN clusters (Figure 7–figure supplement 2Z). The discrepancy in the extent of *dsx* expression in GRNs between the datasets may reflect the differences in the dissected regions: while most male- and female-pheromone sensing GRNs may be *dsx^+^* regardless of position, it may be that *dsx* expression is restricted among *nvy^+^* and *fkh^+^* neurons to those in the regions of the foreleg more likely to contact a mate than food – *i.e*., the foreleg upper tarsal segments. Restriction of *dsx* expression to a subset of neurons within a GRN class would provide a mechanism through which an additional layer of between-bristle variation in activity could be achieved.

### Distinct and shared modules of gene expression in *nvy^+^* and *fkh^+^* GRNs

The three GRN populations that we have discussed – male-sensing, female-sensing, and *nvy^+^* – match known populations in the literature. However, we were unable to find mention of a population that resembled our fourth, which were *pros^+^/acj6^+^/fkh^+^*. Although none of the top 20 DEGs obtained from a comparison with the 3 other GRN populations showed specific expression in the pupal data, there were intriguing similarities with the *nvy^+^* cluster: *CAH2, jus*, and *Glut4EF* each looked specific to or highly enriched in both (Figure 7–figure supplement 2AA-AC). But in the adult dataset these differences were reduced or lost, suggesting that the variation we observed between GRNs in the pupal data may reflect heterochronic differences (Figure 7–figure supplement 2AD-AF). Nonetheless, the adult dataset presented its own similarities between *fkh^+^* and *nvy^+^* GRNs: expression of *Ir76b*, which is known to be widely expressed among olfactory and gustatory receptor neurons and is thought to form heteromeric complexes with more selectively expressed *Ir’s*, was restricted to *fkh^+^* and *nvy^+^* GRNs (Figure 7–figure supplement 2AG) (Abuin *et al*., 2011; Zhang, Ni, & Montell, 2013; Koh *et al*., 2014; Sánchez-Alcañiz *et al*., 2018). The restricted expression of *Ir76b* contrasts with that of another such co-receptor, *Ir25a*, which we found broadly expressed across all four GRNs (Figure 7–figure supplement 2AH). Although detected in fewer cells, *Ir40a*, which is known to be co-expressed with *Ir25a* in the antennal sacculus, was similarly broadly expressed (Knecht *et al*., 2016) (Figure 7–figure supplement 2AI).

The *fkh^+^* population also had something in common with the female-sensing GRNs: across both datasets, the extracellular matrix proteoglycan gene *trol*, which we also observed in tendon cells (Figure 4–figure supplement 3S-V), was enriched in *fkh^+^* and female-sensing GRNs, showed patchy, low-level expression in male-sensing GRNs, and was essentially absent from *nvy^+^* GRNs (Figure 7– figure supplement 3A-D). We recovered this pattern in *trol-GAL4 > UAS-mCD8∷GFP* (Li *et al*., 2017a) first tarsal segments counter-stained with anti-Pros: of the Pros^+^ cells in a single chemosensory bristle, at least two were strongly *trol-GAL4^+^* and at least two were *trol-GAL4^−^* (including the sheath)(Figure 7–figure supplement 3E-G). In some bristles, the remaining Pros^+^ cell was *trol-GAL4^+^* and in others *trol-GAL4^−^*. *trol* performs several roles during the assembly of the nervous system (Park *et al*., 2003; Cho *et al*., 2012) – why it should be limited in its expression among GRNs is unclear. Alongside these shared modules of gene expression, we identified several uniquely expressed genes in *fkh^+^* GRNs, including *Tbh*, which encodes the key limiting enzyme in octopamine synthesis and therefore suggests these neurons are octopaminergic (Figure 7–figure supplement 2AJ,AK)(Nangia *et al*., 2021). Although very sparsely detected, we also observed *Ir60a* to be limited to *fkh^+^* neurons (Figure 7–figure supplement 2AL).

### Heterochrony in the gene expression profiles of sensory organ neurons

In attempting to identify subtype-specific genes among subclasses of neurons, we observed that genes that appeared cluster-specific in the pupal stages sometimes showed widespread expression in the adult nuclei data. For example, *DCX-EMAP*, a gene that has been implicated in mechanotransduction in both campaniform sensilla and the chordotonal receptors of the Johnston’s organ (Bechstedt *et al*., 2010), switches from being exclusive to campaniform sensilla neurons in the pupal data, to being widespread among both campaniform sensilla and mechanosensory neurons in the adult data (Figure 7–figure supplement 4C,G). Several other genes, such as *nAChRalpha7*, *Ccn, CG1090, CG17839*, and *CG34370*, showed a similar pattern (Figure 7–figure supplement 4A-N). Analogously, MSNCBs and sex comb neurons, identifiable as a *vvl^−^* and *vvl^+^/fru^+^/rn^+^* subpopulation of mechanosensory neurons, respectively, appeared to develop ahead of the major body of mechanosensory neurons that innervate mechanosensory bristles, as evidenced by the expression patterns across datasets of genes such as *sosie, CG31221*, and *dpr13* (Figure 7–figure supplement 4O-T). In these cases, the broadening of expression between pupal and adult datasets points to heterochronic differences (*i.e*., a difference in rate, timing, or duration) in development between the neurons of different sensory organ classes, such that the neurons in mechanosensory bristles lag behind those in chemosensory bristles, campaniform sensilla, and the sex comb. Assuming that the heterochronic differences in the neurons apply similarly to the surrounding sensory support cells, it is conceivable that heterochronic differences between different sensory organ classes contribute to the morphological differences that exist between them. We could imagine, for example, that initiating shaft growth earlier, and thereby lengthening the time frame over which shafts are able to grow, could provide a mechanism underlying the transformation of mechanosensory bristles into sex comb teeth.

### Mechanotransduction neurons from different external sensory organ classes express largely shared gene repertoires

Heterochronic differences between sensory organs complicate the identification of organ-specific genes in the pupal data: genes that appear unique at one stage may be widespread at a later timepoint. For that reason, we initially focused on the adult nuclei dataset to identify genes enriched in specific non-GRN neuron populations (Figure 7AI). The majority of the top mechanosensory neuron markers that were returned from a DGE analysis comparing these cells to all other neurons appeared non-specific, being expressed in one or more other clusters (e.g. *Calx, Fife, Dop2R, KrT95D, CG4577*, and *Ten-m*; Figure 7–figure supplement 5A-M). The same applied to the top campaniform sensilla markers, but in this case, despite their lack of complete specificity, many showed a relatively restricted expression profile that extended across both campaniform sensilla and chordotonal organs (e.g. *dati, unc79, CG42458, TyrR, Cngl, CARPB*, and *beat-VI;* Figure 7–figure supplement 5N-Z). This pattern is suggestive of certain molecular commonalities between campaniform sensilla and chordotonal organ neurons, commonalities that make them distinct from other mechanotransduction neurons. In further pursuit of genes specific to each of mechanosensory neurons and campaniform sensilla, we tried another approach: identifying top markers of each of these clusters in the pupal data and mapping their expression in adults to determine whether their expression remains cluster specific. Of these, the transcription factor *Ets65A*, which in the pupal data was restricted to mechanosensory neurons, sex comb neurons, MSNCBs, and campaniform sensilla neurons, remained restricted to mechanosensory neurons and campaniform sensilla neurons in the adult data (Figure 7–figure supplement 5AA,AB). *Ets65A* therefore represents a candidate regulator of mechanosensory identity in external sensory organ neurons.

We applied this same pupal identification and adult mapping approach in sex comb neurons and MSNCBs, populations that didn’t form distinct clusters in the adult data. No individual gene showed clearly restricted expression to either, but there was strong enrichment of genes that were otherwise patchily expressed across cells. *shakB* was strongly enriched in sex comb neurons relative to other mechanosensory bristles in the pupal data and widely present across putative chordotonal and campaniform sensilla neurons in the adult data (Figure 7–figure supplement 5AC,AD). In MSNCBs, *CG42566*, and to a lesser extent *CG33639*, appeared enriched relative to other mechanosensory populations, with both also detected across GRNs, consistent with a chemosensory bristle origin (Figure 7–figure supplement 5AE-AH). Collectively, MSNCBs and sex comb neurons showed clear enrichment of both effectors of the sex determination pathway: *fru* and *dsx* (Figure 7–figure supplement 5AI-AL). The specificity of this enrichment was greater in the case of *dsx*, which across both neuronal datasets was largely absent outside of the comb, GRN, and MSNCB clusters; *fru* was surprisingly widespread in the adult data, including expression in the putative chordotonal clusters and campaniform sensilla. The expression of these two transcription factors provides a clear regulatory mechanism through which the transcriptomic profiles and activity of these derived mechanosensory populations could readily diverge from other mechanosensory bristles – and do so in a sex-specific manner. But ultimately, determining whether any of the transcriptomic differences between the mechanotransduction neurons that innervate each of mechanosensory bristles, chemosensory bristles, the sex comb, and campaniform sensilla translate into functional differences in operation or sensitivity requires further work. It’s clear from this data that some differences between mechanotransduction neurons innervating different organ classes are present, but they appear minor and on the whole less clear cut than between GRN populations.

### Putative chordotonal organ neuron subtypes express specific gene repertoires

In the FCA adult data, we recovered three putative chordotonal organ neuron clusters that were absent from our first tarsal segment dataset because these organs fall outside the dissected area. We refer to the largest as ‘putative chordotonal’ and the two smaller clusters as *erm^+^* and *bab1^+^* putative chordotonal based on a transcription factor they were each specifically enriched for. Many of the genes we identified as specifically enriched in the putative chordotonal organs in our DGE analysis were absent from the pupal dataset, consistent both with their organ-specificity and the absence of chordotonal organs from our dataset. Along with the genes shared among campaniform sensilla neurons and chordotonal organ neurons discussed in the previous section (Figure 7–figure supplement 5N-Z), we identified genes specific to or highly enriched in all chordotonal populations (Figure 7– figure supplement 6A-K), as well as those specific to subpopulations (Figure 7–figure supplement 6L-V). The transcriptomic distinctiveness we detect between putative chordotonal neuron clusters aligns with previous work that has identified multiple, functionally distinct neurons in a single chordotonal organ (e.g. ‘Type A’ and ‘Type B’ neurons; Mamiya *et al*., 2018; McKelvey *et al*., 2021). The next step will be to match the distinct clusters we recover to these different chordotonal neuron classes. In turn, that would raise secondary questions, such as whether each neuron class is present in each of the several chordotonal organs that are housed within the leg.

### The support cells within a mechanosensory organ each express a distinct gene repertoire

Campaniform sensilla, chemosensory bristles, and mechanosensory bristles each consist of four distinct cell types generated through asymmetric divisions of a sensory organ precursor (SOP). The first division gives rise to a progenitor of the socket (tormogen) and shaft (trichogen) cells, while the other to a progenitor of the sheath (thecogen) and neuron(s) (reviewed in Fichelson *et al*., 2005)(Figure 8A). Few marker genes are known for the non-neuronal SOP-descendants and, to the best of our knowledge, none that definitively separate the same cell type between different organs (e.g. mechanosensory vs chemosensory sockets) (Prelic *et al*., 2022). Those markers that are known include: *Su(H)* and *Sox15*, which specifically accumulate in socket cell nuclei (Gho *et al*., 1996; Miller *et al*., 2009); *sv* (*Pax2*), which although initially expressed in all bristle cells during the mitotic phase of development is eventually restricted to the shaft and sheath (Kavaler *et al*., 1999); *nompA*, which is specifically expressed in sheath cells where it is required to connect dendrites to the shaft (Chung *et al*., 2001); and *pros*, which is expressed in sheath cells (Figure 6K-P; Simon *et al*., 2019).

**Figure 8.**
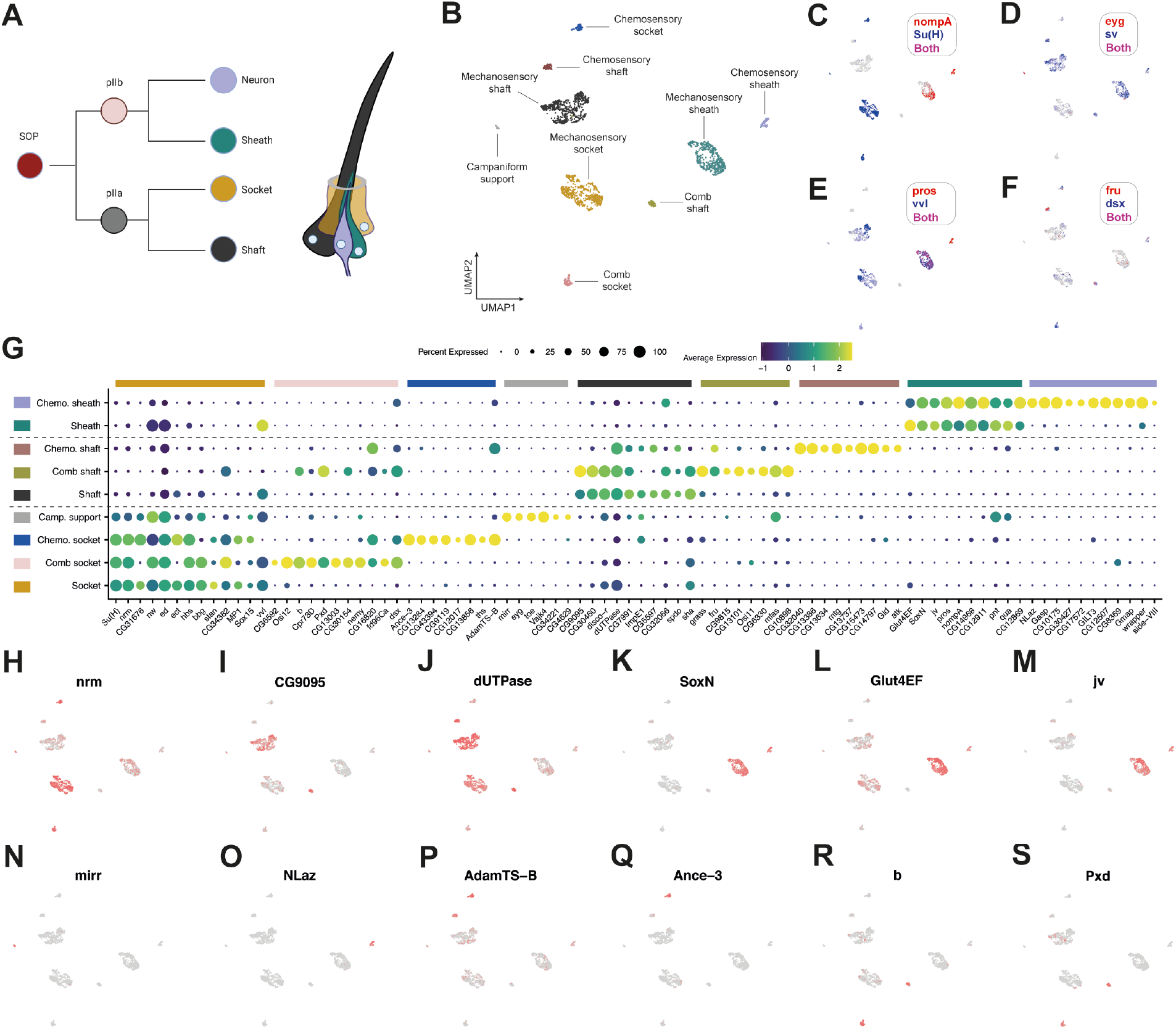
Distinct and shared modules of gene expression between sensory organ support cells. (A) The four constituent cell types of external sensory organs, such as the mechanosensory bristle in this schematic, originate through asymmetric divisions of a sensory organ precursor (SOP) cell (reviewed in Fichelson *et al*., 2005). The SOP divides to produce a pIIa and pIIb daughter cell. pIIa further divides to generate a socket and shaft cell. In the notum, where it’s been studied, pIIb divides into a pIIIb cell and glial cell, the latter of which enters apoptosis soon after birth (Fichelson & Gho, 2003). pIIIb further divides to produce the sheath and neuron. To the best of our knowledge, whether the pIIb glial division occurs in the leg remains untested. (B) Annotated UMAP plot of the bristle cells from the integrated 24h AFP and 30h APF first tarsal segment dataset. The campaniform support cluster included *Su(H)^+^* cells, which suggests it corresponds to socket cells, but it’s possible that it includes a mix of campaniform sensilla accessory cells. (C-F) The UMAP shown in (B) overlaid with the expression of a series of marker genes, either previously published or demonstrated in this study, for different sensilla classes or accessory cell types. (G) A dot plot summarizing the expression patterns of a selection of genes identified as being differentially expressed in each of the clusters given in the UMAP shown in (B). Dotted lines separate the three major classes of sensory support cell. Of the socket markers, *CG31676* is known to be expressed in a subset of olfactory projection neurons (Li *et al*., 2017a); *nw* is a C-type lectin-like gene; *stan*, a cadherin that controls planar cell polarity (Usui *et al*., 1999); and *nrm, ed*, and *hbs* are cell adhesion molecule genes (Kania *et al*., 1993; Kania & Bellen, 1995; Wei *et al*., 2005; Bao & Cagan, 2005; Grillo-Hill & Wolff, 2009). Of the shaft markers, *CG9095* encodes a C-type lectin-like gene; *disco-r* encodes a transcription factor; *dUTPase* encodes a nucleoside triphosphate; *spdo*, encodes a transmembrane domain containing protein that regulates Notch signaling during asymmetric cell division (O’Connor-Giles & Skeath, 2003; Babaoglan *et al*., 2009; Upadhyay *et al*., 2013); and *sha*, which encodes a protein involved in the formation of bristle hairs (Ren *et al*., 2006). Aside from *pros* and *nompA*, the top markers of the sheaths include: the transcription factors *Glut4EF, pnt*, and *SoxN*; *jv*, which encodes a protein involved in actin organization during bristle growth (Shapira *et al*., 2011); *qua*, which encodes an F-actin cross-linking protein (Mahajan-Miklos & Cooley, 1994); and the midline glia marker *wrapper*, which encodes a protein involved in axon ensheathment (Noordermeer *et al*., 1998; Wheeler *et al*., 2009; Stork *et al*., 2009). (H-S) The UMAP shown in (B) overlaid with the expression of genes identified in this study as markers of sensory organ support cell subtypes.

Based on these markers, we identified major shaft, socket, and sheath clusters in our data (Figure 8B-F). The large size of each of these relative to other clusters in the support cell dataset indicates that they belong to the dominant sensory organ in the first tarsal segment: mechanosensory bristles. Further evidence for a mechanosensory origin comes from the observation that each of these clusters was *vvl*^+^, which we previously observed to be expressed in all mechanosensory, but no chemosensory, bristle cells (Figure 6Q-S). Among the DEGs we identified for each of the mechanosensory shaft, socket, and sheath clusters were many that reflected the biology of these sensory support cells (Figure 8G-M): the cell adhesion molecule genes *nrm, ed*, and *hbs* (Kania *et al*., 1993; Kania & Bellen, 1995; Wei *et al*., 2005; Bao & Cagan, 2005; Grillo-Hill & Wolff, 2009) in socket cells; *sha*, which encodes a protein involved in the formation of bristle hairs (Ren *et al*., 2006), in shaft cells; and, in the sheath, a duo of genes, *qua* and *jv*, involved in the organization of actin during bristle growth (Mahajan-Miklos & Cooley, 1994; Shapira *et al*., 2011). Another nod to the biology of the sheath came in its enrichment for *wrapper*, which encodes a protein known to be involved in axon ensheathment (Noordermeer *et al*., 1998). Classically used as a marker of midline glia (Noordermeer *et al*., 1998; Wheeler *et al*., 2009; Stork *et al*., 2009), the expression of *wrapper* in sheaths reinforces their glia-like properties, despite not expressing *repo* or *gcm*. Moreover, it points to general, shared elements in the mechanisms of ensheathment of neuronal processes between these cell types. The enriched expression in sheaths of a trio of transcription factors – *Glut4EF, pnt*, and *SoxN*, of which the latter was completely restricted to sheaths in our dataset – provide a potential route to regulatory divergence from other support cells and glia.

### Homologous support cells show transcriptomic divergence between sensory organ classes

We observed that homologous support cells in different sensory organ classes express both shared and distinct gene repertoires. The top markers for each of the mechanosensory socket, shaft, and sheath clusters were enriched in a set of smaller clusters: three clusters showed socket-like profiles, 2 shaft-like profiles, and 1 sheath-like profile (Figure 8G). These minor clusters therefore appear to be sensory organ cell subtypes from sensilla classes that are less abundant in the first tarsal segment than are mechanosensory bristles. Our visualization of *eyg-GAL4 > UAS-GFP* revealed *eyg-GAL4* expression in all 4 cells in a campaniform sensillum (Figure 6AA), so the presence of *eyg* and *toe* in the smallest cluster suggest these cells correspond to a campaniform sensilla population. This cluster uniquely expressed the transcription factor *mirr* (Figure 8N) and the CPLCP cuticle protein family gene *Vajk4*, and was *Su(H)^+^*, suggesting a socket cell identity, but no further *eyg^+^/toe^+^* clusters were present. Several explanations for why are plausible: (a) this cluster contains a mix of all campaniform sensilla support cells; (b) only the sockets are transcriptomically distinct enough to cluster separately; and (c) only the sockets were recovered in sufficient numbers to cluster separately. That a small, co-clustering group of *eyg^+^* sheath cells were present in the mechanosensory sheath population, rather than forming their own distinct cluster, provides some support for (b) and (c) (Figure 8D). Ultimately, the presence of *eyg* and *toe* across the constituent cells of campaniform sensilla suggest that these genes may be master regulators of campaniform sensillum identity.

The minor sheath population was enriched for *CG42566*, a gene we previously found to be specific to MSNCBs and GRNs among the neurons (Figure 7–figure supplement 5AE,AF), suggesting that this cluster represents descendant cells of the chemosensory pIIb lineage and, therefore, the chemosensory sheath cells specifically. As well as several poorly characterized genes, this cluster showed enriched or unique expression of the lipid-binding protein encoding gene *NLaz* (Figure 8O), the chitin-binding protein encoding gene *Gasp*, and *Side-VIII*. As we observed with neurons, at least some of the differences between mechanosensory and chemosensory sheaths appear to be heterochronic, with chemosensory sheaths developing ahead of their mechanosensory homologues: several genes, including *nompA* and *wrapper*, were widely detected among chemosensory sheath cells, but in mechanosensory sheaths showed localized expression in a region enriched for cells from the 30h dataset (Figure 8–figure supplement 1A-J).

Based on the expression of *Su(H), Sox15*, and *sv*, the remaining clusters have apparent socket and shaft identities. Given the representation of different sensilla classes in the first tarsal segment the most likely classification for the *vvl^−^* clusters are chemosensory sockets and shafts. Except for the extracellular protease gene *AdamTS-B*, which was heavily enriched in both clusters (Figure 8P), there was no clear transcriptomic link between them. Among the top markers for each, the *sv^+^* putative shaft cluster showed strong enrichment for *mtg*, which encodes a chitin binding domain-containing protein that’s required to drive postsynaptic assembly (Rushton, Rohrbough, & Broadie, 2009), while the *Su(H)^+^*/*Sox15^+^* putative socket cluster showed unique expression of *CG43394* (Figure 8–figure supplement 2) and *Ance-3* (Figure 8Q), which encodes a predicted membrane component orthologous to human ACE2, the receptor for SARS-CoV-2 (Zhou *et al*., 2020).

### Sex comb cells are enriched for the expression of melanogenic pathway genes

Unlike the chemosensory support cells, the remaining minor putative shaft and socket clusters shared many of their top markers with one another. These genes included: *b* (Figure 8R) and *Pxd* (Figure 8S), components of the melanogenic pathway (Phillips *et al*., 2005; Rahman *et al*., 2021); *dsx*, the effector of the sex determination pathway (reviewed in Hopkins & Kopp, 2021); and *fd96Ca*, a gene known to contribute to sex comb formation (Ruiz-Losada, Pérez-Reyes, & Estella, 2021) and which we found to be one of the top markers of sex comb neurons in the pupal data (data not shown). These two clusters therefore likely correspond to sex comb shafts and sockets. Whether the genes we find enriched in sex comb sockets and shafts (Figure 8G) are unique to these cells is hard to determine from these data alone due to heterochronic differences between mechanosensory bristles and the sex comb, the latter of which develops earlier. This difference is reflected in the expression of *vvl*, which was substantially reduced in the putative sex comb shaft cluster relative to both the sex comb socket and the mechanosensory populations (Figure 8E,G). Previous work has shown that while Vvl is present in all four SOP-descendent cells in external sense organs on the head and notum at 24h APF, by ~42h it’s restricted to the socket (Inbal, Levanon, & Salzberg, 2003). Despite the heterochronic differences that likely exist, the enrichment of melanogenic pathway genes is consistent with the combs heavily melanized appearance. Future work is required to determine whether the changes to mechanosensory bristle cells that generate the modified sex comb morphology operate primarily through quantitative, qualitative, or heterochronic changes in gene expression.

## Discussion

Sensory perception begins with contact between a stimulus and a specialized sensory organ. Within and between animal species, these organs are highly variable in their form, cellular composition, and molecular characteristics. To understand the genetic underpinnings of this diversity, both in terms of the developmental networks that specify organ development and the molecular profiles that define mature organs, we characterized the transcriptomes of cells in a region of the male *Drosophila* foreleg that carries a structurally and functionally diverse selection of sensory organs.

### Hierarchical and combinatorial transcription factor codes for leg sensory neurons

The discrete identities adopted by cell types depend on cell type-specific expression of transcription factors, which function as regulators of downstream networks of gene expression. We identified a combinatorial transcription factor code unique to each sensory neuron population present in the first tarsal segment (Figure 9). The ‘first order’ differences in transcription factor expression defined neurons involved in mechanosensation (*i.e*., those innervating mechanosensory bristles, the sex comb, and campaniform sensilla, along with the MSNCB), which expressed *Ets65A*, and neurons involved in chemosensation, which expressed *pros*. The ‘second order’ differences defined subtypes within each of these major classes: the expression of *eyg* and *toe* separated campaniform sensilla from other mechanotransduction neurons; *acj6, fru, nvy*, and *fkh* delineated GRN subtypes; and *vvl* separated the mechanosensory neurons of mechanotransduction organs from those that innervate chemosensory bristles. Of these transcription factors, *acj6*, and to a lesser extent *fkh*, have repeatedly cropped up in studies of *Drosophila* neurons, including in subpopulations within the visual, auditory, and olfactory systems (Davis *et al*., 2020; Allen *et al*., 2020; Xie *et al*., 2021; McLaughlin *et al*., 2021; Özel *et al*., 2021; Janssens *et al*., 2022). The expression of these transcription factors in a subset of functionally varied neurons points to shared regulatory architecture between them and, by extension, perhaps to common downstream networks of gene expression.

**Figure 9.**
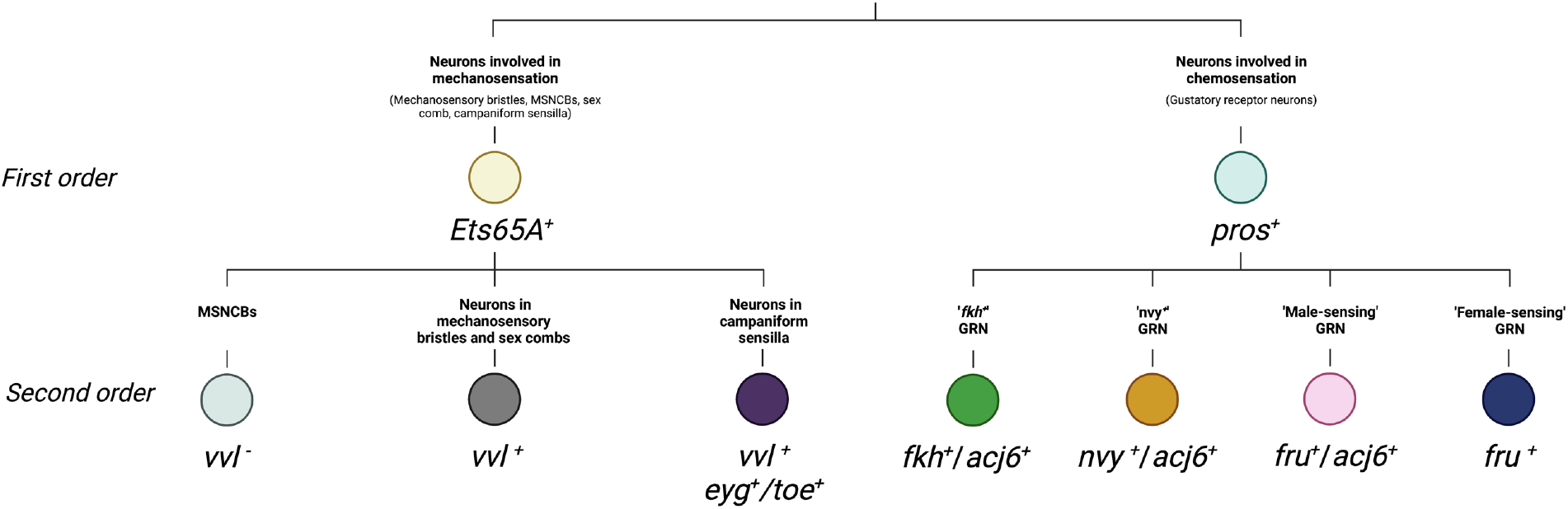
First and second order differences in transcription factor expression between sensory neuron classes. MSNCB = mechanosensory neuron in chemosensory bristle. GRN = gustatory receptor neuron.

### Orthogonal regulatory input from the sex differentiation pathway?

The effectors of the sex determination pathway, *fru* and *dsx*, showed restricted expression across neuron populations. Two GRNs expressed *fru*, as did a subpopulation of mechanosensory neurons innervating the sex comb. *dsx*, on the other hand, was expressed in all GRN populations, sex comb neurons, and in the subpopulation of mechanosensory neurons that innervate chemosensory bristles (MSNCBs). However, the prevalence of *fru* and *dsx* expression differed among neuron classes. *fru* was widely detected in female- and male-sensing GRNs and the sex comb neurons, but was only present in a few *fkh^+^* and *nvy*^+^ GRNs. In the pupal first tarsal segment data, *dsx* was detected across all GRNs, although more patchily in *fkh^+^* neurons than in the other subtypes. In the adult GRNs, *dsx* was more widely detected among female- and male-sensing GRN clusters with a very small number of *dsx^+^* cells present in the remaining GRN populations. The patchiness of expression may reflect gene dropout, with differences in patchiness between populations stemming from quantitative differences in expression. Another explanation, however, is that in some neuron classes, *fru* and *dsx* expression is limited to subsets of cells. In this case, sexual identity would serve as a regulatory input orthogonal to the neuron’s core genetic identity and, by extension, provide a regulatory route to sexually dimorphic gene expression in a subset of neurons within a shared class. The orthogonal nature of this regulatory input is most likely to manifest itself in the *dsx^+^* or *fru^+^* subsets of *fkh^+^* and *nvy^+^* GRNs, where *dsx* and *fru* expression appears to be relatively rare. In contrast, in the male- and female-pheromone sensing GRNs or the sex comb, *fru* expression may form a core part of that cell population’s identity.

### Reconciling the functional diversity of gustatory receptor neurons with their limited transcriptomic diversity

Considerable diversity is known to exist among leg chemosensory taste bristles in the sensitivities and responsiveness they show to a wide range of tastants, as well as in the identities of the receptors they express (Bray & Amrein, 2003; Starostina *et al*., 2012; Koh *et al*., 2014; Ling *et al*., 2014). But in all datasets we analyzed here – two from a highly localized region of the developing foreleg and one from the entire length of each of the three pairs of adult legs – we resolved only the same four GRN populations. How, then, can we reconcile the functional diversity known to exist between leg taste bristles with the limited number of transcriptomically distinct GRN classes that we identified? There are several possible solutions. First, a member of each GRN class might not be present in every bristle on the leg, but rather members of a class may be restricted to subsets of bristles. As each chemosensory bristle on the tarsal segments is thought to be innervated by four GRNs (Nayak & Singh, 1983), this would require that some bristles are innervated by multiple GRNs of the same class. Second, it may be that there are additional functionally and transcriptomically distinct GRN classes in the leg, but that they are rare in comparison to the four classes we resolved. A very small number of taste bristles at the tips of the leg are known to not be innervated by a *ppk25*^+^ female-sensing GRN (Starostina *et al*., 2012). This same region also houses some of the least numerous taste bristle classes – classes that are more likely to contact food than potential mates – that have been identified on the basis of morphology and response to tastants (Ling *et al*., 2014). Bristles in this region of the leg would naturally not have made it into our first tarsal segment datasets and their rarity in the wider leg may have limited their recovery in the adult dataset.

There is a third solution to the disconnect between the known functional diversity and transcriptomic similarity that we recover among GRN populations. It might be that the leg taste bristles are largely – or in every case – innervated by a minimal number of four functionally distinct GRN classes, each defined by a unique combination of transcription factors. But on top of this core transcriptomic program may be layered an additional level of regulation, one that we were unable to resolve here, that enables the same GRN class in different bristles to express a different receptor or membrane channel repertoire. There is evidence for this in the datasets we looked at here. *Ir52c* and *Ir52d*, two genes expressed in our *nvy^+^* population, have been shown by both RT-PCR and GAL4s to be exclusively expressed in the forelegs, where they are coexpressed (Koh *et al*., 2014). In a dataset composed of all three pairs of legs, like the adult nuclei dataset we analyzed, *Ir52c^+^/Ir52d^+^* cells should therefore comprise a very small number of the total recovered cells, their numbers having been diluted by GRNs from bristles on the other legs. This is indeed what we see, but rather than forming their own cluster we find the *Ir52c^+^/Ir52d^+^* cells alongside a greater number that are negative for these two genes in a cluster that also includes cells expressing *Ir52a*, which is known to be expressed in bristles on all legs (Koh *et al*., 2014). Expression patterns such as this, where cells expressing a different set of receptors cocluster, along with the similar representation of cells from each GRN class in our datasets suggest that a ‘minimal GRN class’ model seems likely to underlie most, if not all, taste bristles on the leg. The nature of the extra regulatory layer that allows receptors to be swapped in and out of a core GRN class remains unclear. The positions of the bristles in which some receptors show restricted expression appear stereotyped (Bray & Amrein, 2003; Ling *et al*., 2014), which suggests that additional layers of regulation may involve positional information. It also suggests that any role for stochasticity in receptor expression, as occurs in mammalian olfactory neurons (Magklara & Lomvardas, 2013), might be limited. Finally, restricting the expression of the effectors of sex determination to subsets of bristles would allow for regulatory divergence in neurons from a common GRN class between bristles.

Whatever the differences between the neurons of a single GRN class innervating different bristles may be, our data suggest that the identity of a GRN is more than just the receptors it expresses; that there are deeper transcriptomic features that define a common GRN class. The defining features of a GRN class may relate to the neurotransmitters through which they relay the detection of stimuli or, perhaps, the complement of Ionotropic Receptor (IR) co-receptors they express. On this point, some IRs, including IR25a and IR76b, are known to act as co-factors, forming heteromeric complexes with more selectively expressed IRs (Abuin *et al*., 2011; Zhang *et al*., 2013; Koh *et al*., 2014; Sánchez-Alcañiz *et al*., 2018). We found these co-receptors to vary in their specificity, *Ir25a* being broadly expressed across all four GRNs and *Ir76b* restricted to *fkh^+^* and *nvy^+^* GRNs. We could imagine a modular configuration where co-receptors could set the broad functional range that defines a GRN class – *i.e*., those classes we recover in our single cell data – while additional gene products, differentially expressed between neurons of a given GRN class in different bristles, refine GRN function to generate functional diversity between bristles.

Three of the four GRN classes we identify match the descriptions of functional populations that have been reported in the literature, each of which is involved in the detection of conspecifics. Two of these are the *fru^+^*/*ppk23^+^* ‘female-pheromone-sensing’ and ‘male-pheromone-sensing’ neurons (also known as ‘F cells’ and ‘M cells’), a pair of neurons that respond selectively to the pheromones of the sex they’re named after (Kallman *et al*., 2015). Of the genes we identify as specifically expressed in these cells, *ppk10* and *ppk15* stand out as candidate male pheromone detectors, given their restriction to male-sensing neurons and the known roles of other *ppk* genes in pheromone-sensing (e.g. Lu *et al*., 2012; Starostina *et al*., 2012; Thistle *et al*., 2012; Toda *et al*., 2012; Vijayan *et al*., 2014). The other identifiable GRN population that we resolved was the *nvy^+^*, *Ir52a,c,d*-expressing neurons (Koh *et al*., 2014). These neurons are activated by exposure to conspecific females, but not males, and promote courtship. In this sense, naming *fru^+^*/*ppk23^+^*/*ppk25^+^* GRNs as ‘female-pheromone-sensing neurons’ – a name used by others (Kallman *et al*., 2015) and by us throughout this paper – fails to recognize that these neurons can sit alongside a different GRN with apparently the same broad function in the same bristle. The existence of two distinct classes of female-sensing neurons raises the question of why two should be required. That we find *ppk* genes to be limited to one of the two female-sensing GRNs points to differences in either what the neurons respond to or how they respond. A large number of putative pheromones exist in female *Drosophila*, both in *D. melanogaster* and in other species (Everaerts *et al*., 2010; Khallaf *et al*., 2021). This diversity may necessitate multiple detector neurons in order to accurately discriminate between sex and species, and perhaps to even allow males to perform finer-grain assessments of the condition or mating status of potential partners. Future work will be required to determine whether responsiveness to these different compounds is divided between the two GRN classes, whether there is a logic to that division based on the identity of those compounds, and whether these GRNs project into different neural circuits. Finally, that leaves the one GRN population – the *fkh^+^* population – that we couldn’t identify. Whether this class also contributes to conspecific detection and to what extent its function may vary between bristles remains unclear.

### The not-so-specific expression of glia markers

Three glia cell types have been described in the *Drosophila* peripheral nervous system: (1) wrapping glia, which ensheath individual or bundled axons to support the rapid conductance of action potentials; (2) perineural glia, which form the outer cell layer of the nervous system, positioned below a dense network of extracellular matrix called the neural lamella, and which are thought to be responsible for nutrient uptake via the contacts they make with hemolymph; and (3) subperineural glia, large cells that form a thin layer beneath the perineural glia and which establish septate junctions with one another to provide an important structural component of the blood-brain barrier (reviewed in Stork *et al*., 2008; Bittern *et al*., 2021). All three of these glia are known to be present in the leg (Sasse & Klämbt, 2016) and a number of marker genes are known for each (Schwabe *et al*., 2005; Mayer *et al*., 2009; Xie & Auld, 2011; DeSalvo *et al*., 2014; Lassetter *et al*., 2021). But the glia populations we resolved did not fall neatly along those marker gene lines. *moody* and to a lesser extent *Gli*, which are both used as specific markers of subperineural glia, were widely detected across our *repo^+^* cells, as were the wrapping glia markers *nrv2* and *Ntan1* (Xie & Auld, 2011; Sasse & Klämbt, 2016). *Jupiter*, which is thought to specifically mark perineural glia (Xie & Auld, 2011), was even more widely detected, being expressed in all non-sensory cells. The non-canonical expression patterns we observe in our glia populations with respect to known marker genes raise difficult questions. Have the GAL4s and protein traps we’ve relied on given us a misleading impression of glia type-specific patterns of gene expression and protein localization? Perhaps – it’s clear that for at least some of these markers antibodies label a wider set of cells than do drivers, as in the case of the *apt-GAL4 GMR49G07* (Sasse & Klämbt, 2016). But the surprisingly broad expression of known marker genes is also consistent with an emerging pattern in single cell studies, where expression of many genes is considerably wider than expected from reporter and antibody staining data (e.g. Seroka *et al*., 2022). This phenomenon illustrates the importance of cell type-specific post-transcriptional regulation in enforcing the cell type specificity of gene expression, as has been shown for surprisingly widely expressed genes such as the neuronal marker *elav* (Sanfilippo *et al*., 2016). After all, mRNA levels are not the final output of gene expression (Buccitelli & Selbach, 2020).

### A novel cell type and the construction of the neural lamella

We identified a population that didn’t seem to match the description of any reported in the literature. These cells, enriched for the expression of the transcription factors *Sox100B* and *Lim1*, exhibited several glia-like properties. For one, they expressed the midline glia marker *wrapper* (Noordermeer *et al*., 1998; Wheeler *et al*., 2009; Stork *et al*., 2009). More directly, our stainings showed a glia-like association between these cells and the axon trunks that project from the leg sensory organs to the ventral nerve cord. Enriched expression of *beaten path* genes, *beat-IIa* and *beat-IIb*, was further suggestive of a cell that’s in direct communication with others in the developing nervous system (Li *et al*., 2017b). But from a glia perspective, these cells pose a problem: they’re negative for the canonical glia marker *repo*, a transcription factor expressed in all glia except for those of the midline, and they appear to originate within the leg disc itself, rather than migrating in along with CNS-derived glia. We have three clues for what these cells might be. First, a homolog of *Sox100B, Sox10*, is involved in the specification of glia cells that play similar roles to *Drosophila* wrapping glia in the vertebrate nervous system (oligodendrocytes and Schwann cells)(Finzsch *et al*., 2010; Turnescu *et al*., 2018; Fogarty, Kitzman, & Antonellis, 2020; Saur *et al*., 2021; Ittner *et al*., 2021). Indeed, *Drosophila* Sox100B can rescue vertebrate Schwann cell development in the absence of Sox10 (Cossais *et al*., 2010). The failure of *Sox100B^+^* cells to express the wrapping glia marker *Oaz* would argue against these cells corresponding to wrapping glia, as would their failure to express *repo*. It may therefore be the case that a conserved developmental program drives wrapping-type morphologies in both these cells and the vertebrate glia subtypes. Which leads us to the second clue: while the expression of membrane-bound GFP revealed narrow cell morphologies when driven by *repo-GAL4*, under the control of *Lim1-GAL4* the staining appeared to encircle a larger area, extending more laterally. The *Lim1^+^/Sox100B^+^* cells therefore appear to comprise the outer most layer of the nervous tissue. Which leads us to the third clue: these cells were enriched for *vkg*, a subunit of Collagen IV that’s known to label the neural lamella (Stork *et al*., 2008; Xie & Auld, 2011). Collectively, therefore, these data suggest that the *Sox100B^+^* cells are a novel, axon-associated cell type that’s required for the construction of the neural lamella.

Our description of *Sox100B^+^* cells challenges the idea that the source of the neural lamella is the perineural glia (Edwards, Swales, & Bate, 1993; see also Xie & Auld, 2011 who fail to find an effect of RNAi knockdown of *vkg* on the neural lamella using glia drivers). A more recent alternative hypothesis for the formation of the neural lamella is that its major components – including Collagen IV – are deposited by migrating hemocytes during embryogenesis (Xie & Auld, 2011). This idea is based on the finding that the failure of hemocytes to migrate adjacent to the developing ventral nerve cord is associated with the failure of Collagen IV and another extracellular matrix component, Peroxidasin, to be deposited around it (Olofsson & Page, 2005). Two things are interesting to note here. The first is that with the possible exception of *NimC3, Sox100B^+^* cells bear no clear transcriptomic similarity to hemocytes. The second is that hemocytes and *Sox100B^+^* cells show partially overlapping expression of many extracellular matrix component genes. It’s therefore possible that both hemocytes and *Sox100B*^+^ cells are collectively required for the construction of the neural lamella, each contributing a subset of extracellular matrix components. Resolving the specific role played by *Sox100B*^+^ cells in this process, as well as how labour may be divided between additional cell types, represents a key focus for future work.

## Materials and methods

### Fly strains and husbandry

Flies used in all experiments were raised on a standard cornmeal medium and housed in an incubator at 25°C on a 12:12 cycle. Lines used in this study are detailed in the Key Resources Table.

### Isolation of first tarsal segments for single-cell sequencing

The following procedure was used to collect known-age first tarsal segments for single-cell sequencing. White P1 prepupae (0-1 hour After Puparium Formation, APF) from the DGRP line *RAL-517* (Mackay *et al*., 2012) were collected and sexed based on the presence of testes. Males were transferred to a folded kimwipe, wet with 500μl of water and held inside of a petri dish in an incubator maintained at 25°C on a 12:12 cycle. 1h before the desired age was reached (e.g. 23h after collection for the 24h sample), pupae were removed from puparia using forceps and placed on top of a water-soaked kimwipe. When the desired age was reached, pupae were placed ventral side up on tape. The base of the abdomen was pierced to release some of the fluid pressure and the foreleg removed at the tibia/tarsal joint. The dissected leg was then placed on tape, covered in a drop of 1X Dulbecco’s PBS (DPBS, Sigma, D8537), and a scalpel was used to sever at approximately the mid-point of the second tarsal segment. The first tarsal segment was then eased out of the pupal cuticle and transferred to a glass well on ice containing 100μl of 1X DPBS using a BSA-coated 10μl tip.

### Single-cell suspension preparation

We separately prepared two single-cell suspensions, each composed of 67 first tarsal segments collected following the procedures described above: one from 24h APF male pupae, the other from 30h APF male pupae. For a given timepoint, once 67 first tarsal segments had been collected, the DPBS was removed from the well and replaced with 100μl of dissociation buffer, which consisted of 10X TrypLE (Thermo Fisher Scientific, A12177-01) with a final concentration of 2μl/mL of collagenase (Sigma, C0130). The well was sealed and submerged in a metal bead bath in an incubator at 37°C for 35 minutes. The dissociation buffer was then removed from the well and replaced with 50μl of room temperature DPBS before the tissues were subjected to mechanical dissociation. For this, the solution was pipetted up and down 20 times using a 200μl, widebore, low bind, freshly BSA coated tip (Thermo Fisher Scientific, 2069G) on a 50μl pipette set to 40μl. The tip was fully submerged to avoid bringing air into the suspension. The solution was then pipetted up and down a further 20x using a flame-rounded, BSA-coated 200μl tip, again on a 50μl pipette set to 40μl. Next, the solution was pipetted up and down slowly 3 times using the same flame-rounded tip before 40μl was taken up and transferred to a 2mL, low-bind, wide-bottomed tube on ice. 20μl of DPBS was then slowly dripped around the edges of the well using a different tip to flush the cells into the center. This was then pipetted up and down 3 times using the original flame-rounded 200μl tip and added to the tube. Cell concentration and viability was assayed using an acridine orange/propidium iodide stain and measured using a LUNA-FL Fluorescent Cell Counter averaged across 2 x 5μl aliquots (24h APF sample: 1990 cells/μl, 98% viability; 30h APF sample: 967 cells/μl, 98% viability). Through this approach, we were able to avoid filtration or centrifugation steps, both of which we observed to result in high cell loss in trial runs. Although we didn’t perform a controlled test of it, we also observed that the viability of our cell suspensions was considerably higher when using DPBS compared to PBS.

### Library preparation and sequencing

Barcoded 3’ single cell libraries were prepared from single cell suspensions using the Chromium Next GEM Single Cell 3’ kit v3.1 (10X Genomics, Pleasanton, California) for sequencing according to the recommendations of the manufacturer. The cDNA and library fragment size distribution were verified on a Bioanalyzer 2100 (Agilent, Santa Clara, CA). The libraries were quantified by fluorometry on a Qubit instrument (LifeTechnologies, Carlsbad, CA) and by qPCR with a Kapa Library Quant kit (Kapa Biosystems-Roche) prior to sequencing. The libraries were sequenced on a NovaSeq 6000 sequencer (Illumina, San Diego, CA) with paired-end 150 bp reads. The sequencing generated approximately 50,000 reads per cell and 500 million reads per library.

### scRNA-seq data processing

The alignment, barcode assignment, and UMI counting of the two 10x leg samples were performed using the ‘count’ function in CellRanger (v4.0.0). The reference index was built using the ‘mkref’ function in CellRanger (v3.1.0) and the Ensembl BDGP6.28 *Drosophila melanogaster* genome. The GTF was filtered to remove non-polyA transcripts that overlap with protein-coding gene models, as recommended in the CellRanger tutorial (https://support.10xgenomics.com/single-cell-gene-expression/software/pipelines/latest/using/tutorial_mr). Cell quality filtering and downstream analysis was performed using Seurat (v4.0.5) (Satija *et al*., 2015; Butler *et al*., 2018; Stuart *et al*., 2019; Hao *et al*., 2021) in R (v4.1.1). We began by creating a SeuratObject from the count data using the function ‘CreateSeuratObject’, excluding cells where fewer than 100 genes were detected. Based on the distribution of the genes and transcripts detected per cell, along with the percentage of reads that mapped to mitochondria, we used the following cell-level filters: 24h dataset, >450 genes/cell, <5000 genes/cell, >2500 transcripts/cell, <10% mitochondrial reads/cell; 30h dataset, >425 genes/cell, <5000 genes/cell, >1400 transcripts/cell, <10% mitochondrial reads. This led to the removal of 1141 (leaving 9877) and 1486 (leaving 10332) cells from the 24h dataset and 30h dataset, respectively. A median of 2083 genes and 11292 transcripts were detected per cell in the resulting 24h dataset, but only 1245 genes and 5050 transcripts in the 30h dataset. This discrepancy is due at least in part to the substantially reduced % of reads mapped confidently to the transcriptome in the 30h sample (24.0% vs 81.1%), the cause of which we were unable to identify but which did not obviously impact the downstream clustering.

### scRNA-seq data clustering

Next, we applied gene-level filtering to the 24h and 30h datasets, retaining only those genes expressed in 3 or more cells, and used the function ‘SCTransform’ implemented in Seurat (v4.0.5) to normalize and scale the full 24h and 30h datasets (Hafemeister & Satija, 2019). We used 5000 variable features and regressed out variation due to the percentage of mitochondrial reads. UMAPs of the full datasets were constructed using the ‘FindNeighbors’, ‘FindClusters’, and ‘RunUMAP’ functions using principal components (PCs) 1 to 100 and a clustering resolution of 0.7. To identify doublets produced by stochastic encapsulation of multiple cells within a single bead during the microfluidic process, we used the R package DoubletFinder (v2.0.3) (McGinnis, Murrow, & Gartner, 2019). We observed that when run on the full 24h dataset, DoubletFinder heavily targeted a subregion of the neuron cluster (~29% of all doublets were in the mechanosensory neuron cluster; Figure 10A). Compared to other cells in the neuron cluster, the identified doublets were heavily enriched for the mitotic marker *string* and cell adhesion gene *klingon*, the latter of which was restricted to the doublet-enriched region of the mechanosensory neuron cluster as well as a subset of GRNs and mechanosensory sheaths (Figure 10B,C). Similar targeting of neurons and sheaths by DoubletFinder was observed in the 30h dataset (Figure 10L). Given this distribution, we reasoned that when run on the entire dataset, DoubletFinder was classifying early differentiating neurons and sheaths, which share a progenitor cell, as doublets. This may be due to their displaying an intermediate transcriptional profile. We therefore ran DoubletFinder separately on the epithelial and non-epithelial datasets. To generate these, we subsetted our data into epithelial cells, identifiable as the large body of contiguous clusters in each dataset (circled in Figures 10A,L), and all remaining clusters before reapplying gene-level filtering (retaining those genes expressed in 3 or more cells), re-running SCTransform (regressing out variation due to the percentage of mitochondrial reads), and reclustering (hereafter all three of these processes will be included under the term ‘reclustering’)(Parameters: 24h and 30h datasets: epithelial: PCs = 1:150, variable features = 5000, r = 0.7; non-epithelial: PCs = 1:50, variable features = 5000, r = 0.7).

**Figure 10.**
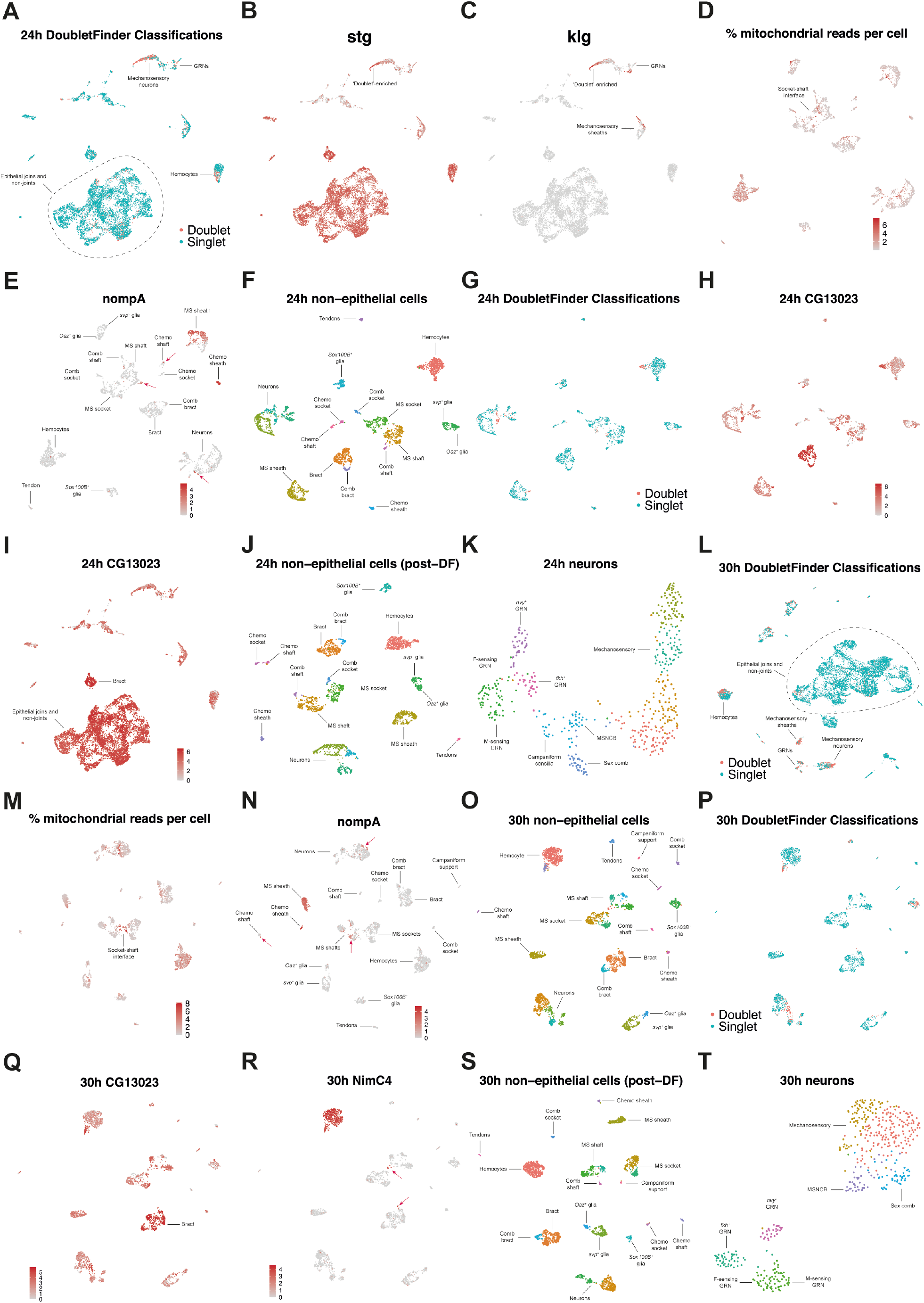
Processing the 24h AFP and 30h APF male first tarsal segment scRNA-seq datasets. (A) A UMAP plot of the 24h dataset after low quality cells have been removed (*i.e*. those that fail to meet the criteria of >450 genes/cell, <5000 genes/cell, >2500 transcripts/cell, <10% mitochondrial reads/cell). Putative doublets identified by DoubletFinder are coloured red and singlets blue. Note how in this data, Doublets are heavily enriched within a subregion of mechanosensory neurons and hemocytes. ~29% of cells in the mechanosensory neuron cluster were labelled as doublets. (B) The UMAP plot shown in (A) overlaid with the expression of the mitotic marker *stg*. Note how the same region of the mechanosensory neuron cluster that is enriched for ‘doublets’ is also enriched for *stg*. (C) The UMAP plot shown in (A) overlaid with the expression of *klg*, a gene we identified as one of the top markers of the ‘doublets’ when compared to other mechanosensory cells in the cluster. The only other regions of the UMAP where *klg* expression is detected is in closely associated subsets of mechanosensory sheaths and GRNs. By itself, this restricted expression pattern suggests that these cells are not *bona fide* doublets. Rather, and considered alongside the *stg* expression, it suggests that these cells are early differentiating neurons. Given that neurons and sheaths are formed from the same cell division, it further suggests that *klg* is an early marker of both cell types. (D) The percentage of reads per cell that map to mitochondrial genes overlaid on a UMAP of all the 24h non-epithelial cells. Note the presence of high % cells at the interface between the socket and shaft clusters. (E) The expression of the sheath marker *nompA* overlaid on a UMAP of all the 24h non-epithelial cells. Note the presence of *nompA^+^* cells in other sensory support cell clusters (shafts and neurons; see pink arrows). The presence of *nompA* is likely indicative of doublets arising through incomplete dissociation of these tightly associated cells within sensory bristles. ‘MS’ = mechanosensory. (F) A UMAP plot of all 24h non-epithelial cells after removal of sensory support cells with >2 *nompA* transcripts detected or where more than 5% of reads mapped to mitochondrial genes. (G) The UMAP shown in (F) with putative doublets identified by DoubletFinder coloured red and singlets blue. When run on this subsetted data, the identified doublets are more evenly dispersed among clusters than when run on the full dataset (as shown in A). (H) The UMAP shown in (F) overlaid with the expression of *CG13023*, one of the top markers of the doublets identified in the neuron cluster. *CG13023* is enriched in all areas of the UMAP where doublets were identified as well as in the bract clusters. Bracts develop from epithelial cells which suggests that the doublets identified are likely to be *bona fide* doublets that include an epithelial cell. (I) As (H) but overlaid on the full 24h UMAP shown in (A). *CG13023* is enriched in epithelial cells, further supporting the conclusion that the *CG13023* enriched cells identified by DoubletFinder are *bona fide* doublets. (J) A UMAP plot of the 24h non-epithelial cells clustered after the removal of doublets identified in (G). (K) A UMAP plot of the neurons subsetted from (J) and reclustered. (L) A UMAP plot of the 30h dataset, post cell-filtering, with putative doublets identified by DoubletFinder coloured red and singlets blue. As in the 24h data, doublets are heavily enriched within a subregion of mechanosensory neurons and hemocytes. (M) A UMAP plot of the subsetted non-epithelial cells overlaid with the percentage of reads per cell that map to mitochondrial genes. Note the presence of high % cells at the interface between the socket and shaft clusters. (N) The expression of the sheath marker *nompA* overlaid on a UMAP of all the 30h non-epithelial cells. As in the 24h dataset, note the presence of *nompA^+^* cells in other sensory support cell clusters (shafts and neurons; see pink arrows). (O) A UMAP plot of all 30h non-epithelial cells after removal of sensory support cells with >2 *nompA* transcripts detected or where more than 5% of reads mapped to mitochondrial genes. (P) The UMAP shown in (O) with putative doublets identified by DoubletFinder coloured red and singlets blue. As with the 24h data, when run on the non-epithelial subset of cells, the identified doublets are more evenly dispersed among clusters than when run on the full dataset (as shown in (L)). (Q) As in the 24h data, cells identified as doublets are enriched for *CG13023*, a gene that shows high expression in epithelial cells, from which bracts derive. The full 30h UMAP shows an equivalent pattern to that observed in the full 24h UMAP in (I) (data not shown). (R) As shown in (P), DoubletFinder identified several doublets at the interface between the mechanosensory shafts and socket clusters, as well as on the periphery of the bracts. Closer inspection of other cells in this region (see arrows) revealed that many of the neighboring cells identified as singlets expressed high levels of the hemocyte marker NimC4, suggesting that they too were doublets. These cells were manually highlighted and removed. (S) A UMAP plot of the 30h non-epithelial cells clustered after the removal of doublets identified in (P). (T) A UMAP plot of the neurons subsetted from (S) and reclustered. No campaniform sensilla (*i.e. eyg^+^/toe^+^* cells) neurons are readily detected in this UMAP.

At this stage the non-epithelial cell datasets contained 2919 (24h) and 3163 (30h) cells. We then further lowered the mitochondrial percent threshold to <5%, which removed a further 15 (24h) and 38 (30h) cells, and non-sheath sensory support cells with >2 *nompA* transcripts (24h: 44 cells; 30h: 96 cells; Figure 10D,E,M,N). *nompA* is a known sheath marker (Chung *et al*., 2001) and its presence in cells in other sensory support cell clusters indicates possible doublets arising from incomplete dissociation. This left a total of 2860 (24h) and 3029 (30h) cells, which we reclustered (Figure 10F,O)(Parameters: PCs = 1:50, variable features = 5000, r = 0.7). We then ran DoubletFinder, using the estimated doublet rates outlined in the 10x V3 user guide for the number of cells we recovered (24h: 8.4%, ~11,000 cells; 30h: 9.3% ~11,800 cells; parameters: 1:50 PCs, pN = 0.25, pK = 0.05, without homotypic adjustment). In both datasets, the identified doublets were more evenly dispersed among clusters compared to when run on the full dataset (Figure 10G,P). Moreover, genes enriched in the identified doublets were non-specific and often enriched in epithelial and bract (an identity induced in epithelial cells) clusters (*e.g. CG13023*)(Figure 10H,I,Q), suggestive of them being doublets containing epithelial cells. Cells classified as doublets were removed (217 in 24h; 259 in 30h). In the 30h dataset, a further 9 cells that were enriched for the hemocyte marker *NimC4* were removed as presumed doublets from the bract cluster and from the interface between the mechanosensory shaft and socket clusters (Figure 10R). Each dataset was then reclustered (Figure 10J,S)(Parameters 24h: PCs = 1:35, variable features = 5000, r = 0.7; 30h: PCs = 1:30, variable features = 5000, r = 0.7). The 24h and 30h non-epithelial datasets were then integrated using the ‘SelectIntegrationFeatures’ (nfeatures = 2000), ‘prepSCTIntegration’, ‘RunPCA’, ‘FindIntegrationAnchors’ (normalization method = SCT, dims = 1:35, reduction = rpca, k anchor =5), and ‘IntegrateData’ functions in Seurat. Non-sensory cells (clusters enriched for *NimC4^+^*, *repo^+^*, *Sox100B^+^*, *aos^+^*, and *drm^+^*/*vvl^+^*) and sensory support cells (clusters enriched for *nompA^+^*, *Su(H)^+^*, and *sv^+^*) were then separately subsetted and reclustered for downstream analysis (non-sensory: PCs = 1:25, r = 0.4; bristle: PCs = 22, r = 0.2). We identified neurons separately in the non-integrated datasets based on *fne* expression, reclustered them (Parameters 24h and 30h: PCs = 1:30, var features = 3000, r = 1.5; 24h neuron dataset = 514 cells; 30h neuron dataset = 473 cells; Figure 10K,T), and integrated them separately (same parameters as for the integration detailed above but with k anchor of 20).

For epithelial cells, we ran DoubletFinder and separately reclustered the two datasets (Parameters 24h: PCs = 1:150, variable features = 5000, r = 0.7, with homotypic adjustment, 6374 cells; 30h: PCs = 1:30, variable features = 5000, r = 0.7, with homotypic adjustment, 6502 cells). We then integrated the two datasets using the approach outlined above (Parameters: nfeatures = 2000, normalization method = SCT, dims = 1:35, reduction = rpca, k anchor =5), additionally regressing out variation due to cell cycle stage – specifically, the difference between the G2M and S phase scores (calculated following the steps in the Seurat tutorial; https://satijalab.org/seurat/archive/v3.0/cell_cycle_vignette.html). The integrated data were then clustered (PCs = 1:30, r = 0.6), the joints removed and clustered separately (PCs = 10, r = 0.2), and the remaining non-joint cells clustered (PCs = 20, r = 0.4).

### scRNA-seq data analysis

Differential gene expression analyses were conducted on the normalized count data. Data was normalized using the Seurat function ‘NormalizeData’ with normalization method ‘LogNormalize’, where counts for each cell are divided by the total counts for that cell and multiplied by a scale factor (the default of 10000) and then natural-log transformed using log1p. To perform the differential gene analysis, we used a Wilcoxon rank sum test implemented through the ‘FindConservedMarkers’ function, grouping cells by their dataset of origin (24h or 30h) and specifying that genes were only tested if they were present in >10% of the cells in the focal population. Dot plots and plots of gene expression overlaid on UMAPs were generated using the ‘DotPlot’ and ‘FeaturePlot’ functions, respectively.

### Fly Cell Atlas data

To assess the robustness of our developmental neuron classifications in a different, adult dataset, we analyzed the neuronal populations from the Fly Cell Atlas leg dataset (Li *et al*., 2022). This dataset differed in several ways from those we’ve generated here: ours is single cell, while the FCA data is single nuclei; ours is pupal (24h APF and 30h APF), while the FCA data is from adults; ours is from the first tarsal segment of the foreleg, while the FCA data is from the full length of all three pairs of legs. We downloaded the ‘10x stringent’ leg loom data (https://cloud.flycellatlas.org/index.php/s/ZX56j2CcMXnHXYc), from which we extracted the gene expression matrix using the ‘open_loom’ and ‘get_dgem’ functions in the ‘SCopeLoomR’ package (https://github.com/aertslab/SCopeLoomR/). We created a SeuratObject using this data, filtering out genes expressed in <3 cells and cells with <100 genes. The dataset was clustered using the functions described above (PCs = 50, r = 1.7) and neurons identified based on *fne* expression. We then extracted male neurons from this integrated, mixed-sex dataset, which were identifiable from the cell metadata, and reclustered (variable features = 3000, PCs = 1:25, r = 1).

### Fixation, immunohistochemistry, and microscopy

White P1 prepupae (0-1 hour After Puparium Formation, APF) were collected and sexed under light microscope based on the presence of testes. Males were then placed on a damp kimwipe and aged in an incubator at 25°C until the desired timepoint. Pupae were then placed on their side on sticky tape and a razor blade used to cut away the dorsal half. The cut pupae were then fixed in a 4% paraformaldehyde solution (125μl 32% PFA, 675μl H2O, 200μl 5X TN) for 50 minutes on a rotator at room temperature and stored at 4°C. For leg dissections, fixed pupae were removed from the puparia in 1X TNT and tears made in the pupal cuticle at the femur-tibia boundary using forceps. The tibia through to ta5 region was then pulled through the tear, freeing it from the pupal cuticle. At this point, legs expressing fluorescent proteins were mounted in Fluoromount 50 (SouthernBiotech). For immunohistochemistry, the dissected leg region was blocked with 5% goat serum (200μl 10% goat serum with 200μl 1X TNT) overnight at 4°C. Legs were then incubated with primary antibody solution overnight at 4°C, washed for 21 minutes 4 times in 1X TNT, incubated with the secondary antibody solution for 2 hours at room temperature, washed for 21 minutes 4 times in 1X TNT, and then mounted in Fluoromount 50 (SouthernBiotech). All stages, from dissection through to staining, were carried out in a glass well. Antibodies were used in solution with 1X TN and a final concentration of 2% goat serum (see Key Resource Table for identities and concentrations). Confocal images were taken using an Olympus FV1000 laser scanning confocal microscope or a Zeiss 980 Airyscan. Image stacks were processed using Z-series projection in ImageJ.

## Data and code availability

Raw sequencing reads and preprocessed sequence data are available through NCBI GEO with accession code GSE215073. All code used in this paper is available on the Open Science Framework with DOI 10.17605/OSF.IO/BA8TF. Processed Seurat objects for each dataset (including all subsetted, annotated datasets) used in this study are available at the same address.

## Acknowledgements

We thank Diana Burkart-Waco, Emily Kumimoto, and Hong Qiu for preparing libraries from our single cell suspensions and performing the sequencing; John Larue, Aidan Angus-Henry, and Tiezheng Fan for assistance with collecting, fixing, and mounting pupae; Kohtaro Tanaka for assistance in developing the first tarsal segment dissection technique; Alex Majane, Evan Witt, and Allison Jevitt for answering questions that helped us design our tissue dissociation protocol; Michael Paddy and Thomas Wilkop for microscopy assistance; and Michel Gho for sharing unpublished data on Pros expression. Thanks also to Liqun Luo, Cedric Soler, Makoto Sato, David Bilder, Hui-Yu Ku, and the Bloomington Stock Center for *Drosophila* strains, and to the Developmental Studies Hybridoma Bank, Steve Russell, Sarah Certel, Vidya Chandrasekaran, Herbert Jackle, Richard Mann, Marc Freeman, and Megan Corty for antibodies. We would like to acknowledge the use of a Zeiss 980 confocal microscope made available through an NIH grant S10OD026702. The sequencing was carried out by the DNA Technologies and Expression Analysis Core at the UC Davis Genome Center, supported by NIH Shared Instrumentation Grant 1S10OD010786-01. This work was supported by a long-term fellowship from the Human Frontier Science Program Organization (LT000123/2020-L) to B.R.H. and grant R35 GM122592 to A.K.

## Author contributions

B.R.H.: Conceptualization, Methodology, Software, Validation, Formal analysis, Investigation, Data curation, Writing – original draft preparation, Writing – review and editing, Visualization, Funding acquisition.

O.B.: Methodology, Validation, Investigation.

A.K.: Conceptualization, Methodology, Data curation, Writing – review and editing, Funding acquisition, Resources, Project administration, Project supervision.

---

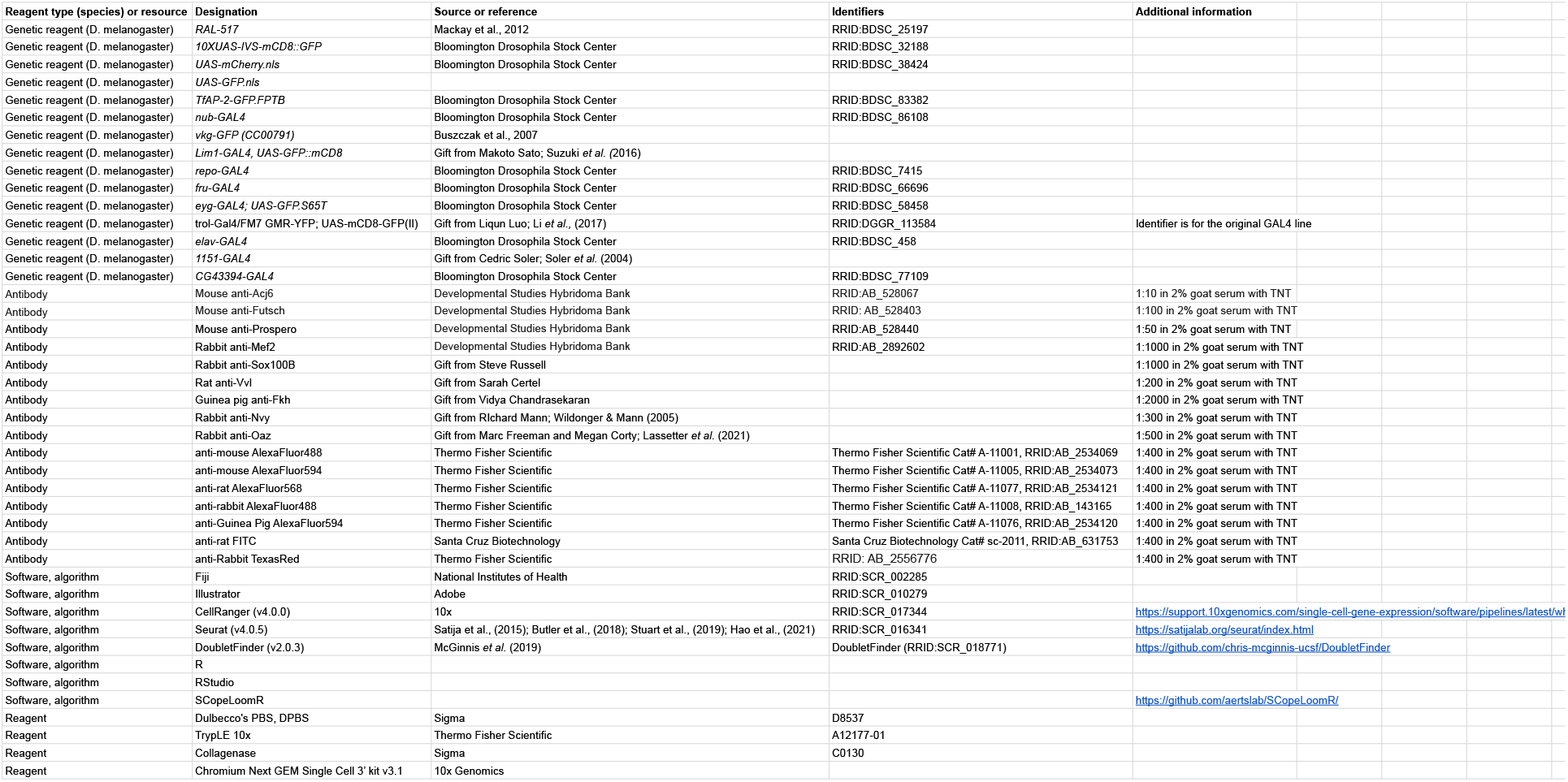
Key resource table.

## References

Abuin, L., Bargeton, B., Ulbrich, M.H., Isacoff, E.Y., Kellenberger, S. & Benton, R. (2011) Functional Architecture of Olfactory Ionotropic Glutamate Receptors. Neuron 69, 44–60. Neuron.

Akay, T., Haehn, S., Schmitz, J. & Büschges, A. (2004) Signals from load sensors underlie interjoint coordination during stepping movements of the stick insect leg. Journal of Neurophysiology 92, 42–51. American Physiological Society.

Del álamo, D., Terriente, J. & Díaz-Benjumea, F.J. (2002) Spitz/EGFr signalling via the Ras/MAPK pathway mediates the induction of bract cells in Drosophila legs. Development 129, 1975–1982. The Company of Biologists.

Allen, A.M., Neville, M.C., Birtles, S., Croset, V., Treiber, C.D., Waddell, S., Goodwin, S.F. & Mann, R.S. (2020) A single-cell transcriptomic atlas of the adult Drosophila ventral nerve cord. eLife 9. eLife Sciences Publications Ltd.

Almanzar, N., Antony, J., Baghel, A.S., Bakerman, I., Bansal, I., Barres, B.A., Beachy, P.A., Berdnik, D., Bilen, B., Brownfield, D., Cain, C., Chan, C.K.F., Chen, M.B., Clarke, M.F., Conley, S.D., ET AL. (2020) A single-cell transcriptomic atlas characterizes ageing tissues in the mouse. Nature 583, 590–595. Nature Publishing Group.

Armand, P., Knapp, A.C., Hirsch, A.J., Wieschaus, E.F. & Cole, M.D. (1994) A novel basic helix-loop-helix protein is expressed in muscle attachment sites of the Drosophila epidermis. Molecular and Cellular Biology 14, 4145–4154. American Society for Microbiology.

Awasaki, T. & Kimura, K.I. (1997) pox-neuro is required for development of chemosensory bristles in Drosophila. Journal of Neurobiology 32, 707–721.

Babaoglan, A.B., O’Connor-Giles, K.M., Mistry, H., Schickedanz, A., Wilson, B.A. & Skeath, J.B. (2009) Sanpodo: A context-dependent activator and inhibitor of Notch signaling during asymmetric divisions. Development 136, 4089–4098. The Company of Biologists.

Bakken, T.E., Hodge, R.D., Miller, J.A., Yao, Z., Nguyen, T.N., Aevermann, B., Barkan, E., Bertagnolli, D., Casper, T., Dee, N., Garren, E., Goldy, J., Graybuck, L.T., Kroll, M., Lasken, R.S., ET AL. (2018) Single-nucleus and single-cell transcriptomes compared in matched cortical cell types. PLoS ONE 13.

Bao, S. & Cagan, R. (2005) Preferential adhesion mediated by hibris and roughest regulates morphogenesis and patterning in the drosophila eye. Developmental Cell 8, 925–935. Cell Press.

Basil, M.C., Cardenas-Diaz, F.L., Kathiriya, J.J., Morley, M.P., Carl, J., Brumwell, A.N., Katzen, J., Slovik, K.J., Babu, A., Zhou, S., Kremp, M.M., Mccauley, K.B., Li, S., Planer, J.D., Hussain, S.S., ET AL. (2022) Human distal airways contain a multipotent secretory cell that can regenerate alveoli. Nature 604, 120–126. Nature Publishing Group.

Bechstedt, S., Albert, J.T., Kreil, D.P., Müller-Reichert, T., Göpfert, M.C. & Howard, J. (2010) A doublecortin containing microtubule-associated protein is implicated in mechanotransduction in Drosophila sensory cilia. Nature Communications 1.

Beckervordersandforth, R.M., Rickert, C., Altenhein, B. & Technau, G.M. (2008) Subtypes of glial cells in the Drosophila embryonic ventral nerve cord as related to lineage and gene expression. Mechanisms of Development 125, 542–557.

Bittern, J., Pogodalla, N., Ohm, H., Brüser, L., Kottmeier, R., Schirmeier, S. & Klämbt, C. (2021) Neuron–glia interaction in the Drosophila nervous system. Developmental Neurobiology 81, 438–452. John Wiley & Sons, Ltd.

Bomidi, C., Robertson, M., Coarfa, C., Estes, M.K. & Blutt, S.E. (2021) Single-cell sequencing of rotavirus-infected intestinal epithelium reveals cell-type specific epithelial repair and tuft cell infection. Proceedings of the National Academy of Sciences of the United States of America 118. National Academy of Sciences.

Bopp, D., Jamet, E., Baumgartner, S., Burri, M. & Noll, M. (1989) Isolation of two tissue-specific Drosophila paired box genes, Pox meso and Pox neuro. EMBO Journal 8, 3447–3457. John Wiley & Sons, Ltd.

Bray, S. & Amrein, H. (2003) A Putative Drosophila Pheromone Receptor Expressed in Male-Specific Taste Neurons Is Required for Efficient Courtship. Neuron 39, 1019–1029. Cell Press.

Buccitelli, C. & Selbach, M. (2020) mRNAs, proteins and the emerging principles of gene expression control. Nature Reviews Genetics 21, 630–644. Nature Publishing Group.

Butler, A., Hoffman, P., Smibert, P., Papalexi, E. & Satija, R. (2018) Integrating single-cell transcriptomic data across different conditions, technologies, and species. Nature Biotechnology.

Cao, J., Packer, J.S., Ramani, V., Cusanovich, D.A., Huynh, C., Daza, R., Qiu, X., Lee, C., Furlan, S.N., Steemers, F.J., Adey, A., Waterston, R.H., Trapnell, C. & Shendure, J. (2017) Comprehensive single-cell transcriptional profiling of a multicellular organism. Science 357, 661–667. American Association for the Advancement of Science.

Chanana, B., Graf, R., Koledachkina, T., Pflanz, R. & VorbrÜGgen, G. (2007) αPS2 integrin-mediated muscle attachment in Drosophila requires the ECM protein Thrombospondin. Mechanisms of Development 124, 463–475.

Chari, T., Weissbourd, B., Gehring, J., Ferraioli, A., Leclère, L., Herl, M., Gao, F., Chevalier, S., Copley, R.R., Houliston, E., Anderson, D.J. & Pachter, L. (2021) Whole-animal multiplexed single-cell RNA-seq reveals transcriptional shifts across Clytia medusa cell types. Science Advances 7, 1683. American Association for the Advancement of Science.

Cho, J.Y., Chak, K., Andreone, B.J., Wooley, J.R. & Kolodkin, A.L. (2012) The extracellular matrix proteoglycan perlecan facilitates transmembrane semaphorin-mediated repulsive guidance. Genes and Development 26, 2222–2235.

Chung, Y.D., Zhu, J., Han, Y.G. & Kernan, M.J. (2001) nompA Encodes a PNS-Specific, ZP Domain Protein Required to Connect Mechanosensory Dendrites to Sensory Structures. Neuron 29, 415–428. Cell Press.

Cossais, F., Sock, E., Hornig, J., Schreiner, S., Kellerer, S., Bösl, M.R., Russell, S. & Wegner, M. (2010) Replacement of mouse Sox10 by the Drosophila ortholog Sox100B provides evidence for co-option of SoxE proteins into vertebrate-specific gene-regulatory networks through altered expression. Developmental Biology 341, 267–281. Dev Biol.

Croset, V., Treiber, C.D. & Waddell, S. (2018) Cellular diversity in the Drosophila midbrain revealed by single-cell transcriptomics. eLife 7. eLife Sciences Publications Ltd.

Dambly-Chaudière, C., Jamet, E., Burri, M., Bopp, D., Basler, K., Hafen, E., Dumont, N., Spielmann, P., Ghysen, A. & Noll, M. (1992) The paired box gene pox neuro: A determiant of poly-innervated sense organs in Drosophila. Cell 69, 159–172. Elsevier.

Davis, F.P., Nern, A., Picard, S., Reiser, M.B., Rubin, G.M., Eddy, S.R. & Henry, G.L. (2020) A genetic, genomic, and computational resource for exploring neural circuit function. eLife 9. eLife Sciences Publications Ltd.

Deng, M., Wang, Y., Zhang, L., Yang, Y., Huang, S., Wang, J., Ge, H., Ishibashi, T. & Yan, Y. (2019) Single cell transcriptomic landscapes of pattern formation, proliferation and growth in Drosophila wing imaginal discs. Development (Cambridge) 146.

Desalvo, M.K., Hindle, S.J., Rusan, Z.M., Orng, S., Eddison, M., Halliwill, K. & Bainton, R.J. (2014) The Drosophila surface glia transcriptome: Evolutionary conserved blood-brain barrier processes. Frontiers in Neuroscience 8, 346. Frontiers Media S.A.

Dinges, G.F., Chockley, A.S., Bockemühl, T., Ito, K., Blanke, A. & Büschges, A. (2021) Location and arrangement of campaniform sensilla in Drosophila melanogaster. Journal of Comparative Neurology 529, 905–925. John Wiley & Sons, Ltd.

Ebbs, M.L. & Amrein, H. (2007) Taste and pheromone perception in the fruit fly Drosophila melanogaster. Pflugers Archiv European Journal of Physiology 454, 735–747. Springer Verlag.

Edwards, J.S., Swales, L.S. & Bate, M. (1993) The differentiation between neuroglia and connective tissue sheath in insect ganglia revisited: The neural lamella and perineurial sheath cells are absent in a mesodermless mutant of Drosophila. Journal of Comparative Neurology 333, 301–308.

Evans, C.J., Hartenstein, V. & Banerjee, U. (2003) Thicker than blood: Conserved mechanisms in Drosophila and vertebrate hematopoiesis. Developmental Cell 5, 673–690.

Everaerts, C., Farine, J.-P., Cobb, M. & Ferveur, J.-F. (2010) Drosophila Cuticular Hydrocarbons Revisited: Mating Status Alters Cuticular Profiles. PLoS ONE 5, e9607.

Everetts, N.J., Worley, M.I., Yasutomi, R., Yosef, N. & Hariharan, I.K. (2021) Single-cell transcriptomics of the Drosophila wing disc reveals instructive epithelium-to-myoblast interactions. eLife 10. eLife Sciences Publications Ltd.

Fichelson, P., Audibert, A., Simon, F. & Gho, M. (2005) Cell cycle and cell-fate determination in Drosophila neural cell lineages. Trends in Genetics 21, 413–420.

Fichelson, P. & Gho, M. (2003) The glial cell undergoes apoptosis in the microchaete lineage of Drosophila. Development 130, 123–133. Development.

Finzsch, M., Schreiner, S., Kichko, T., Reeh, P., Tamm, E.R., Bösl, M.R., Meijer, D. & Wegner, M. (2010) Sox10 is required for Schwann cell identity and progression beyond the immature Schwann cell stage. Journal of Cell Biology 189, 701–712. The Rockefeller University Press.

Fogarty, E.A., Kitzman, J.O. & Antonellis, A. (2020) SOX10-regulated promoter use defines isoform-specific gene expression in Schwann cells. BMC Genomics 21, 1–28. BioMed Central.

Franzdóttir, S.R., Engelen, D., Yuva-Aydemir, Y., Schmidt, I., Aho, A. & Klämbt, C. (2009) Switch in FGF signalling initiates glial differentiation in the Drosophila eye. Nature 460, 758–761.

Frenkel, L., Muraro, N.I., Beltrán González, A.N., Marcora, M.S., Bernabó, G., Hermann-Luibl, C., Romero, J.I., Helfrich-Förster, C., Castaño, E.M., Marino-Busjle, C., Calvo, D.J. & Ceriani, M.F. (2017) Organization of Circadian Behavior Relies on Glycinergic Transmission. Cell Reports 19, 72–85.

Fu, Y., Huang, X., Zhang, P., Van De Leemput, J. & Han, Z. (2020) Single-cell RNA sequencing identifies novel cell types in Drosophila blood. Journal of Genetics and Genomics 47, 175–186. Elsevier.

Gho, M., Lecourtois, M., Géraud, G., Posakony, J.W. & Schweisguth, F. (1996) Subcellular localization of suppressor of hairless in Drosophila sense organ cells during Notch signalling. Development 122, 1673–1682. The Company of Biologists.

Gilsohn, E. & Volk, T. (2010) Slowdown promotes muscle integrity by modulating integrin-mediated adhesion at the myotendinous junction. Development 137, 785–794. The Company of Biologists.

Godt, D., Couderc, J.L., Cramton, S.E. & Laski, F.A. (1993) Pattern formation in the limbs of Drosophila: Bric a brac is expressed in both a gradient and a wave-like pattern and is required for specification and proper segmentation of the tarsus. Development 119, 799–812. Development.

Grigorian, M., Mandal, L. & Hartenstein, V. (2011) Hematopoiesis at the onset of metamorphosis: Terminal differentiation and dissociation of the Drosophila lymph gland. Development Genes and Evolution 221, 121–131. Dev Genes Evol.

Grillo-Hill, B.K. & Wolff, T. (2009) Dynamic cell shapes and contacts in the developing Drosophila retina are regulated by the Ig cell adhesion protein hibris. Developmental Dynamics 238, 2223–2234. John Wiley & Sons, Ltd.

Guo, L., Zhang, N. & Simpson, J.H. (2022) Descending neurons coordinate anterior grooming behavior in Drosophila. Current Biology 32, 823–833.e4. Cell Press.

Hafemeister, C. & Satija, R. (2019) Normalization and variance stabilization of single-cell RNA-seq data using regularized negative binomial regression. Genome Biology 20, 1–15. BioMed Central Ltd.

Halter, D.A., Urban, J., Rickert, C., Ner, S.S., Ito, K., Travers, A.A. & Technau, G.M. (1995) The homeobox gene repo is required for the differentiation and maintenance of glia function in the embryonic nervous system of Drosophila melanogaster. Development 121, 317–332. The Company of Biologists.

Hannah-Alava, A. (1958) Morphology and chaetotaxy of the legs of Drosophila melanogaster. Journal of Morphology 103, 281–310. John Wiley & Sons, Ltd.

Hao, I., Green, R.B., Dunaevsky, O., Lengyel, J.A. & Rauskolb, C. (2003) The odd-skipped family of zinc finger genes promotes Drosophila leg segmentation. Developmental Biology 263, 282–295. Academic Press Inc.

Hao, Y., Hao, S., Andersen-Nissen, E., Mauck, W.M., Zheng, S., Butler, A., Lee, M.J., Wilk, A.J., Darby, C., Zager, M., Hoffman, P., Stoeckius, M., Papalexi, E., Mimitou, E.P., Jain, J., ET AL. (2021) Integrated analysis of multimodal single-cell data. Cell 184, 3573–3587.e29. Cell.

He, P., Williams, B.A., Trout, D., Marinov, G.K., Amrhein, H., Berghella, L., Goh, S.T., Plajzer-Frick, I., Afzal, V., Pennacchio, L.A., Dickel, D.E., Visel, A., Ren, B., Hardison, R.C., Zhang, Y., ET AL. (2020) The changing mouse embryo transcriptome at whole tissue and single-cell resolution. Nature 583, 760–767. Nature Publishing Group.

He, Z., Luo, Y., Shang, X., Sun, J.S. & Carlson, J.R. (2019) Chemosensory sensilla of the Drosophila wing express a candidate ionotropic pheromone receptor. PLOS Biology 17, e2006619. Public Library of Science.

Held, L.I. (1979) A high-resolution morphogenetic map of the second-leg basitarsus in Drosophila melanogaster. Wilhelm Roux’s Archives of Developmental Biology 187, 129–150.

Held, L.I. (1990) Sensitive periods for abnormal patterning on a leg segment in Drosophila melanogaster. Rouxs Archives of Developmental Biology 199, 31–47. Springer.

Held, L.I. (2002) Bristles induce bracts via the EGFR pathway on Drosophila legs. Mechanisms of Development 117, 225–234. Elsevier.

Hopkins, B.R. & Kopp, A. (2021) Evolution of sexual development and sexual dimorphism in insects. Current Opinion in Genetics & Development 69, 129–139. Elsevier Ltd.

Hultmark, D. & Andó, I. (2022) Hematopoietic plasticity mapped in Drosophila and other insects. eLife 11.

Hung, R.-J.J., Hu, Y., Kirchner, R., Liu, Y., Xu, C., Comjean, A., Tattikota, S.G., Li, F., Song, W.R., Sui, S.H., Perrimon, N., Xu, C., Comjean, A., Tattikota, S.G., Song, W.R., ET AL. (2020) A cell atlas of the adult Drosophila midgut. Proceedings of the National Academy of Sciences of the United States of America 117, 1514–1523.

Inbal, A., Levanon, D. & Salzberg, A. (2003) Multiple roles for u-turn/ventral veinless in the development of Drosophila PNS. Development 130, 2467–2478. Development.

Ittner, E., Hartwig, A.C., Elsesser, O., Wüst, H.M., Fröb, F., Wedel, M., Schimmel, M., Tamm, E.R., Wegner, M. & Sock, E. (2021) SoxD transcription factor deficiency in Schwann cells delays myelination in the developing peripheral nervous system. Scientific Reports 11, 1–11. Nature Publishing Group.

Janssens, J., Aibar, S., Taskiran, I.I., Ismail, J.N., Gomez, A.E., Aughey, G., Spanier, K.I., De Rop, F. V., González-Blas, C.B., Dionne, M., Grimes, K., Quan, X.J., Papasokrati, D., Hulselmans, G., Makhzami, S., ET AL. (2022) Decoding gene regulation in the fly brain. Nature 601, 630–636. Nature Publishing Group.

Kallman, B.R., Kim, H. & Scott, K. (2015) Excitation and inhibition onto central courtship neurons biases Drosophila mate choice. eLife 4. eLife Sciences Publications Ltd.

Kania, A. & Bellen, H.J. (1995) Mutations in neuromusculin, a gene encoding a cell adhesion molecule, cause nervous system defects. Roux’s Archives of Developmental Biology 204, 259–270. Springer-Verlag.

Kania, A., Han, P.L., Kim, Y.T. & Bellen, H. (1993) Neuromusculin, a Drosophila Gene Expressed in Peripheral Neuronal Precursors and Muscles, Encodes a Cell Adhesion Molecule. Neuron 11, 673–687. Elsevier.

Kavaler, J., Fu, W., Duan, H., Noll, M. & Posakony, J.W. (1999) An essential role for the Drosophila Pax2 homolog in the differentiation of adult sensory organs. Development 126, 2261–2272.

Keil, T.A. (1997a) Functional Morphology of Insect Mechanoreceptors. Microsc. Res. Tech 39, 506–531. Wiley-Liss, Inc.

Keil, T.A. (1997b) Comparative morphogenesis of sensilla: A review. International Journal of Insect Morphology and Embryology 26, 151–160.

Kerber, B., Monge, I., Mueller, M., Mitchell, P.J. & Cohen, S.M. (2001) The AP-2 transcription factor is required for joint formation and cell survival in Drosophila leg development. Development 128, 1231–1238. The Company of Biologists.

Khallaf, M.A., Cui, R., Weißflog, J., Erdogmus, M., Svatoš, A., Dweck, H.K.M., Valenzano, D.R., Hansson, B.S. & Knaden, M. (2021) Large-scale characterization of sex pheromone communication systems in Drosophila. Nature Communications 12, 1–14. Nature Publishing Group.

Knecht, Z.A., Silbering, A.F., Ni, L., Klein, M., Budelli, G., Bell, R., Abuin, L., Ferrer, A.J., Samuel, A.D.T., Benton, R. & Garrity, P.A. (2016) Distinct combinations of variant ionotropic glutamate receptors mediate thermosensation and hygrosensation in drosophila. eLife 5. eLife Sciences Publications Ltd.

Kocks, C., Cho, J.H., Nehme, N., Ulvila, J., Pearson, A.M., Meister, M., Strom, C., Conto, S.L., Hetru, C., Stuart, L.M., Stehle, T., Hoffmann, J.A., Reichhart, J.M., Ferrandon, D., Rämet, M., ET AL. (2005) Eater, a transmembrane protein mediating phagocytosis of bacterial pathogens in Drosophila. Cell 123, 335–346. Elsevier B.V.

Koh, T.W., He, Z., Gorur-Shandilya, S., Menuz, K., Larter, N.K., Stewart, S. & Carlson, J.R. (2014) The Drosophila IR20a Clade of Ionotropic Receptors Are Candidate Taste and Pheromone Receptors. Neuron 83, 850–865.

Kopp, A. (2011) Drosophila sex combs as a model of evolutionary innovations. Evolution and Development 13, 504–522.

Kudron, M.M., Victorsen, A., Gevirtzman, L., Hillier, L.W., Fisher, W.W., Vafeados, D., Kirkey, M., Hammonds, A.S., Gersch, J., Ammouri, H., Wall, M.L., Moran, J., Steffen, D., Szynkarek, M., Seabrook-Sturgis, S., ET AL. (2018) The ModERN Resource: Genome-Wide Binding Profiles for Hundreds of Drosophila and Caenorhabditis elegans Transcription Factors. Genetics 208, 937–949. Oxford Academic.

Kurucz, É., Márkus, R., Zsámboki, J., Folkl-Medzihradszky, K., Darula, Z., Vilmos, P., Udvardy, A., Krausz, I., Lukacsovich, T., Gateff, E., Zettervall, C.J., Hultmark, D. & Andó, I. (2007) Nimrod, a Putative Phagocytosis Receptor with EGF Repeats in Drosophila Plasmatocytes. Current Biology 17, 649–654.

Lai, E.C. & Orgogozo, V. (2004) A hidden program in Drosophila peripheral neurogenesis revealed: Fundamental principles underlying sensory organ diversity. Developmental Biology 269, 1–17. Academic Press Inc.

Lassetter, A.P., Corty, M.M., Barria, R., Sheehan, A.E., Hill, J., Aicher, S.A., Fox, A.N. & Freeman, M.R. (2021) Glial TGFβ activity promotes axon survival in peripheral nerves. bioRxiv, 2021.09.02.458753. Cold Spring Harbor Laboratory.

Lawrence, P.A., Struhl, G. & Morata, G. (1979) Bristle patterns and compartment boundaries in the tarsi of Drosophila. Journal of Embryology and Experimental Morphology VOL.51, 195–208.

Li, H., Horns, F., Wu, B., Xie, Q., Li, J., Li, T., Luginbuhl, D.J., Quake, S.R. & Luo, L. (2017a) Classifying Drosophila Olfactory Projection Neuron Subtypes by Single-Cell RNA Sequencing. Cell 171, 1206–1220.e22. Cell Press.

Li, H., Janssens, J., De Waegeneer, M., Kolluru, S.S., Davie, K., Gardeux, V., Saelens, W., David, F.P.A., Brbić, M., Spanier, K., Leskovec, J., Mclaughlin, C.N., Xie, Q., Jones, R.C., Brueckner, K., ET AL. (2022) Fly Cell Atlas: A single-nucleus transcriptomic atlas of the adult fruit fly. Science 375.

Li, H., Watson, A., Olechwier, A., Anaya, M., Sorooshyari, S.K., Harnett, D.P., Lee, H.-K. (PETER), Vielmetter, J., Fares, M.A., Garcia, K.C., özkan, E., Labrador, J.-P. & Zinn, K. (2017b) Deconstruction of the beaten Path-Sidestep interaction network provides insights into neuromuscular system development. eLife 6. eLife Sciences Publications Ltd.

Lilly, B., Zhao, B., Ranganayakulu, G., Paterson, B.M., Schulz, R.A. & Olson, E.N. (1995) Requirement of MADS domain transcription factor D-MEF2 for muscle formation in Drosophila. Science 267, 688–693. Science.

Ling, F., Dahanukar, A., Weiss, L.A., Kwon, J.Y. & Carlson, J.R. (2014) The Molecular and Cellular Basis of Taste Coding in the Legs of Drosophila. Journal of Neuroscience 34, 7148–7164. Society for Neuroscience.

Loker, R. & Mann, R.S. (2022) Divergent expression of paralogous genes by modification of shared enhancer activity through a promoter-proximal silencer. Current Biology 32, 3545–3555.e4. Cell Press.

Lu, B., Lamora, A., Sun, Y., Welsh, M.J. & Ben-Shahar, Y. (2012) Ppk23-dependent chemosensory functions contribute to courtship behavior in drosophila melanogaster. PLoS Genetics 8, e1002587. Public Library of Science.

Mackay, T.F.C., Richards, S., Stone, E.A., Barbadilla, A., Ayroles, J.F., Zhu, D., Casillas, S., Han, Y., Magwire, M.M., Cridland, J.M., Richardson, M.F., Anholt, R.R.H., Barrón, M., Bess, C., Blankenburg, K.P., ET AL. (2012) The Drosophila melanogaster Genetic Reference Panel. Nature 482, 173–178. Nature Publishing Group.

Magklara, A. & Lomvardas, S. (2013) Stochastic gene expression in mammals: Lessons from olfaction. Trends in Cell Biology 23, 449–456. NIH Public Access.

Mahajan-Miklos, S. & Cooley, L. (1994) The villin-like protein encoded by the Drosophila quail gene is required for actin bundle assembly during oogenesis. Cell 78, 291–301.

Mamiya, A., Gurung, P., Correspondence, J.C.T. & Tuthill, J.C. (2018) Neural Coding of Leg Proprioception in Drosophila Article Neural Coding of Leg Proprioception in Drosophila. Neuron 100, 636–650.e6.

Massey, J.H., Chung, D., Siwanowicz, I., Stern, D.L. & Wittkopp, P.J. (2019) The yellow gene influences Drosophila male mating success through sex comb melanization. eLife 8, 673756. Cold Spring Harbor Laboratory.

Mayer, F., Mayer, N., Chinn, L., Pinsonneault, R.L., Kroetz, D. & Bainton, R.J. (2009) Evolutionary conservation of vertebrate blood-brain barrier chemoprotective mechanisms in drosophila. Journal of Neuroscience 29, 3538–3550. J Neurosci.

Mcginnis, C.S., Murrow, L.M. & Gartner, Z.J. (2019) DoubletFinder: Doublet Detection in Single-Cell RNA Sequencing Data Using Artificial Nearest Neighbors. Cell Systems 8, 329–337.e4. Cell Press.

Mckelvey, E.G.Z., Gyles, J.P., Michie, K., Barquín Pancorbo, V., Sober, L., Kruszewski, L.E., Chan, A. & Fabre, C.C.G. (2021) Drosophila females receive male substrate-borne signals through specific leg neurons during courtship. Current Biology 31, 3894–3904.e5. Cell Press.

Mclaughlin, C.N., Brbić, M., Xie, Q., Li, T., Horns, F., Kolluru, S.S., Kebschull, J.M., Vacek, D., Xie, A., Li, J., Jones, R.C., Leskovec, J., Quake, S.R., Luo, L. & Li, H. (2021) Single-cell transcriptomes of developing and adult olfactory receptor neurons in drosophila. eLife 10, 1–37. eLife Sciences Publications Ltd.

Mellert, D.J., Knapp, J.M., Manoli, D.S., Meissner, G.W. & Baker, B.S. (2010) Midline crossing by gustatory receptor neuron axons is regulated by fruitless, doublesex and the Roundabout receptors. Development 137, 323–332. Development.

Meyer, S., Schmidt, I. & Klämbt, C. (2014) Glia ECM interactions are required to shape the Drosophila nervous system. Mechanisms of Development 133, 105–116. Elsevier.

Miller, S.W., Avidor-Reiss, T., Polyanovsky, A. & Posakony, J.W. (2009) Complex interplay of three transcription factors in controlling the tormogen differentiation program of Drosophila mechanoreceptors. Developmental Biology 329, 386–399. Academic Press.

Mirth, C. & Akam, M. (2002) Joint development in the Drosophila leg: Cell movements and cell populations. Developmental Biology 246, 391–406.

Mittnenzweig, M., Mayshar, Y., Cheng, S., Ben-Yair, R., Hadas, R., Rais, Y., Chomsky, E., Reines, N., Uzonyi, A., Lumerman, L., Lifshitz, A., Mukamel, Z., Orenbuch, A.H., Tanay, A. & Stelzer, Y. (2021) A single-embryo, single-cell time-resolved model for mouse gastrulation. Cell 184, 2825–2842.e22. Cell Press.

Monge, I., Krishnamurthy, R., Sims, D., Hirth, F., Spengler, M., Kammermeier, L., Reichert, H. & Mitchell, P.J. (2001) Drosophila transcription factor AP-2 in proboscis, leg and brain central complex development. Development 128, 1239–1252. The Company of Biologists.

Montell, C. (2009) A taste of the Drosophila gustatory receptors. Current Opinion in Neurobiology 19, 345–353. NIH Public Access.

Mou, X., Duncan, D.M., Baehrecke, E.H. & Duncan, I. (2012) Control of target gene specificity during metamorphosis by the steroid response gene E93. Proceedings of the National Academy of Sciences of the United States of America 109, 2949–2954. National Academy of Sciences.

Nangia, V., O’Connell, J., Chopra, K., Qing, Y., Reppert, C., Chai, C.M., Bhasiin, K. & Colodner, K.J. (2021) Genetic reduction of tyramine β hydroxylase suppresses Tau toxicity in a Drosophila model of tauopathy. Neuroscience Letters 755, 135937. Elsevier.

Natori, K., Tajiri, R., Furukawa, S. & Kojima, T. (2012) Progressive tarsal patterning in the Drosophila by temporally dynamic regulation of transcription factor genes. Developmental Biology 361, 450–462. Academic Press.

Nayak, S. V. & Singh, R.N. (1983) Sensilla on the tarsal segments and mouthparts of adult Drosophila melanogaster meigen (Diptera : Drosophilidae). International Journal of Insect Morphology and Embryology 12, 273–291.

Ng, C.S. & Kopp, A. (2008) Sex combs are important for male mating success in Drosophila melanogaster. Behavior Genetics 38, 195–201.

Noordermeer, J.N., Kopczynski, C.C., Fetter, R.D., Bland, K.S., Chen, W.Y. & Goodman, C.S. (1998) Wrapper, a novel member of the Ig superfamily, is expressed by midline glia and is required for them to ensheath commissural axons in Drosophila. Neuron 21, 991–1001. Neuron.

Nottebohm, E., Dambly-Chaudière, C. & Ghysen, A. (1992) Connectivity of chemosensory neurons is controlled by the gene poxn in Drosophila. Nature 359, 829–832. Nature Publishing Group.

Nottebohm, E., Usui, A., Therianos, S., Kimura, K. ICHI, Dambly-Chaudiére, C. & Ghysen, A. (1994) The gene poxn controls different steps of the formation of chemosensory organs in drosophila. Neuron 12, 25–34. Elsevier.

O’Connor-Giles, K.M. & Skeath, J.B. (2003) Numb inhibits membrane localization of sanpodo, a four-pass transmembrane protein, to promote asymmetric divisions in Drosophila. Developmental Cell 5, 231–243.

Olofsson, B. & Page, D.T. (2005) Condensation of the central nervous system in embryonic Drosophila is inhibited by blocking hemocyte migration or neural activity. Developmental Biology 279, 233–243. Academic Press.

Özel, M.N., Simon, F., Jafari, S., Holguera, I., Chen, Y.C., Benhra, N., El-Danaf, R.N., Kapuralin, K., Malin, J.A., Konstantinides, N. & Desplan, C. (2021) Neuronal diversity and convergence in a visual system developmental atlas. Nature 589, 88–95. Nature Publishing Group.

Park, Y., Rangel, C., Reynolds, M.M., Caldwell, M.C., Johns, M., Nayak, M., Welsh, C.J.R., Mcdermott, S. & Datta, S. (2003) Drosophila Perlecan modulates FGF and Hedgehog signals to activate neural stem cell division. Developmental Biology 253, 247–257. Academic Press.

Peng, Y., Han, C. & Axelrod, J.D. (2012) Planar Polarized Protrusions Break the Symmetry of EGFR Signaling during Drosophila Bract Cell Fate Induction. Developmental Cell 23, 507–518.

Phillips, A.M., Smart, R., Strauss, R., Brembs, B. & Kelly, L.E. (2005) The Drosophila black enigma: The molecular and behavioural characterization of the black1 mutant allele. Gene 351, 131–142. Elsevier.

Plass, M., Solana, J., Alexander Wolf, F., Ayoub, S., Misios, A., Glažar, P., Obermayer, B., Theis, F.J., Kocks, C. & Rajewsky, N. (2018) Cell type atlas and lineage tree of a whole complex animal by single-cell transcriptomics. Science 360. American Association for the Advancement of Science.

Prelic, S., Pal Mahadevan, V., Venkateswaran, V., Lavista-Llanos, S., Hansson, B.S. & Wicher, D. (2022) Functional Interaction Between Drosophila Olfactory Sensory Neurons and Their Support Cells. Frontiers in Cellular Neuroscience 15, 555. Frontiers Media S.A.

Pringle, J.W.S. (1938) Proprioception In Insects. Journal of Experimental Biology 15, 114–131. The Company of Biologists.

Qiu, C., Cao, J., Martin, B.K., Li, T., Welsh, I.C., Srivatsan, S., Huang, X., Calderon, D., Noble, W.S., Disteche, C.M., Murray, S.A., Spielmann, M., Moens, C.B., Trapnell, C. & Shendure, J. (2022) Systematic reconstruction of cellular trajectories across mouse embryogenesis. Nature Genetics 54, 328–341. Nature Publishing Group.

Rahman, S.R., Terranova, T., Tian, L. & Hines, H.M. (2021) Developmental Transcriptomics Reveals a Gene Network Driving Mimetic Color Variation in a Bumble Bee. Genome Biology and Evolution 13. Oxford University Press.

Rauskolb, C. & Irvine, K.D. (1999) Notch-Mediated Segmentation and Growth Control of the Drosophila Leg. Developmental Biology 210, 339–350. Academic Press.

Reed, C.T., Murphy, C. & Fristrom, D. (1975) The ultrastructure of the differentiating pupal leg of Drosophila melanogaster. Wilhelm Roux’s Archives of Developmental Biology 178, 285–302. Springer-Verlag.

Ren, N., He, B., Stone, D., Kirakodu, S. & Adler, P.N. (2006) The shavenoid gene of Drosophila encodes a novel actin cytoskeleton interacting protein that promotes wing hair morphogenesis. Genetics 172, 1643–1653.

Rizki, M.T. & Rizki, R.M. (1959) Functional significance of the crystal cells in the larva of Drosophila melanogaster. The Journal of biophysical and biochemical cytology 5, 235–240.

Robinett, C.C., Vaughan, A.G., Knapp, J.-M. & Baker, B.S. (2010) Sex and the Single Cell. II. There Is a Time and Place for Sex. PLoS Biology 8, e1000365.

Rodriguez, I., Hernandez, R., Modolell, J. & Ruiz-Gomez, M. (1990) Competence to develop sensory organs is temporally and spatially regulated in Drosophila epidermal primordia. EMBO Journal 9, 3583–3592.

Ross, J., Kuzin, A., Brody, T. & Odenwald, W.F. (2015) cis-regulatory analysis of the Drosophila pdm locus reveals a diversity of neural enhancers. BMC Genomics 16. BioMed Central.

Ruiz-Losada, M., Pérez-Reyes, C. & Estella, C. (2021) Role of the Forkhead Transcription Factors Fd4 and Fd5 During Drosophila Leg Development. Frontiers in Cell and Developmental Biology 9, 2072. Frontiers Media S.A.

Rushton, E., Rohrbough, J. & Broadie, K. (2009) Presynaptic secretion of mind-the-gap organizes the synaptic extracellular matrix-integrin interface and postsynaptic environments. Developmental Dynamics 238, 554–571. John Wiley & Sons, Ltd.

Rust, K., Byrnes, L.E., Yu, K.S., Park, J.S., Sneddon, J.B., Tward, A.D. & Nystul, T.G. (2020) A single-cell atlas and lineage analysis of the adult Drosophila ovary. Nature Communications 11, 1–17. Nature Publishing Group.

Sánchez-Alcañiz, J.A., Silbering, A.F., Croset, V., Zappia, G., Sivasubramaniam, A.K., Abuin, L., Sahai, S.Y., Münch, D., Steck, K., Auer, T.O., Cruchet, S., Neagu-Maier, G.L., Sprecher, S.G., Ribeiro, C., Yapici, N., ET AL. (2018) An expression atlas of variant ionotropic glutamate receptors identifies a molecular basis of carbonation sensing. Nature Communications 9, 1–14. Nature Publishing Group.

Sanfilippo, P., Smibert, P., Duan, H. & Lai, E.C. (2016) Neural specificity of the RNA-binding protein elav is achieved by post-transcriptional repression in non-neural tissues. Development (Cambridge) 143, 4474–4485. Company of Biologists.

Sasse, S. & Klämbt, C. (2016) Repulsive Epithelial Cues Direct Glial Migration along the Nerve. Developmental Cell 39, 696–707. Cell Press.

Satija, R., Farrell, J.A., Gennert, D., Schier, A.F. & Regev, A. (2015) Spatial reconstruction of single-cell gene expression data. Nature Biotechnology 33, 495–502. Nature Publishing Group.

Saur, A.L., Fröb, F., Weider, M. & Wegner, M. (2021) Formation of the node of Ranvier by Schwann cells is under control of transcription factor Sox10. Glia 69, 1464–1477. John Wiley & Sons, Ltd.

Schmidt, H.R. & Benton, R. (2020) Molecular mechanisms of olfactory detection in insects: Beyond receptors: Insect olfactory detection mechanisms. Open Biology 10. Royal Society Publishing.

Schwabe, T., Bainton, R.J., Fetter, R.D., Heberlein, U. & Gaul, U. (2005) GPCR signaling is required for blood-brain barrier formation in Drosophila. Cell 123, 133–144. Elsevier B.V.

Seroka, A., Lai, S.L. & Doe, C.Q. (2022) Transcriptional profiling from whole embryos to single neuroblast lineages in Drosophila. Developmental Biology 489, 21–33. Elsevier Inc.

Shapira, S., Bakhrat, A., Bitan, A. & Abdu, U. (2011) The Drosophila javelin Gene Encodes a Novel Actin-Associated Protein Required for Actin Assembly in the Bristle. Molecular and Cellular Biology 31, 4582–4592. American Society for Microbiology.

Simon, F., Fichelson, P., Gho, M. & Audibert, A. (2009) Notch and Prospero repress proliferation following cyclin E overexpression in the Drosophila bristle lineage. PLoS Genetics 5, e1000594. Public Library of Science.

Simon, F., Ramat, A., Louvet-Vallée, S., Lacoste, J., Burg, A., Audibert, A. & Gho, M. (2019) Shaping of Drosophila neural cell lineages through coordination of cell proliferation and cell fate by the BTB-ZF transcription factor tramtrack-69. Genetics 212, 773–788. Genetics.

Soler, C., Daczewska, M., Da Ponte, J.P., Dastugue, B. & Jagla, K. (2004) Coordinated development of muscles and tendons of the Drosophila leg. Development 131, 6041–6051. The Company of Biologists.

Spieth, H.T. (1974) Courtship behavior in Drosophila. Annual review of entomology 19, 385–405. Annual Reviews 4139 El Camino Way, P.O. Box 10139, Palo Alto, CA 94303-0139, USA.

Starostina, E., Liu, T., Vijayan, V., Zheng, Z., Siwicki, K.K. & Pikielny, C.W. (2012) A drosophila DEG/ENaC subunit functions specifically in gustatory neurons required for male courtship behavior. Journal of Neuroscience 32, 4665–4674. Society for Neuroscience.

Stocker, R.F. (1994) The organization of the chemosensory system in Drosophila melanogaster: a rewiew. Cell and Tissue Research 275, 3–26. Springer.

Stork, T., Bernardos, R. & Freeman, M.R. (2012) Analysis of glial cell development and function in Drosophila. Cold Spring Harbor Protocols 7, 1–17. Cold Spring Harbor Laboratory Press.

Stork, T., Engelen, D., Krudewig, A., Silies, M., Bainton, R.J. & Klämbt, C. (2008) Organization and function of the blood-brain barrier in Drosophila. Journal of Neuroscience 28, 587–597. Society for Neuroscience.

Stork, T., Sheehan, A., Tasdemir-Yilmaz, O.E. & Freeman, M.R. (2014) Neuron-Glia interactions through the heartless fgf receptor signaling pathway mediate morphogenesis of drosophila astrocytes. Neuron 83, 388–403. Cell Press.

Stork, T., Thomas, S., Rodrigues, F., Silies, M., Naffin, E., Wenderdel, S. & Klämbt, C. (2009) Drosophila Neurexin IV stabilizes neuron-glia interactions at the CNS midline by binding to Wrapper. Development (Cambridge, England) 136, 1251–1261. Development.

Stuart, T., Butler, A., Hoffman, P., Hafemeister, C., Papalexi, E., Mauck, W.M., Hao, Y., Stoeckius, M., Smibert, P. & Satija, R. (2019) Comprehensive Integration of Single-Cell Data. Cell 177, 1888–1902.e21. Cell.

Subramanian, A., Wayburn, B., Bunch, T. & Volk, T. (2007) Thrombospondin-mediated adhesion is essential for the formation of the myotendinous junction in Drosophila. Development 134, 1269–1278. Development.

Suzuki, T., Hasegawa, E., Nakai, Y., Kaido, M., Takayama, R. & Sato, M. (2016) Formation of Neuronal Circuits by Interactions between Neuronal Populations Derived from Different Origins in the Drosophila Visual Center. Cell Reports 15, 499–509. Cell Press.

Tan, L., Zhang, K.X., Pecot, M.Y., Nagarkar-Jaiswal, S., Lee, P.T., Takemura, S.Y., Mcewen, J.M., Nern, A., Xu, S., Tadros, W., Chen, Z., Zinn, K., Bellen, H.J., Morey, M. & Zipursky, S.L. (2015) Ig Superfamily Ligand and Receptor Pairs Expressed in Synaptic Partners in Drosophila. Cell 163, 1756–1769. NIH Public Access.

Tanaka, K., Barmina, O. & Kopp, A. (2009) Distinct developmental mechanisms underlie the evolutionary diversification of Drosophila sex combs. Proceedings of the National Academy of Sciences 106, 4764–4769.

Tanaka, K., Barmina, O., Sanders, L.E., Arbeitman, M.N. & Kopp, A. (2011) Evolution of sex-specific traits through changes in HOX-dependent doublesex expression. PLoS Biology 9, e1001131.

Tattikota, S.G., Cho, B., Liu, Y., Hu, Y., Barrera, V., Steinbaugh, M.J., Yoon, S.-H., Comjean, A., Li, F., Dervis, F., Hung, R.-J., Nam, J.-W., Ho Sui, S., Shim, J. & Perrimon, N. (2020) A single-cell survey of Drosophila blood. eLife 9, 1–35. eLife Sciences Publications Ltd.

Thistle, R., Cameron, P., Ghorayshi, A., Dennison, L. & Scott, K. (2012) Contact Chemoreceptors Mediate Male-Male Repulsion and Male-Female Attraction during Drosophila Courtship. Cell 149, 1140–1151. Elsevier.

Tobler, H. (1969) [Influencing of bristle differentiation and pattern formation by mitomycin in Drosophila melanogaster]. Experientia 25, 213–214.

Toda, H., Zhao, X. & Dickson, B.J. (2012) The Drosophila Female Aphrodisiac Pheromone Activates ppk23+ Sensory Neurons to Elicit Male Courtship Behavior. Cell Reports 1, 599–607. Cell Press.

Tokunaga, C. (1962) Cell lineage and differentiation on the male foreleg of Drosophila melanogaster. Developmental Biology 4, 489–516. Dev Biol.

Turnescu, T., Arter, J., Reiprich, S., Tamm, E.R., Waisman, A. & Wegner, M. (2018) Sox8 and Sox10 jointly maintain myelin gene expression in oligodendrocytes. Glia 66, 279–294. Glia.

Tuthill, J.C. & Wilson, R.I. (2016) Mechanosensation and Adaptive Motor Control in Insects. Current Biology 26, R1022–R1038. Elsevier.

Upadhyay, A., Kandachar, V., Zitserman, D., Tong, X. & Roegiers, F. (2013) Sanpodo controls sensory organ precursor fate by directing notch trafficking and binding γ-secretase. Journal of Cell Biology 201, 439–448.

Usui, T., Shima, Y., Shimada, Y., Hirano, S., Burgess, R.W., Schwarz, T.L., Takeichi, M. & Uemura, T. (1999) Flamingo, a seven-pass transmembrane cadherin, regulates planar cell polarity under the control of Frizzled. Cell 98, 585–595. Cell Press.

Vijayan, V., Thistle, R., Liu, T., Starostina, E. & Pikielny, C.W. (2014) Drosophila Pheromone-Sensing Neurons Expressing the ppk25 Ion Channel Subunit Stimulate Male Courtship and Female Receptivity. PLoS Genetics 10, e1004238. Public Library of Science.

Vosshall, L.B. & Stocker, R.F. (2007) Molecular architecture of smell and taste in Drosophila. Annual Review of Neuroscience 30, 505–533. Annual Reviews.

Walker, R.G., Willingham, A.T. & Zuker, C.S. (2000) A Drosophila mechanosensory transduction channel. Science 287, 2229–2234. American Association for the Advancement of Science.

Wayburn, B. & Volk, T. (2009) LRT, a tendon-specific leucine-rich repeat protein, promotes muscle-tendon targeting through its interaction with Robo. Development 136, 3607–3615. The Company of Biologists.

Wei, S.Y., Escudero, L.M., Yu, F., Chang, L.H., Chen, L.Y., Ho, Y.H., Lin, C.M., Chou, C.S., Chia, W., Modolell, J. & Hsu, J.C. (2005) Echinoid is a component of adherens junctions that cooperates with DE-cadherin to mediate cell adhesion. Developmental Cell 8, 493–504. Cell Press.

Wheeler, S.R., Banerjee, S., Blauth, K., Rogers, S.L., Bhat, M.A. & Crews, S.T. (2009) Neurexin IV and Wrapper interactions mediate Drosophila midline glial migration and axonal ensheathment. Development 136, 1147–1157.

Wu, B., Li, J., Chou, Y.H., Luginbuhl, D. & Luo, L. (2017) Fibroblast growth factor signaling instructs ensheathing glia wrapping of Drosophila olfactory glomeruli. Proceedings of the National Academy of Sciences of the United States of America 114, 7505–7512.

Xie, Q., Brbic, M., Horns, F., Kolluru, S.S., Jones, R.C., Li, J., Reddy, A.R., Xie, A., Kohani, S., Li, Z., Mclaughlin, C.N., Li, T., Xu, C., Vacek, D., Luginbuhl, D.J., ET AL. (2021) Temporal evolution of single-cell transcriptomes of drosophila olfactory projection neurons. eLife 10, 1–55. eLife Sciences Publications Ltd.

Xie, X. & Auld, V.J. (2011) Integrins are necessary for the development and maintenance of the glial layers in the Drosophila peripheral nerve. Development 138, 3813–3822. Development.

Xie, X., Shi, Q., Wu, P., Zhang, X., Kambara, H., Su, J., Yu, H., Park, S.Y., Guo, R., Ren, Q., Zhang, S., Xu, Y., Silberstein, L.E., Cheng, T., Ma, F., ET AL. (2020) Single-cell transcriptome profiling reveals neutrophil heterogeneity in homeostasis and infection. Nature Immunology 21, 1119–1133. Nature Publishing Group.

Xiong, W.C., Okano, H., Patel, N.H., Blendy, J.A. & Montell, C. (1994) repo encodes a glial-specific homeo domain protein required in the Drosophila nervous system. Genes and Development 8, 981–994.

Yildirim, K., Petri, J., Kottmeier, R. & Klämbt, C. (2019) Drosophila glia: Few cell types and many conserved functions. Glia 67, 5–26. John Wiley and Sons Inc.

Yuasa, Y., Okabe, M., Yoshikawa, S., Tabuchi, K., Xiong, W.C., Hiromi, Y. & Okano, H. (2003) Drosophila homeodomain protein REPO controls glial differentiation by cooperating with ETS and BTB transcription factors. Development 130, 2419–2428. The Company of Biologists.

Zhang, M.J., Pisco, A.O., Darmanis, S. & Zou, J. (2021) Mouse aging cell atlas analysis reveals global and cell type-specific aging signatures. eLife 10. eLife Sciences Publications Ltd.

Zhang, Y. V., Ni, J. & Montell, C. (2013) The molecular basis for attractive salt-taste coding in drosophila. Science 340, 1334–1338. American Association for the Advancement of Science.

Zhou, P., Yang, X. LOU, Wang, X.G., Hu, B., Zhang, L., Zhang, W., Si, H.R., Zhu, Y., Li, B., Huang, C.L., Chen, H.D., Chen, J., Luo, Y., Guo, H., Jiang, R. DI, ET AL. (2020) A pneumonia outbreak associated with a new coronavirus of probable bat origin. Nature 579, 270–273. Nature Publishing Group.

Zill, S., Schmitz, J. & Büschges, A. (2004) Load sensing and control of posture and locomotion. Arthropod Structure and Development 33, 273–286. Elsevier.

