## Supplementary material for "Charting the development of *Drosophila* leg sensory organs at single-cell resolution": Figure supplements

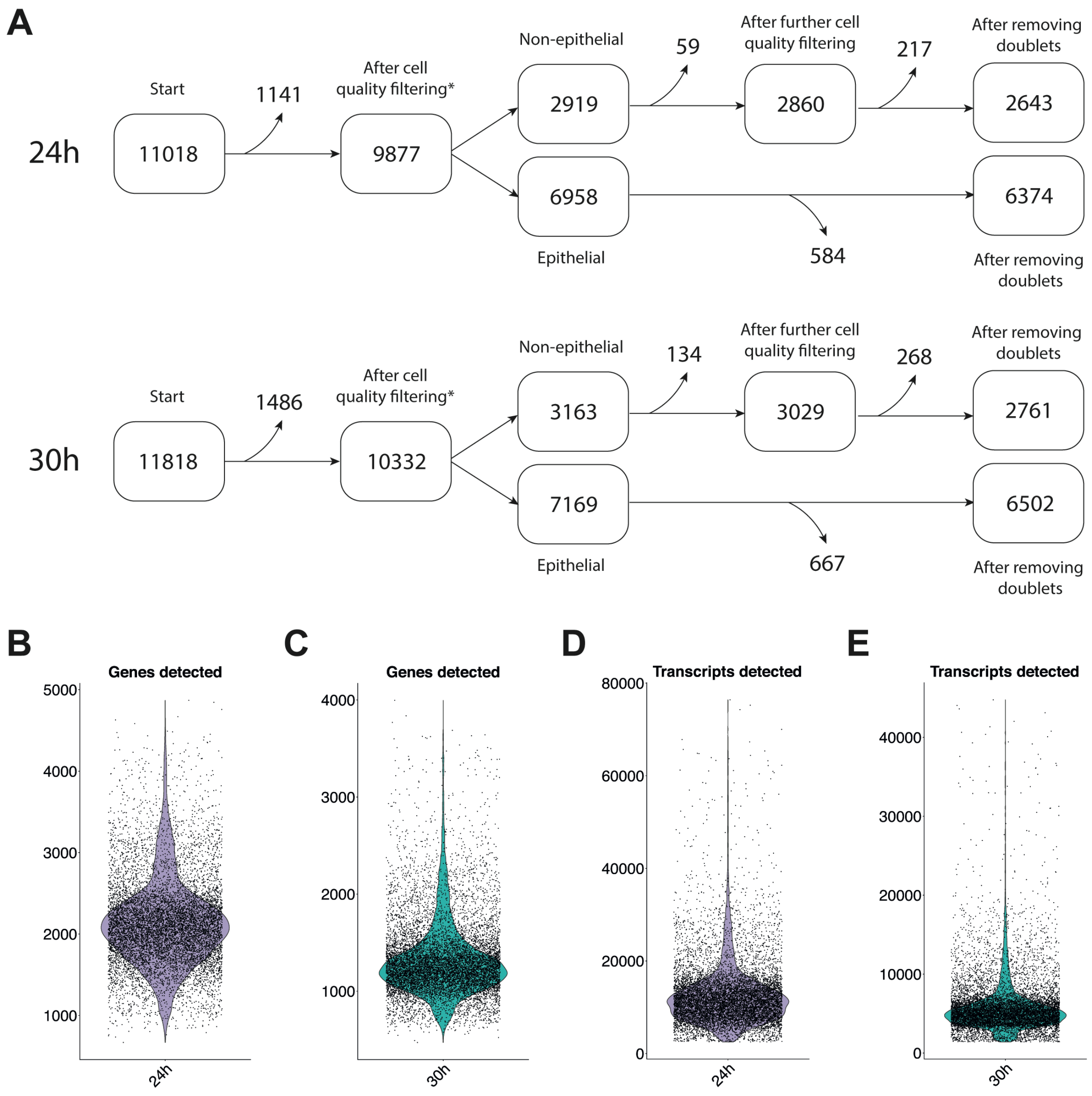


**Figure 2–figure supplement 1. Dataset metrics.**

(A) A schematic detailing how many cells were filtered out at each stage of processing. Cells were initially filtered from the full dataset and retained based on the number of genes detected per cell (24h: >450 & < 5000; 30h: > 425 & < 5000), transcripts detected per cell (24h: > 2500; 30h: > 1400), and the percentage of transcripts that map to mitochondrial genes (24h & 30h: <10%). The datasets were then split into epithelial and non-epithelial cells based on cluster identity (the asterisks here relate to panels (B-E)). Additional filtering was then performed on the non-epithelial cells, removing cells with >5% mitochondrial reads and non-sheath bristle cells in which >2 transcripts of the sheath marker *nompA* were detected, which likely correspond to undissociated doublets. Further doublets were then identified in each dataset using DoubletFinder (McGinnis, Murrow, & Gartner, 2019) and removed. In the 30h dataset, an additional 9 cells positive for the hemocyte marker *NimC4* were identified at the interface between the mechanosensory socket and shaft cluster and at the edge of the bract cluster. These putative hemocyte-bristle cell doublets were also removed.

(B-E) Violin plots showing the distribution of (B) genes detected per cell in the 24h dataset, (C) genes detected per cell in the 30h dataset, (D) transcripts detected per cell in the 24h dataset, and (E) transcripts detected per cell in the 30h dataset. Panels show the distribution in the full datasets after initial filtering based on cell-level quality control metrics (*i.e.,* the distributions at the positions indicated by asterisks in A).

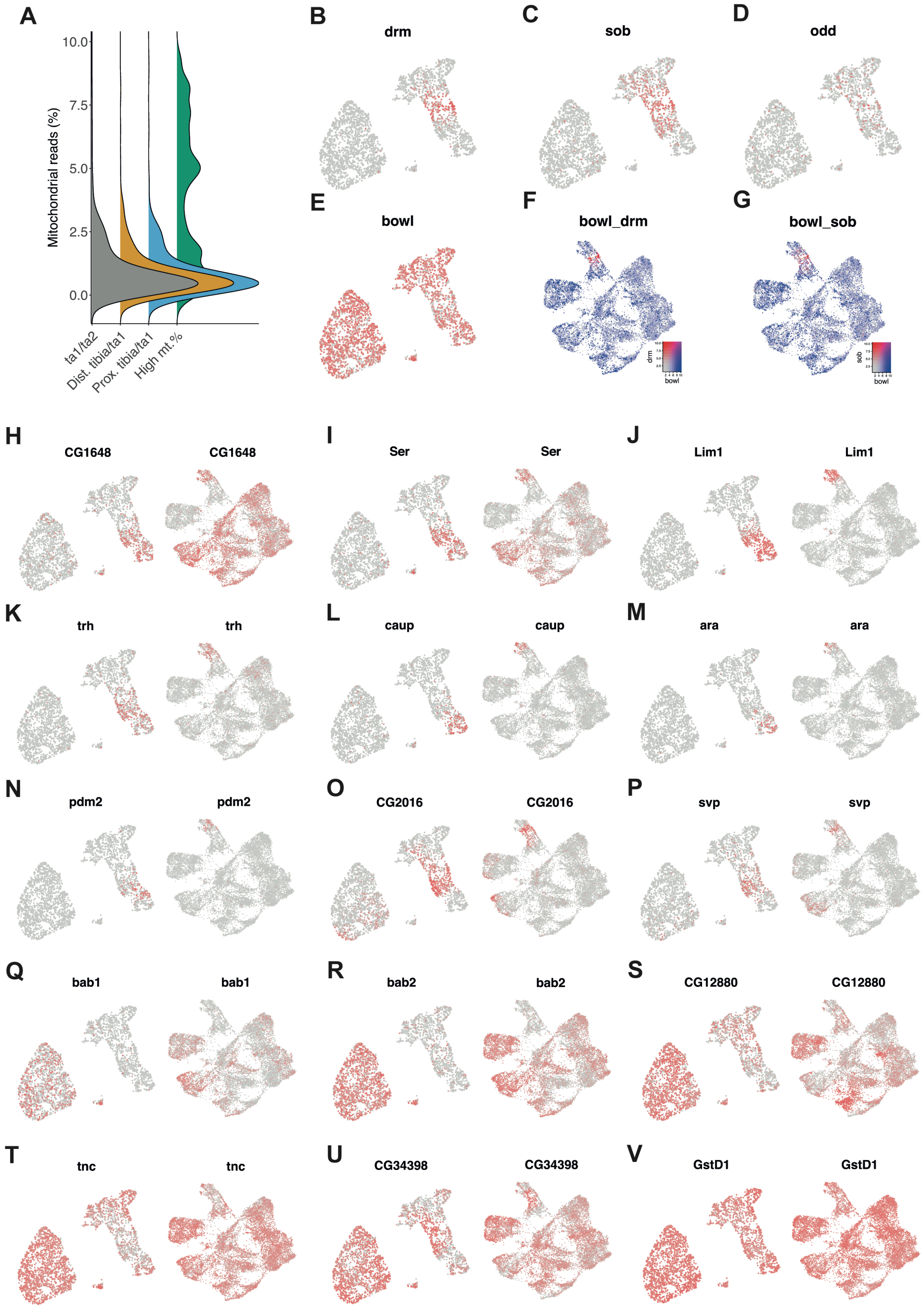


**Figure 3–figure supplement 1. Joint markers.**

**Figure 3–figure supplement 1 (continued)**
(A) The smallest of the clusters identified in our clustering analysis (Figure 3F) showed highly variable internal expression patterns (*i.e.* the top markers of this cluster were generally expressed in a relatively small number of its constituent cells). Moreover, this cluster, coloured dark green in this plot, showed a markedly higher representation of cells with high mitochondrial read counts, suggesting it may be composed of or enriched for damaged cells. We therefore excluded it from further analysis.
(B-E) UMAP plots of the joint dataset overlaid with the expression of *odd-skipped* family transcription factors. *drm, sob,* and *odd* are known to be expressed in the distal edge of each leg segment except tarsal segments 1-4 (Hao *et al.*, 2003). Consistent with its widespread expression among epithelial cells in our dataset, *bowl* has been shown to display an overlapping but broader expression pattern (extending into tarsal segments 1-4) than *odd, drm,* and *sob* (Hao *et al.*, 2003).
(F-G) UMAP plots of the full joint and non-joint epithelial dataset overlaid with the expression of *bowl* (blue) and *drm* (red; F) or *sob* (red; G).
(H-V) For each panel, a UMAP plot of the joint dataset (left) and full joint and non-joint epithelial dataset (right) is overlaid with the expression of a given gene identified during the joint differential gene expression analysis. (H-I) *CG1648* and *Ser* show widespread expression among the proximal tibia/ta1 joint and epithelial cells but are excluded from the other joint clusters. This is expected for *Ser* as *Ser*$^{+}$ cells form patterning boundaries in the developing leg that activate joint formation in distally adjacent cells via Notch (Bishop *et al.*, 1999). (J-N) *Lim1, trh, caup, ara,* and *pdm2* show specific expression in the proximal tibia/ta1 cluster. Of these, *Lim1* is known to be expressed in the tibia where it is required for specification of the tarsus (Tsuji *et al.*, 2000; see also Figure 5V); *nub*$^{1}$ alleles give rise to compromised leg development (Cifuentes & García-Bellido, 1997); *pdm2* shares *cis*-regulatory architecture with *nub*, but, to the best of our knowledge, no role for it in leg development has been characterized (Loker & Mann, 2022); and again, to the best of our knowledge, no roles for the *Iro-C* genes *ara* and *caup* have been reported in leg development.
(O-P) Two of the top DEGs for the proximal tibia/ta1 joint, *CG2016* and *svp,* additionally show enriched expression within a common subregion of what we do not label as joint tissue. Given that this region is *TfAP-2*$^{-}$ it's unlikely that this corresponds to the ta2/ta3 joint and instead more likely corresponds to those cells adjacent to the ta1/ta2 joint. (Q-R) Both *bab* paralogs (*bab1* and *bab2*) are enriched outside of the tibial joint clusters and show particular enrichment within the ta1/ta2 joint cluster, consistent with their known upregulation in the distal portion of the tarsal segments (Godt *et al.*, 1993; Chu, Si Dong, & Panganiban, 2002). (S-V) Several of the top DEGs for the ta1/ta2 cluster (*CG12880*, *tnc, CG34938, GstD1*) do not appear to be specific, rather they show joint-enriched but widespread expression among epithelial cells.

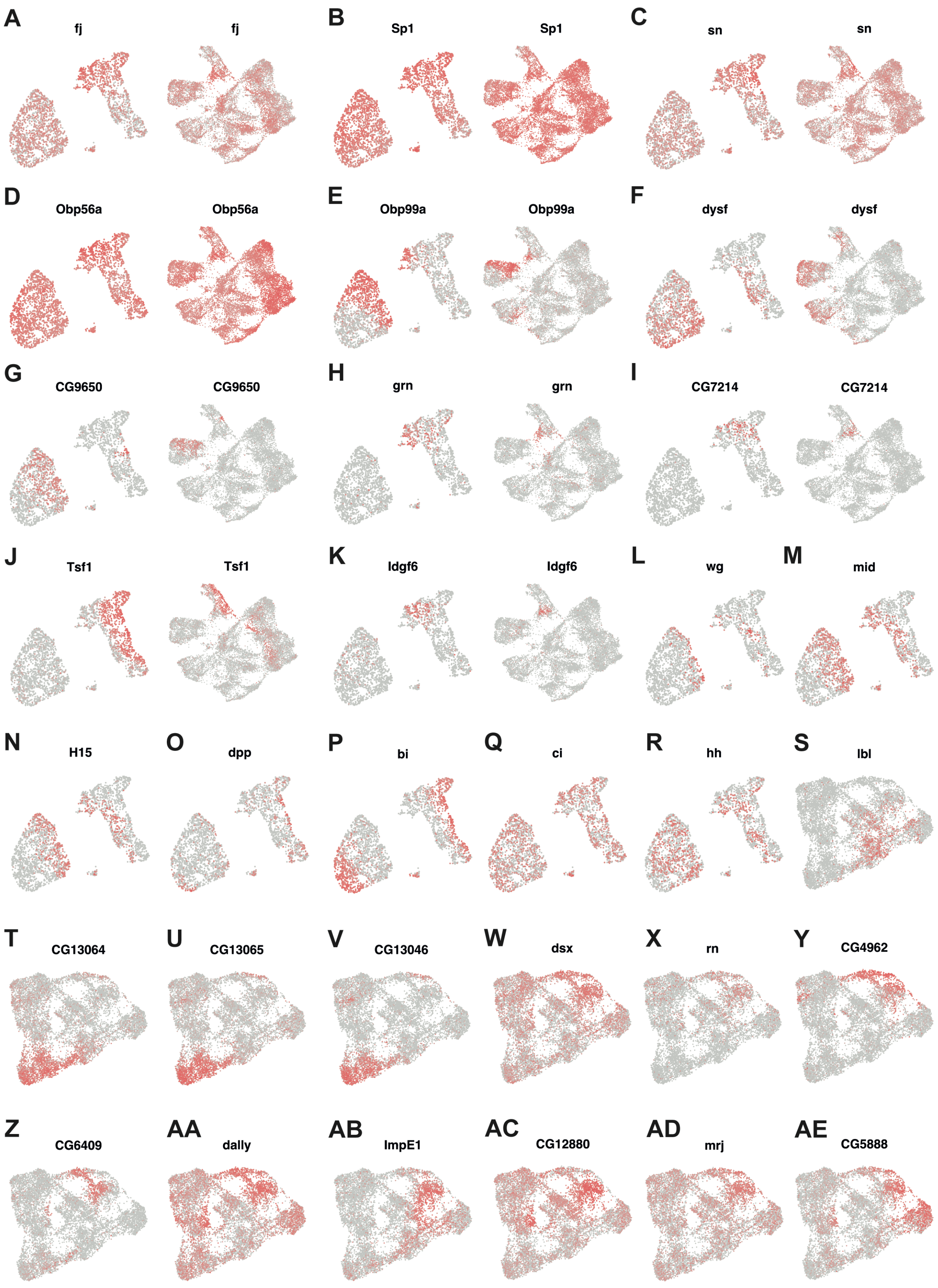


**Figure 3–figure supplement 2. Joint and non-joint epithelial markers.**

**Figure 3–figure supplement 2 (continued)**

(A-K) For each panel, a UMAP plot of the joint dataset (left) and full joint and non-joint epithelial dataset (right) is overlaid with the expression of a given gene identified during the joint cluster differential gene expression analysis. (A-D) Several of the top differentially expressed genes (DEGs) for the distal tibia/ta1 (*fj, Sp1, sn, Obp56a*) do not appear to be specific, rather they show localized enrichment in joint cells alongside widespread expression in non-joint epithelial cells. One such gene is *fj,* which we find widely expressed across all joint and non-joint clusters but enriched in the distal tibia/ta1 cluster (A; Figure 3I). *fj* is known to be required for regional growth along the leg's proximal-distal axis and in imaginal discs shows rings of expression that are complementary to Nub (Villano & Katz, 1995; Rauskolb & Irvine, 1999). However, our data show a separation between the regions of peak *nub* and *fj* expression (compare Figure 3I with 3D). Several of the remaining top DEGs for the ta1/ta2 (E-G) and distal tibia/ta1 (H-K) joint clusters show more specific expression patterns.

(L-R) UMAP plots of the joint dataset overlaid with the expression of genes specifying positional identity. Positional identity is clear at the level of the dorsal-ventral axis (Ventral: *wg, mid, H15;* Dorsal: *dpp, bi*; L-P), but not anterior-posterior axis (Anterior: *ci*; Posterior: *hh*; Q-R).

(S-AE) UMAP plots of the non-joint epithelial dataset overlaid with the expression of genes identified as enriched within subregions.

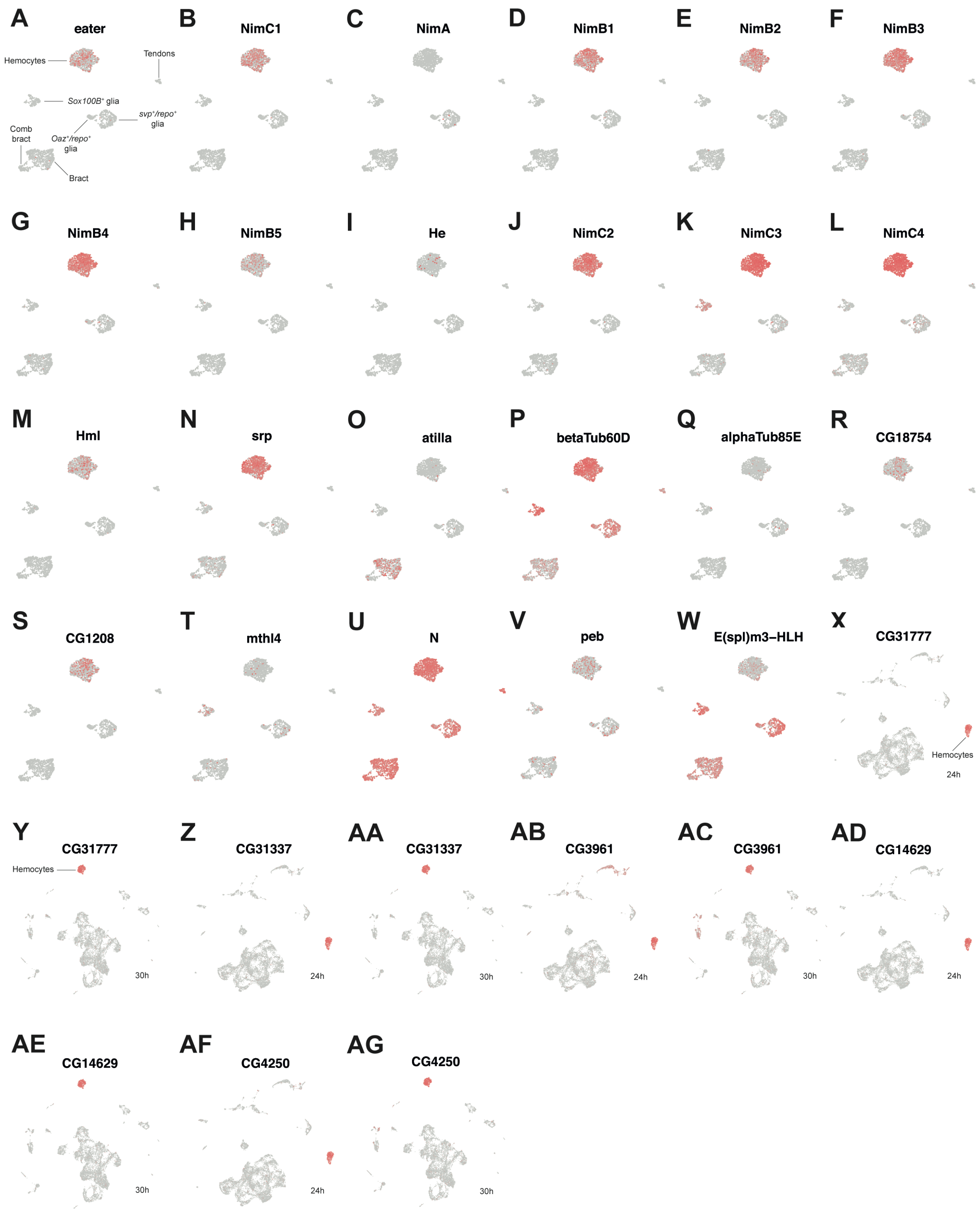


**Figure 4–figure supplement 1. Pupal leg hemocytes form a uniform population.**

**Figure 4–figure supplement 1 (continued)**
(A-W) UMAP plots of the non-sensory dataset overlaid with gene expression. (A-N) A selection of hemocyte markers, most of which are part of a cluster of NIM-repeat containing genes on chromosome 2 that also includes the hemocyte-specific *He.* RT-PCR work has previously shown that all of these, except *nimA*, are transcribed in larval hemocytes (Kurucz *et al.*, 2007). We find that all these NIM genes, except *nimA*, are expressed in our hemocyte cluster, as are *Hml* and *srp,* which are known to be expressed in both differentiating and mature plasmatocytes (Evans, Hartenstein, & Banerjee, 2003; Kocks *et al.*, 2005). (O-T) Genes identified by Tattikota *et al.* (2020; see also Hultmark & Andó, 2022) as enriched in lamellocytes. We saw no obvious subclustering in relation to these genes: they were either widely expressed among hemocytes (*betaTub60D*, *CG1208*), too patchily expressed among hemocytes to reflect a clear subpopulation (*atilla, alphaTub85E, CG18754, mthl4*), or absent from our dataset entirely (*CG31219, CG14610, CG15347, CG12133*). (U-W) The same study found that crystal cells showed highest enrichment of *PPO1, PPO2, lz, N, peb,* and *E(spl)m3-HLH.* As with lamellocytes, we saw no obvious subclustering in relation to these genes: *PPO1, PPO2,* and *lz* were absent from our dataset, and *N, peb,* and *E(spl)m3-HLH* showed non-specific expression. We also mapped the top markers of many of the plasmatocyte subclusters identified by Tattikota *et al.* (2020)(*Mmp1, IM18, CecA2, CecC, Mtk, DptB, Drs, Prx2540-1, Prx2540-2, CG12896, Abl, Snoo, CG15550, CG6023, mthl7, Cys, CG8860,* and *COX8*; data not shown), but saw no clear subclustering.
(X-AG) Alternating UMAP plots of the full 24h (X, Z, AB, AD, AF) and 30h (Y, AA, AC, AE, AG) datasets overlaid with expression of a subset of hemocyte marker genes identified in this study.

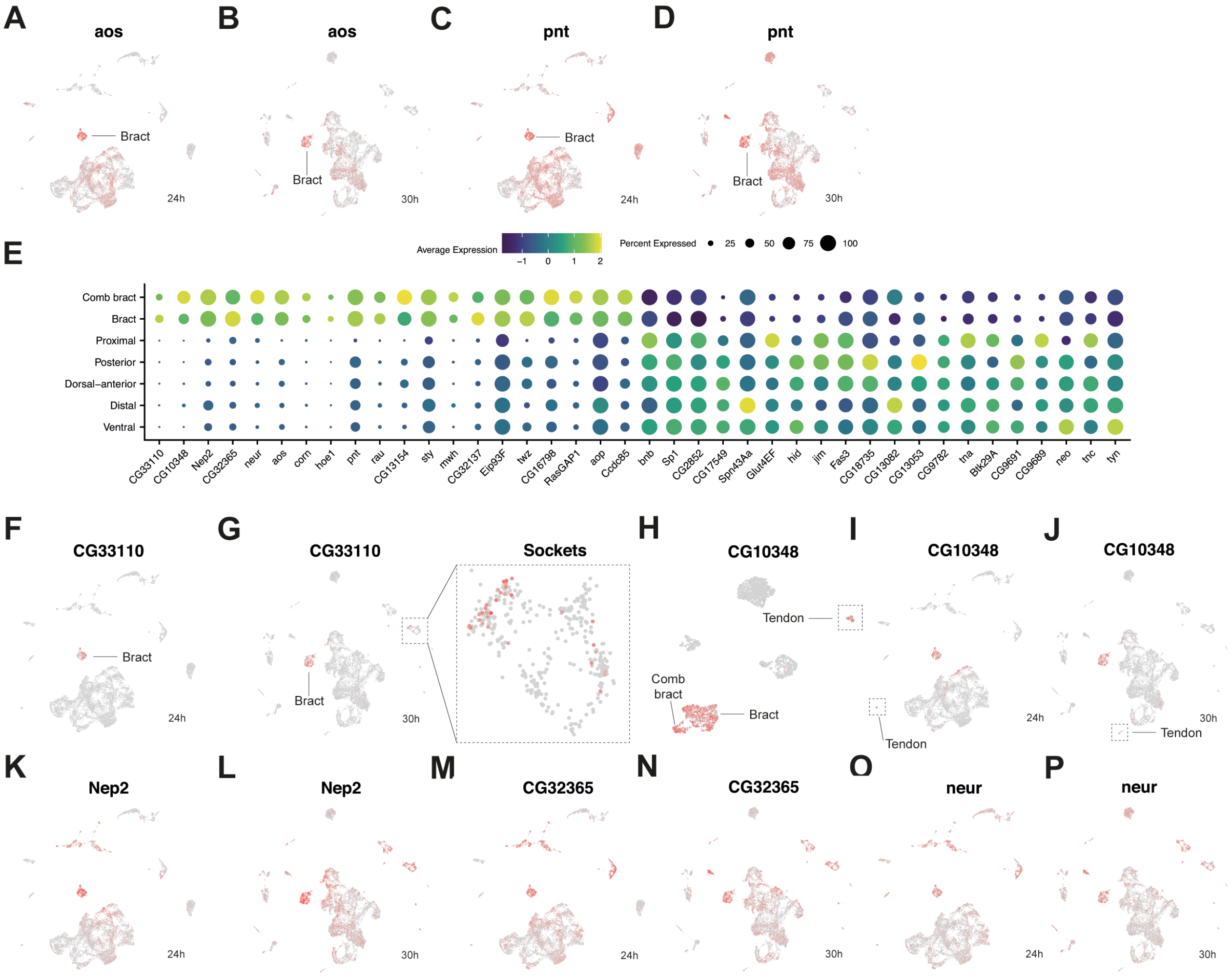


**Figure 4–figure supplement 2. Induction of bract cell identity in epithelial cells is accompanied by a remodeling of the transcriptome.**

(A-D) Alternating UMAP plots of the full 24h (A,C) and 30h (B,D) datasets overlaid with expression of the bract markers *pnt* and *aos,* EGFR signalling components with known roles in bract formation (del Álamo, Terriente, & Díaz-Benjumea, 2002; Peng, Han, & Axelrod, 2012; Mou *et al.*, 2012).

(E) A dot plot of the top 20 differentially expressed genes from a bract versus non-joint epithelial comparison. Although the comparison was made between all bracts and all non-joint epithelial cells, the breakdown per subcluster is depicted (‘distal’ here refers to the sex comb bearing region). Of these, the ecdysone-induced transcription factor *Eip93F* (also called *E93*), which we found to be upregulated in bract cells, is known to be expressed in the epithelial cell that will develop as a bract, where it enables *Dll* to respond to EGFR signalling (Mou *et al.*, 2012).

(F) UMAP plot of the full 24h dataset overlaid with the expression of the top bract marker, *CG33110,* which encodes a predicted fatty acid elongase.

(G) As (F) but in the 30h dataset, alongside an inset zooming in on the socket cell cluster. The expression of *CG33110* in sockets was more pronounced in the 30h compared to 24h dataset.

(H) A UMAP plot of the non-sensory dataset overlaid with expression of *CG10348*, a top bract marker. Expression is clearly enriched in bract cells, as well as tendon cells, which are highlighted by a dashed box.

(I-P) Alternating UMAP plots of the full 24h (I,K,M,O) and 30h (J,L,N,P) datasets overlaid with several top bract markers. In the case of most of these genes, expression is detected outside the bracts and often enriched in the sensory organ cells.

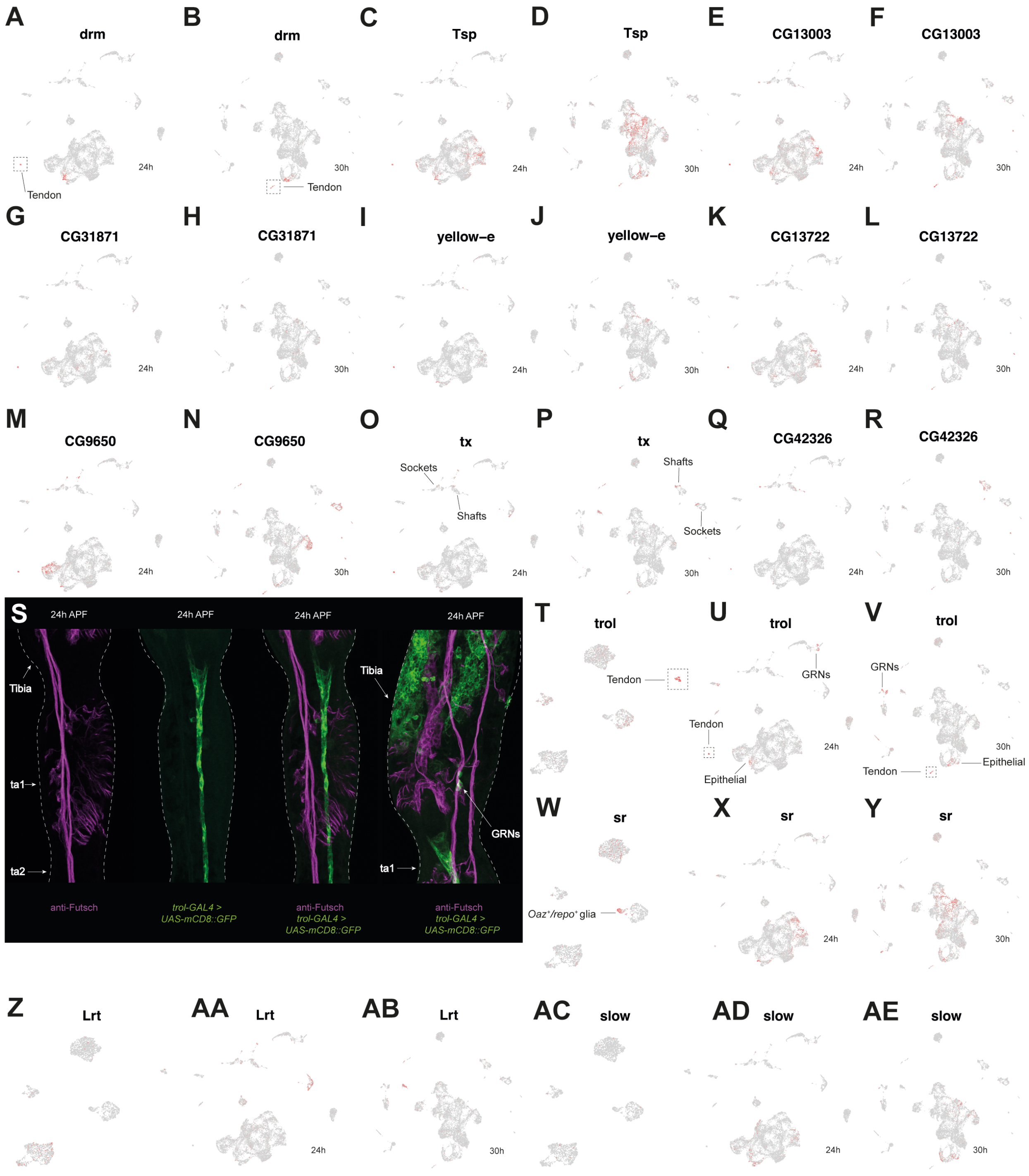


**Figure 4–figure supplement 3. Characterizing the tendon transcriptome**

**Figure 4–figure supplement 3 (continued)**

(A-R) Alternating UMAP plots of the full 24h (A,C,E,G,I,K,M,O,Q) and 30h (B,D,F,H,J,L,N,P,R) datasets overlaid with several top tendon markers. (A-N) For *drm, Tsp, CG13003, CG31871, yellow-e, CG13722,* and *CG9650,* expression was observed in both tendon cells and a subregion of epithelial cells. For *tx* (O,P) and *CG42326* (Q,R), we also observed expression in a subset of shaft and socket cells. In both cases, this socket and shaft expression was more widespread at 30h.

(S) Confocal images of 24h APF male *trol-GAL4 > UAS-mCD8::GFP* (green) (Li *et al.*, 2017) legs counterstained with the neuronal marker anti-Futsch (magenta). The first three images show the separate and merged channels from an image of the first tarsal segment. The staining follows much the same pattern showed by anti-Vvl and *1151-GAL4 > UAS-mCherry.nls* in the tendon cells (Figure 4G-J). Note the concentration of staining around the tibia/ta1 joint, the position of the levator and depressor tendons. The right-hand image shows the distal tibia and proximal ta1 with merged channels. Note the presence of extensive *trol-GAL4* staining in epithelial cells in the tibia – no equivalent epithelial staining was observed in the tarsus. The epithelial *trol-GAL4* staining observed in the tibia was not present in the region proximal to the tibia/ta1 joint. *trol-GAL4* staining was also observed in gustatory receptor neurons (GRNs) (see Figure 7–figure supplement 3).

(T-V) Expression of *trol* overlaid on the non-sensory (T), full 24h (U), and full 30h (V) UMAP plots. Note the expression of *trol* in a subset of GRNs (as shown in (S) and Figure 7–figure supplement 3). Some localized expression is present in a region of the epithelial clusters that corresponds to the proximal tibia/ta1 portion of our joint UMAPs, rather than the distal tibia/ta1 region, and therefore likely reflects the tibia/ta1 joint staining rather than that in the more distal tibia, which falls outside of our dissected region.

(W-AE) Alternating UMAPs showing expression of known tendon genes *sr, Lrt,* and *slow* overlaid on the non-sensory (W,Z,AC), full 24h (X,AA,AD), and full 30h (Y,AB,AE) datasets. Note the enriched expression of *sr* in a non-tendon region of the non-sensory UMAP (W). These cells correspond to the *Oaz*$^{+}$/*repo*$^{+}$ glia.

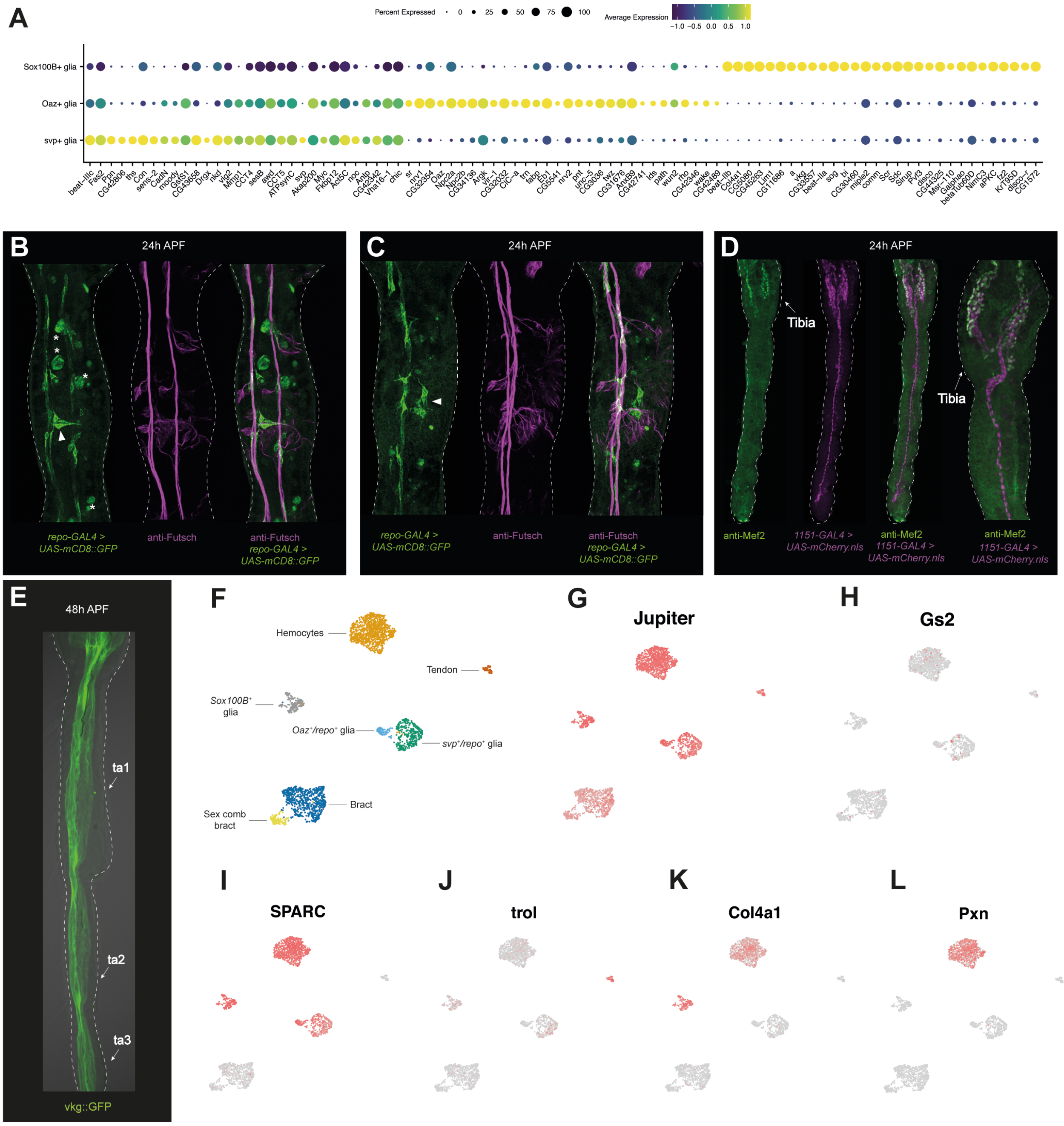


**Figure 5–figure supplement 1. Annotation of glia clusters.**

(A) A dot plot of the top differentially expressed genes from between-glia comparisons (*e.g., Oaz*$^+$*/repo*$^+$ glia *vs. svp*$^+$*/repo*$^+$ and *Sox100B*$^+$ glia, *etc.*).

(B-C) 24h APF male first tarsal segments from *repo-GAL4 > UAS-mCD8::GFP* (green) counterstained with anti-Futsch (magenta). Note how the staining pattern includes cells hugging the axon trunks, but also, and unlike in the *Lim1-GAL4* staining, cells that branch away from them. This latter feature is particularly clear around the sex comb (see white triangles). Note also that some circular, non-specific fat body staining is present (examples marked by asterisks in (B)).

(D) 24h APF male forelegs from *1151-GAL4 > UAS-mCherry.nls* (magenta) males counterstained with anti-Mef2 (green). *1151-GAL4* is a myoblast and tendon cell marker while Mef2 is a transcription factor essential for cardiac, visceral, and somatic muscle development (Lilly *et al.,* 1995). Anti-Mef2 staining is restricted to the tibia.

(E) Confocal image of a male 48h pupal leg from a *vkg::GFP* line. Both transmitted light and GFP channels are shown merged.

(F) Annotated UMAP plot of non-sensory cells.

(G-L) The UMAP shown in (F) overlaid with the expression of: (G) *Jupiter,* a perineural glia marker (Xie & Auld, 2011) that here shows widespread expression; (H) *Gs2,* a surface glia marker that here shows patchy expression in the *svp*$^+$*/repo*$^+$ cluster (Sasse & Klämbt, 2016). (I-L) A series of extracellular matrix components that show varying expression profiles across *repo*$^+$ glia, *Sox100B*$^+$ glia, and hemocytes.

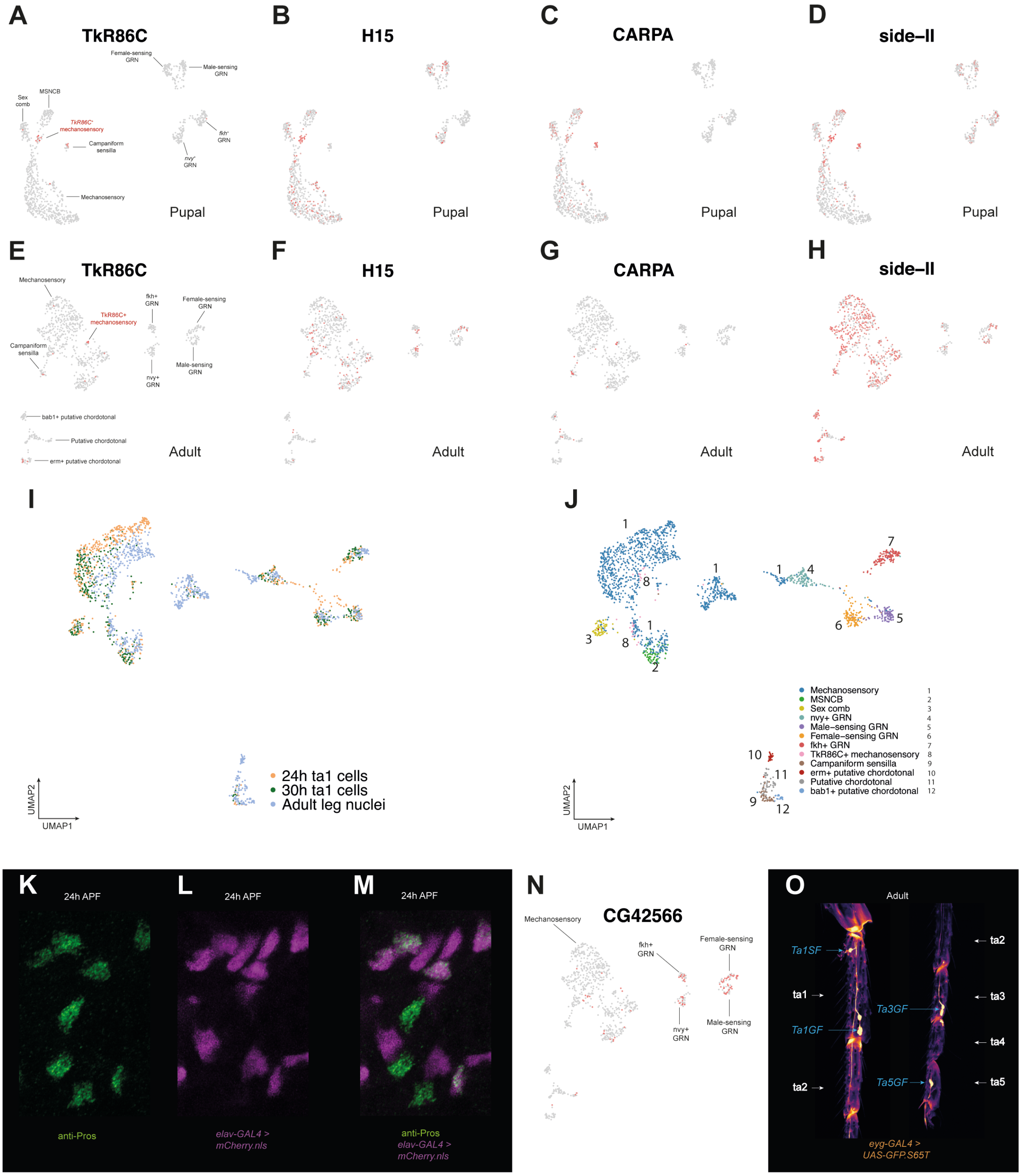


**Figure 6–figure supplement 1. Annotation of sensory neuron populations**

(A-H) In our unsupervised clustering analysis of both the pupal neuron and the FCA neuron datasets, a small subpopulation of mechanosensory neurons clustered separately, labelled in (A) and (E) as *TkR86C*$^{+}$ mechanosensory neurons. These clusters were enriched for *TkR86C* (A,E)*,* which encodes a receptor for the neuropeptide tachykinin and plays a critical role in male-specific neural circuits that control aggression (Asahina *et al.*, 2014). (B-D) In the pupal dataset, these cells were also enriched for the ventral marker *H15*, *CARPA*, and *Side-II.* (F-H) However, in the adult data only *CARPA* showed any suggestion of being enriched in these cells relative to the other neuron populations. Because of their scarcity, coupled with the absence of strongly specific genes, it's unclear whether the *TkR86C*$^{+}$ cluster represents a distinct population. We cannot rule out that in the pupal data they may simply correspond to more developmentally advanced mechanosensory neurons and/or a population from a particular subregion of the leg, the clustering of which is driven by the shared expression of positional markers such as *H15*.

(I) A UMAP plot of an integrated dataset of the 24h APF male first tarsal segment single-cell RNA-seq data, 30h APF male first tarsal segment single-cell RNA-seq data, and adult all leg male neuron single-nuclei RNA-seq data. Cells are coloured according to the dataset of origin.

**Figure 6–figure supplement 1 (continued)**

(J) The UMAP plot given in (I) but this time cells are coloured according to the cluster annotation they were assigned based on separate clustering and analysis of the pupal cell and adult nuclei datasets (*i.e.* those presented in Figure 6A and G). Note how the *nvy*$^+$ cluster includes both cells labelled as *nvy*$^+$ GRNs and mechanosensory neurons. Because of this divergent classification, and the broader differences between the datasets in the tissues, their ages, and the dissociation protocol used, we opted to analyze the two datasets separately.

(K-M) Confocal images of mechanosensory bristles in a subregion of the first tarsal segment from *elav-GAL4 > UAS-mCherry.nls* (magenta) males, counterstained with anti-Pros (green). The channels are shown separately and merged. Note the presence of both Pros$^+$/*elav-GAL4*$^+$ and Pros$^+$/*elav-GAL4*$^-$ cells. This heterogeneity likely reflects between-bristle variation in the developmental stage of mechanosensory sheath cells such that soon after division mechanosensory sheaths are *elav-GAL4*$^+$(see also Simon *et al.*, 2019). *elav-GAL4* expression is later lost from mechanosensory sheaths, restricting it to neurons.

(N) A UMAP plot of the Fly Cell Atlas single-nuclei adult male leg neuron data overlaid with the expression of *CG42566.* Expression of *CG42566* is largely restricted to the four gustatory receptor neuron (GRNs) classes. In the pupal data, it is also expressed in a subpopulation of *vvl*$^-$ mechanosensory neurons, which we presume to correspond to the mechanosensory neurons that innervate chemosensory bristles (MSNCBs). It's possible that the patchy expression of *CG42566* in the mechanosensory neuron cluster observed here in the adult data reflects MSNCBs interspersed among mechanosensory neurons. Interestingly, *CG42566* is present in *nvy*$^+$ GRNs in the adult data but absent from them in the pupal data, which may reflect between-GRN developmental timing differences.

(O) Confocal images of adult male *eyg-GAL4 > UAS-GFP.S65T* legs. On the left, the first two tarsal segments (ta1 and ta2) are shown; on the right, the second through to the fifth (ta2-ta5). Blue arrows and text denote the name and position of visible, stained campaniform sensilla. The naming follows the nomenclature of Dinges *et al.* (2021). Under this nomenclature, the first three characters denote the tarsal segment; the fourth, whether the sensilla are found singly (S) or in a group (G) of two (as in distal ta1 and ta3) or more (as in ta5); and the fifth, that they are on the front (F) leg. In Ta1GF, note how the cell body of the neuron sits a little further back from the accessory cells, a position it may adopt due to the rotation of the sex comb.

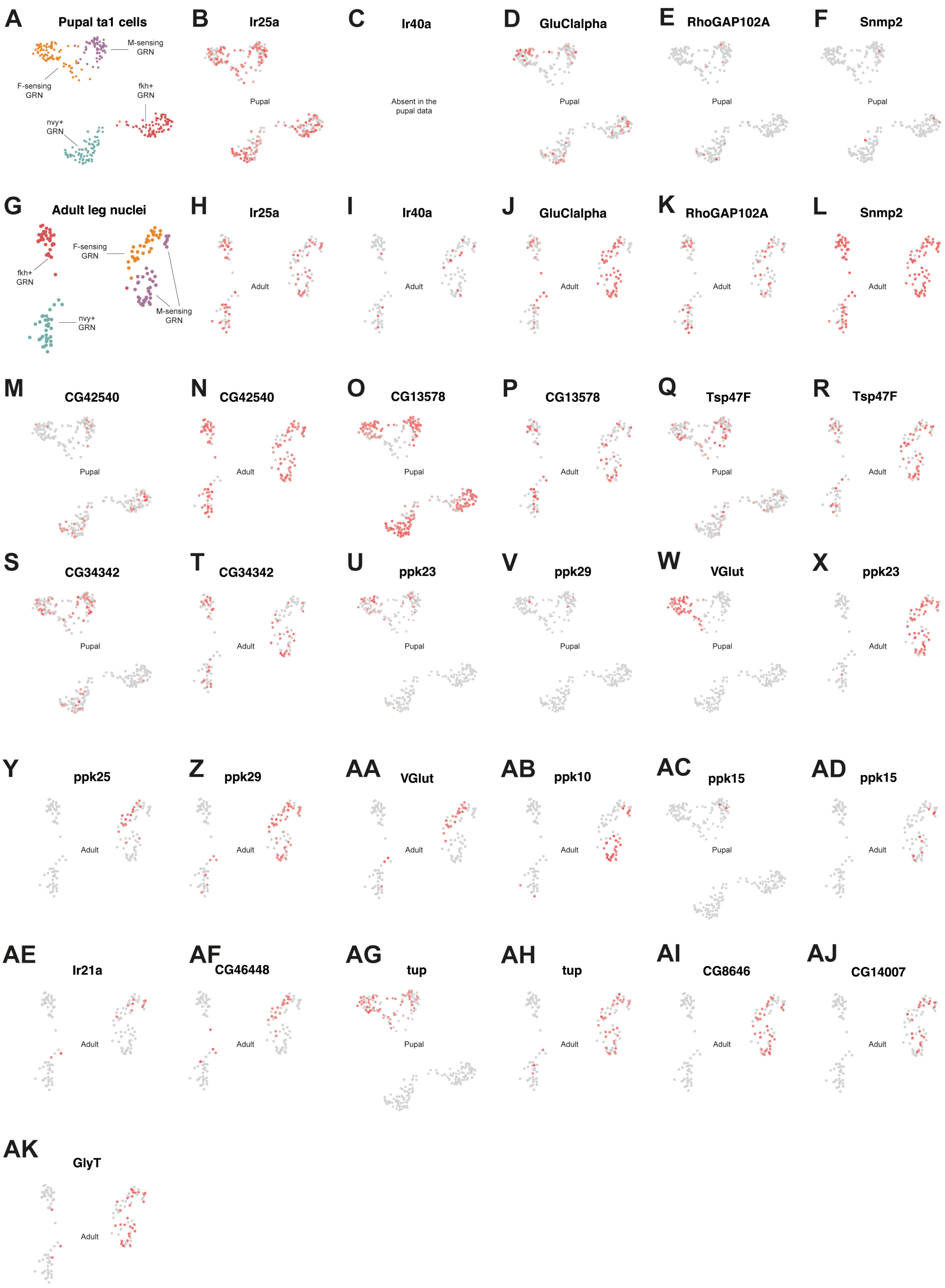


**Figure 7–figure supplement 1. Gene expression profiles of gustatory receptor neurons I**

**Figure 7–figure supplement 1 (continued)**

(A-T) UMAPs showing the annotated GRN (gustatory receptor neuron) clusters identified in the integrated pupal neuron data (A) and FCA adult leg data (G) and then overlaid with the expression of genes identified as being specifically expressed or enriched in all GRNs relative to all other neuron populations.

(U-AK) The expression of a set of genes that are enriched in the female-sensing and/or male-sensing GRN clusters overlaid on the UMAPs described in (A) and (G). Of these, *ppk23* and *ppk29* are known from previous work to be expressed in both neuron types, while *VGlut* is restricted to female-sensing neurons (Lu *et al.*, 2012; Starostina *et al.*, 2012; Thistle *et al.*, 2012; Toda, Zhao, & Dickson, 2012; Vijayan *et al.*, 2014; Kallman, Kim, & Scott, 2015). We additionally detect the transcription factor *tup* in both populations and *ppk10* and *ppk15* in the male-sensing population. Note that in (AD), 2 of the 3 *ppk15*$^{+}$ cells outside the main body of M-sensing GRN cells fall within the *acj6*$^{+}$ region we identified as likely being additional M-sensing cells (see Figure 7AH). Taking the two datasets together, *ppk15* represents a strong candidate for a gene involved in male pheromone detection, as does *ppk10.* A gene plotted for just one of the two datasets indicates its absence from the other.

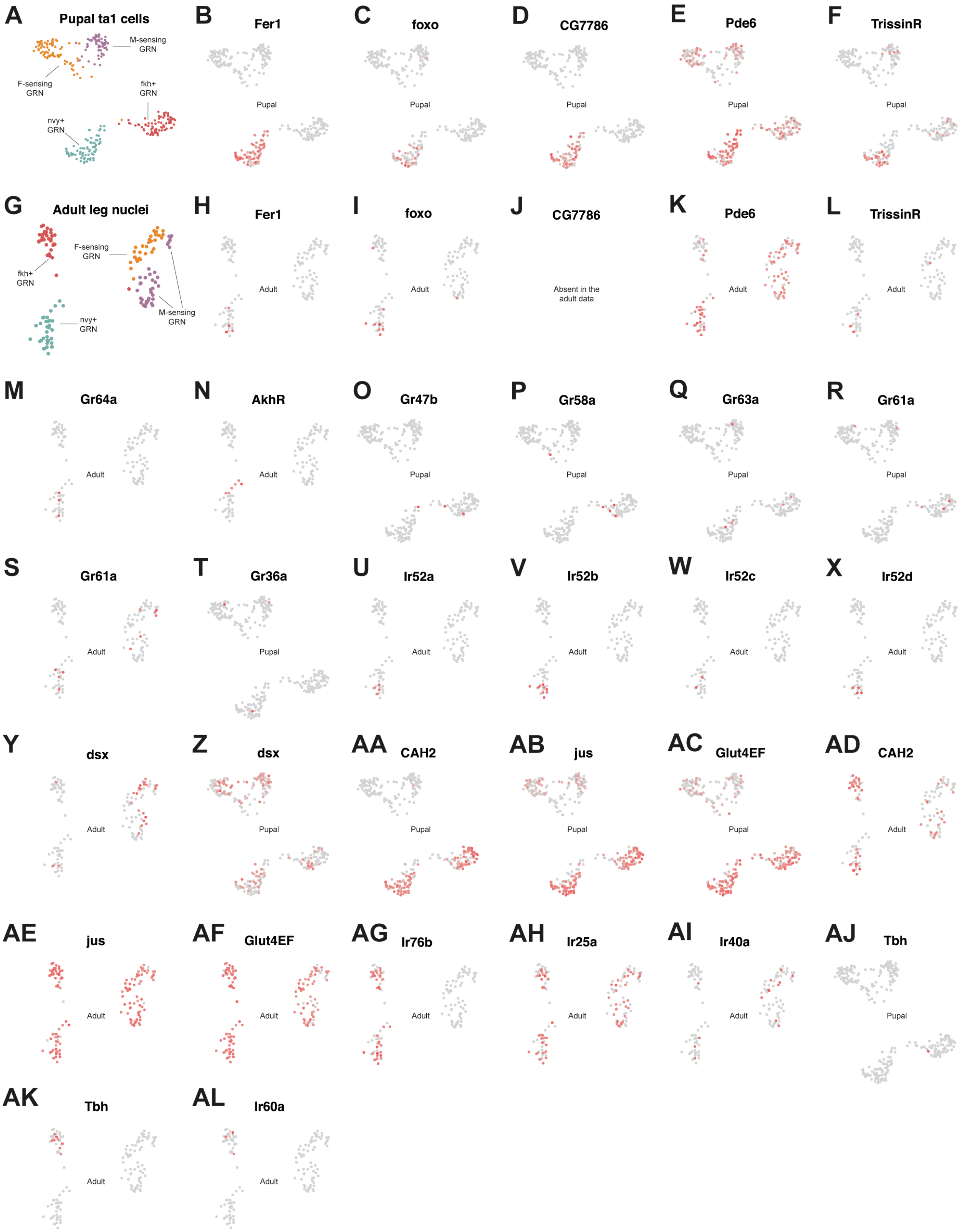


**Figure 7–figure supplement 2. Gene expression profiles of gustatory receptor neurons II**

**Figure 7–figure supplement 2 (continued)**
(A-N) UMAPs showing the annotated GRN (gustatory receptor neuron) clusters identified in the integrated pupal neuron data (A) and FCA adult leg data (G) and then overlaid with the expression of genes identified as being specifically expressed or enriched in the *nvy*$^+$ GRN cluster.
(O-T) As well as searching through DGEs, we ran through the expression of all 60 gustatory receptor genes (Clyne, Warr, & Carlson, 2000; Scott *et al.*, 2001; Dunipace *et al.*, 2001; Robertson, Warr, & Carlson, 2003; Ling *et al.*, 2014). Of these, we detected *Gr36a, Gr47b, Gr58a,* and *Gr63a* exclusively in the pupal ta1 dataset, *Gr64a* exclusively in the adult leg dataset, and *Gr61a* in both. In all cases, expression was limited to just a handful of cells precluding us from confidently assigning *Grs* to specific GRNs. We failed to detect *Gr68a,* a receptor that has been shown to be specifically expressed in the male foreleg (Bray & Amrein, 2003), in either dataset.
(U-X) *Ir52a-d*, the expression of which is shown here overlaid on the UMAP presented in (G), were specifically expressed in the *nvy*$^+$ GRN cluster and only detected in the adult dataset.
(Y-Z) The expression of *dsx* appears more widespread among the GRNs in the pupal data compared to the adult data. This may reflect differences in the dissected regions, with *dsx* expression restricted to only the *nvy*$^+$ and *fkh*$^+$ GRNs on the foreleg, while *dsx* is expressed in all *fru*$^+$ neurons regardless of the leg they're on.
(AA-AL) A selection of genes enriched specifically in *fkh*$^+$ GRNs or that are shared between *fkh*$^+$ and *nvy*$^+$ GRNs. Note how initially restricted expression in the pupal data of *CAH2* (AA), *jus* (AB), and *Glut4EF* (AF) expands to widespread expression in the adult data (AD-AF). Genes presented for just one of the two UMAPs indicates its absence from the other dataset.

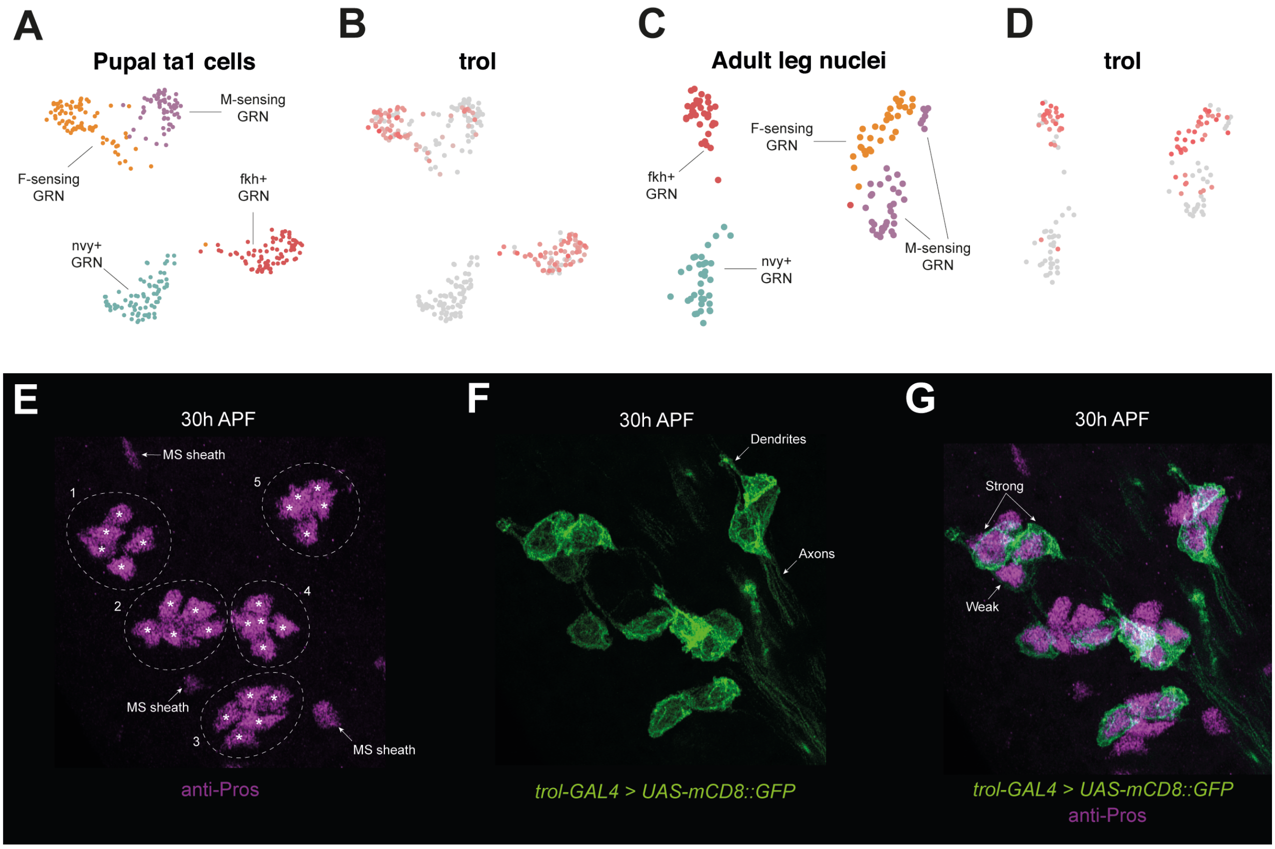


**Figure 7–figure supplement 3. Variable expression of the extracellular matrix proteoglycan gene *trol* between GRN classes.**

(A) A UMAP showing the annotated GRN (gustatory receptor neuron) clusters identified in the integrated pupal neuron data.

(B) The UMAP shown in (A) overlaid with the expression of *trol*, which encodes the extracellular matrix proteoglycan Perlecan.

(C) A UMAP showing the annotated GRN clusters identified in the Fly Cell Atlas single-nuclei adult male leg neuron data.

(D) The UMAP shown in (C) overlaid with the expression of *trol*.

(E-G) Confocal images of 30h APF male first tarsal segments from *trol-GAL4 > UAS-mCD8::GFP* (green) counter-stained with anti-Pros (magenta). We imaged at 30h APF because at 24h APF *trol-GAL4* expression in the bristles of the first tarsal segment was either undetectable or very weak. However, at 24h APF *trol-GAL4* expression was clear in the more distal segments, suggesting these bristles may develop ahead of those in the more proximal segments. The imaged region contains 5 chemosensory bristles, which are circled and numbered in (E). anti-Pros marks 5 nuclei (marked with an asterisk) in each bristle: 4 correspond to GRNs and 1 to the sheath. Individual $Pros^+$ nuclei are visible outside of the circled chemosensory bristles shown in (E). These correspond to the sheath cells of mechanosensory (MS) bristles. In (G), note how there is between-bristle variation in the number of *trol-GAL4*$^+$ cells. Bristle 1: 2 strongly positive, 1 weakly positive, 2 negative; Bristle 2: 2 positive, 3 negative; Bristle 3: 2 positive, 3 negative; Bristle 4: 3 positive, 2 negative; Bristle 5: 3 positive, 2 negative. This variability matches the expression profile shown in the UMAPs (B,D), where F-sensing and *fkh*$^+$ GRNs show strong *trol* expression and M-sensing GRNs show variable expression.

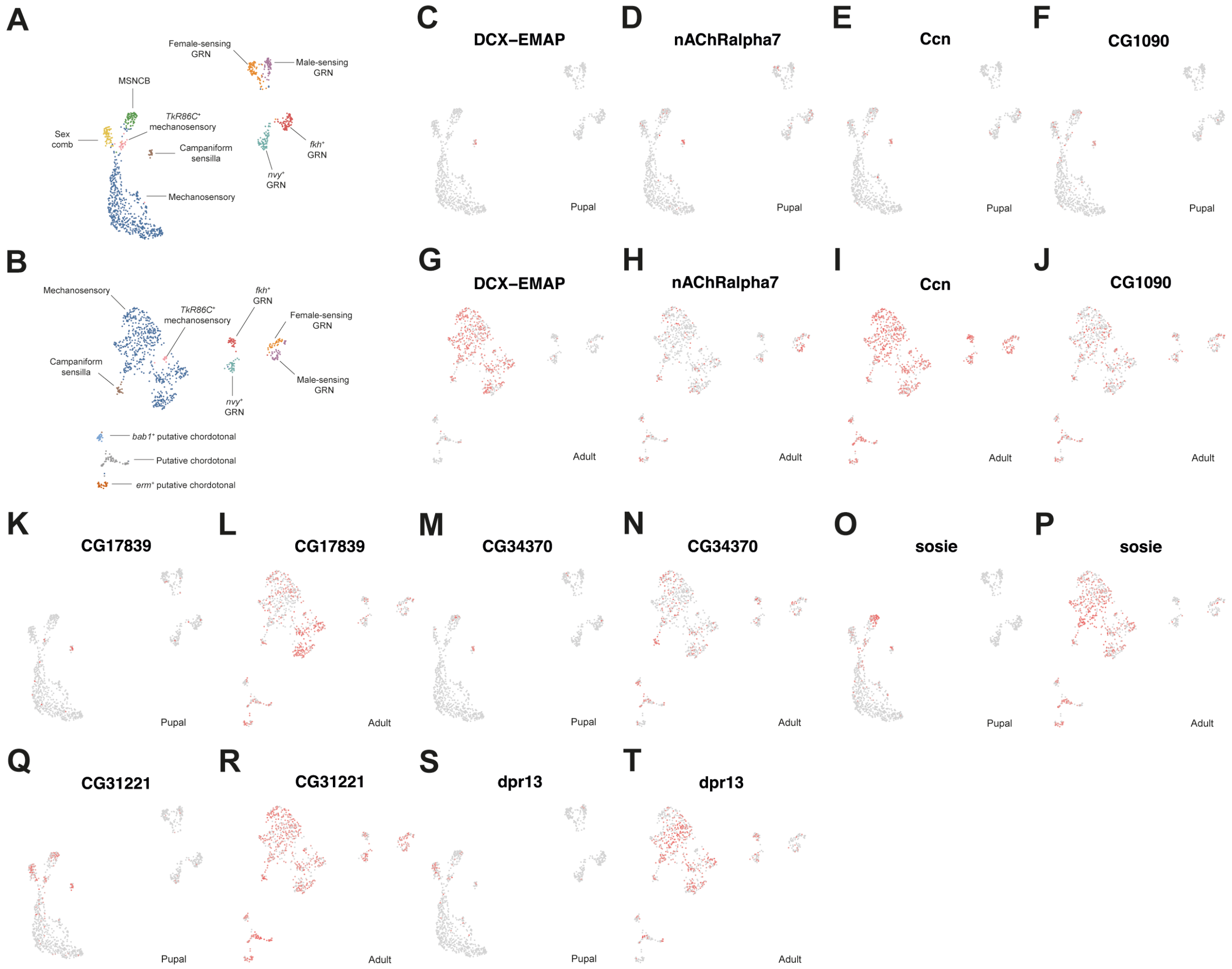


**Figure 7–figure supplement 4. Heterochronic differences between sensory neuron classes.**

(A) Annotated UMAP of the pupal integrated neuron data. GRN = gustatory receptor neuron; MSNCB = Mechanosensory neuron in chemosensory bristle.

(B) Annotated UMAP of male neuronal cells subsetted from the Fly Cell Atlas single nuclei RNA-seq leg dataset (Li *et al.*, 2022).

(C-N) The UMAPs described in (A) and (B) overlaid with a selection of genes identified among the top markers of campaniform sensilla neurons in the pupal data and which show a loss of specificity when moving from the pupal to adult dataset.

(O-T) The UMAPs described in (A) and (B) overlaid with a selection of genes identified among the top markers of MSNCBs and/or sex comb neurons in the pupal data and which show a loss of specificity when moving from the pupal to adult dataset.

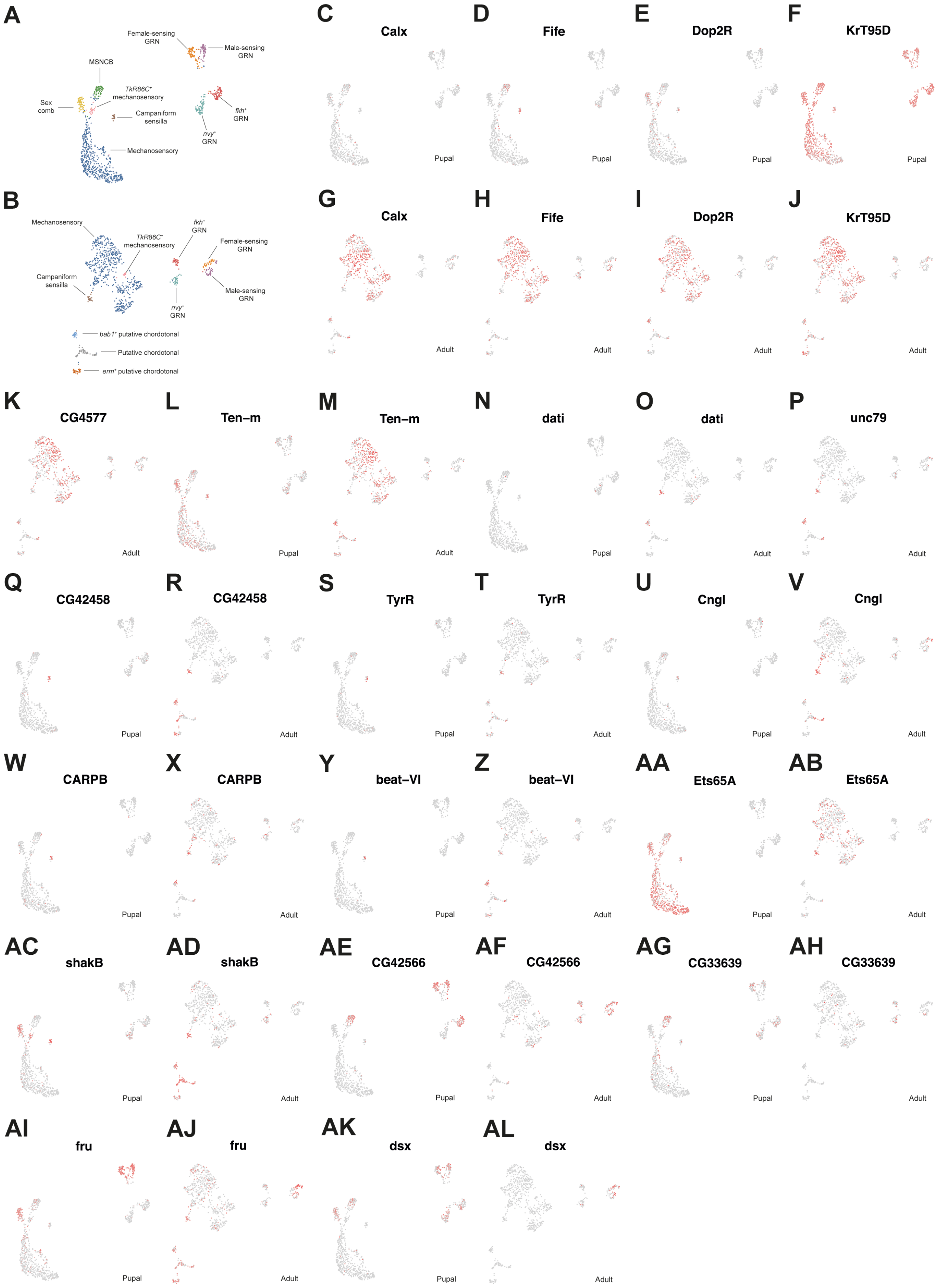


**Figure 7–figure supplement 5. Gene expression in external mechanotransduction neurons.**

**Figure 7–figure supplement 5 (continued)**
(A) Annotated UMAP of the pupal integrated neuron data. GRN = gustatory receptor neuron; MSNCB = Mechanosensory neuron in chemosensory bristle.
(B) Annotated UMAP of male neuronal cells subsetted from the Fly Cell Atlas single nuclei RNA-seq leg dataset (Li *et al.*, 2022).
(C-AJ) The UMAPs described in (A) and (B) overlaid with a selection of genes showing enriched expression in the different external sensory organ neuron classes involved in mechanotransduction, namely mechanosensory neurons, MSNCBs, sex comb neurons, and campaniform sensilla. (A-M) Genes identified as top markers of mechanosensory neurons in the adult FCA data, but all show expression in other populations. (N-Z) Genes identified as top markers of campaniform sensilla neurons in the adult FCA dataset. Note how many are also expressed in chordotonal organ populations, but few or no mechanosensory neuron populations. (AA-AB) Across both datasets, *Ets65A* appears largely restricted to mechanosensory neurons, MSNCBs, sex comb neurons, and campaniform sensilla. (AC-AD) Although relatively widely expressed in the adult data, *shakB* show marked enrichment in sex comb neurons in the pupal data. (AE-AH) *CG42566* and to a lesser extent *CG33639* appear enriched in MSNCBs. (AI-AL) The two effectors of sexual differentiation, *fru* and *dsx*, show distinct expression profiles from one another.

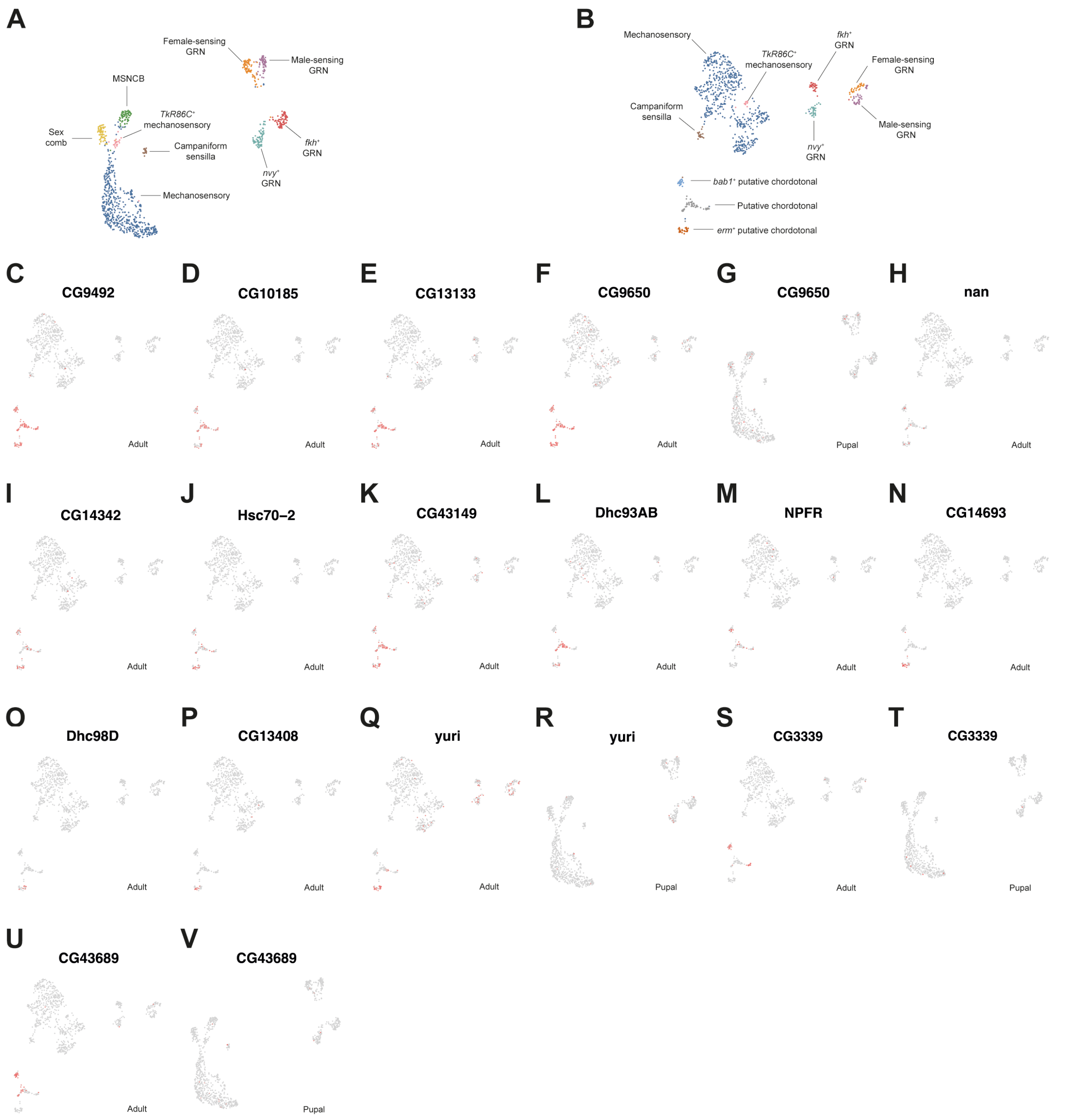


**Figure 7–figure supplement 6. Gene expression in putative chordotonal organ neurons.**

(A) Annotated UMAP of the pupal integrated neuron data. GRN = gustatory receptor neuron; MSNCB = Mechanosensory neuron in chemosensory bristle.

(B) Annotated UMAP of male neuronal cells subsetted from the Fly Cell Atlas single nuclei RNA-seq leg dataset (Li *et al.*, 2022).

(C-V) UMAPs of the pupal and adult neurons overlaid with the expression of genes identified as being enriched in the putative chordotonal clusters in the adult data. Note that of these, only *CG3339, CG9650, CG43689,* and *yuri* were present in the pupal neuron dataset. (C-K) *CG9492, CG10185, CG13133, CG9650, nan, CG14342, Hsc70-2,* and *CG43149* were found across all putative chordotonal organ cells and largely or entirely absent from the pupal ta1 neuron data. (L) *Dhc93AB* was present in all chordotonal clusters except for the *bab1*$^{+}$ population. (M) *NPFR* was present in all chordotonal clusters except for the *erm*$^{+}$ population. (N-P) The population enriched for the expression of the transcription factor *erm* showed specific expression of *CG14693, Dhc98D,* and *CG13408*. (Q-R) Among chordotonal populations, the transcription factor *yuri* was largely restricted to the *erm*$^{+}$ cluster, but also showed expression in GRNs. (S-V) The *bab1*$^{+}$ population showed enriched expression of *CG3339* and *CG43689*, although both genes showed spatially restricted expression in a subset of cells in the major chordotonal cluster.

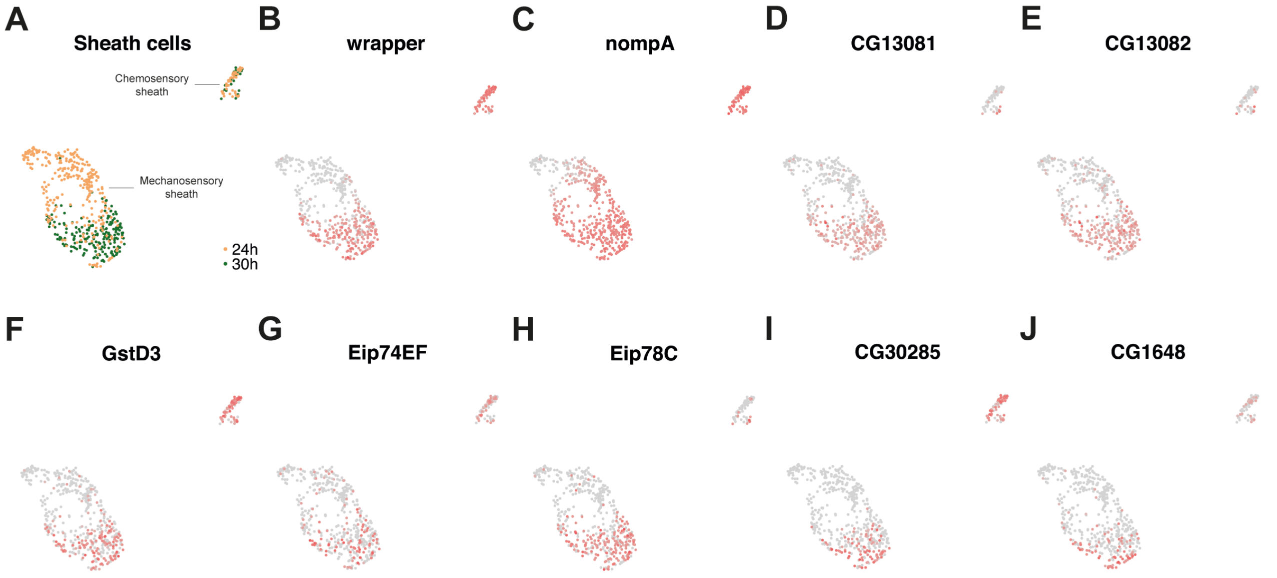


**Figure 8–figure supplement 1. Heterochronic differences between mechanosensory and chemosensory sheath cells.**

(A) A subset of the sensory support cell UMAP shown in Figure 8B showing only the mechanosensory and chemosensory sheath populations. Cells are coloured in relation to their dataset of origin and therefore, by extension, the timepoint after puparium formation at which they were collected.

(B-J) The UMAP shown in (A) overlaid with the expression of genes in the mechanosensory sheath cluster that we identified as being significantly upregulated in cells from the 30h dataset compared to those from the 24h dataset.

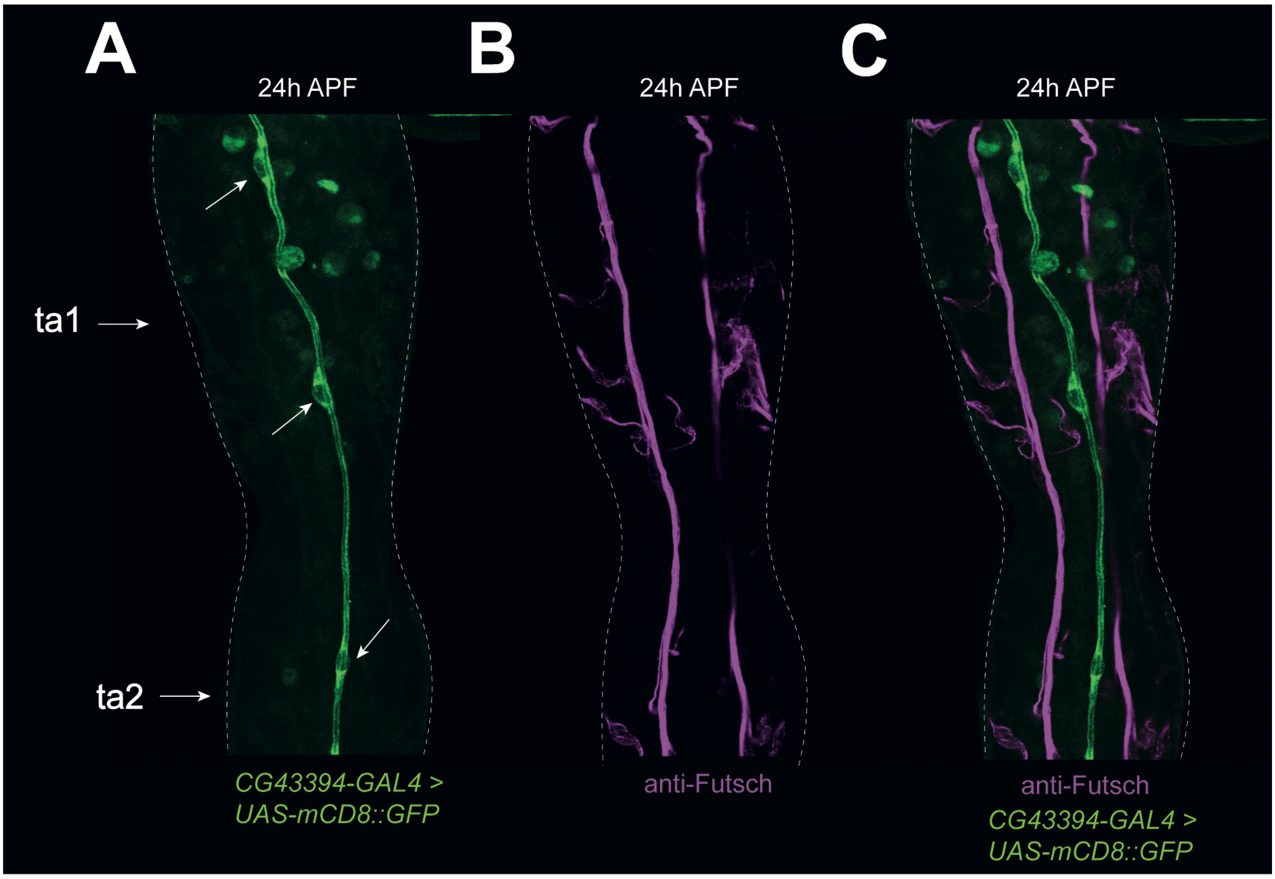


**Figure 8–figure supplement 2. Visualizing the expression of *CG43394*.**
Confocal images of a 24h APF male first tarsal segment from *CG43394-GAL4 > UAS-mCD8::GFP* counter-stained with the neuronal marker anti-Futsch. Visible cell bodies are highlighted with arrows in (A). *CG43394* was one of the top markers for a cluster we identified as socket cells from chemosensory bristles. This annotation was based on the strong transcriptomic overlap between the cluster and the other socket populations, including the shared expression of the known socket transcription factors *Su(H)*$^{+}$ and *Sox15*$^{+}$ (Figure 8G). However, the *CG43394-GAL4 > UAS-mCD8::GFP* staining does not appear to correspond to chemosensory sockets, but rather to a trachea-like channel running through the middle of the tarsal segments. While we cannot exclude the possibility that the cluster we annotated as chemosensory sockets in fact corresponds to these cells, we have reason to doubt that it does. In addition to the socket-like transcriptomic profile of the cells, it seems unlikely that a cell type with ~2 cells in the first tarsal segment would generate a cluster of equivalent size to the *CG43394*$^{+}$ population. For comparison, this cluster included 69 cells, compared to the 84, 74, and 76 in each of the putative sex comb sockets and shafts and chemosensory sheaths, respectively (~11 of each are found in a single ta1).
